# A convergent malignant phenotype in B-cell acute lymphoblastic leukemia involving the splicing factor SRRM1

**DOI:** 10.1101/2021.12.13.472370

**Authors:** Adria Closa, Marina Reixachs-Solé, Antonio C. Fuentes-Fayos, Katharina E. Hayer, Juan Luis Melero, Fabienne R. S. Adriaanse, Romy S. Bos, Manuel Torres-Diz, Stephen Hunger, Kathryn G. Roberts, Charles Mullighan, Ronald W. Stam, Andrei Thomas-Tikhonenko, Justo P. Castaño, Raúl M. Luque, Eduardo Eyras

## Abstract

A significant proportion of infant B-cell acute lymphoblastic leukemia (B-ALL) patients remains with a dismal prognosis due to yet undetermined mechanisms. We performed a comprehensive multicohort analysis of gene expression, gene fusions, and RNA splicing alterations to uncover molecular signatures potentially linked to the observed poor outcome. We identified 87 fusions with significant allele frequency across patients and shared functional impacts, suggesting common mechanisms across fusions. We further identified a gene expression signature that predicts high risk independently of the gene fusion background and includes the upregulation of the splicing factor *SRRM1*. Experiments in B-ALL cell lines provided further evidence for the role of SRRM1 on cell survival, proliferation, and invasion. Supplementary analysis revealed that SRRM1 potentially modulates splicing events associated with poor outcomes through protein-protein interactions with other splicing factors. Our findings reveal a potential convergent mechanism of aberrant RNA processing that sustains a malignant phenotype independently of the underlying gene fusion, and that could potentially complement current clinical strategies in infant B-ALL.

## Introduction

B-cell acute lymphoblastic leukemia (B-ALL) is the most common form of childhood cancer worldwide and one of the leading causes of cancer-related deaths in children (1). B-ALL presents a general lack of mutations in gene drivers that could be therapeutically targeted and are common in solid tumors and other leukemias (2). In contrast, B-ALL presents frequent chromosomal translocations that lead to the expression of gene fusions that are associated with marked differences in response to chemotherapy and survival. For instance, the frequent *ETV6-RUNX1* and *TCF3-PBX1* fusions have been associated with better prognosis (3), whereas *BCR-ABL1*, the highly frequent rearrangements of the gene *KMT2A* (*KMT2A-r*), and the less frequent *TCF3-HLF* are associated with poor prognosis (3, 4). Other rarer, less studied fusions remain with an uncertain prognosis.

The prevalence of gene fusions has spurred multiple efforts to identify treatments that target them or their downstream effectors, albeit with limited success (5). In particular, many fusions involve transcription factors, which are difficult to target directly (6). Despite these challenges, combination chemotherapy and recent advances in Chimeric Antigen Receptor (CAR) T-cell therapy have led to a 90% increase in the 5-year survival rate in children younger than 15 years and a 75% increase for adolescents (15-19 years). However, infant B-ALL remains with a bad prognosis, especially for the *KMT2A-r* cases, which occur in about 80% of the infant patients during embryonic/fetal hematopoiesis (7). A total of 135 different translocation partners have been identified for *KMT2A*, with the most frequent ones being members of transcriptional elongation complexes, accounting for 90% of all *KMT2A-r* cases (8). Furthermore, many of these fusions may occur at any age and are also frequent in acute myeloid leukemia (AML) (8). Genome-sequencing studies of *KMT2A-r* B-ALL have confirmed a very low frequency of somatic mutations, suggesting that *KMT2A-r* may not require additional alterations to induce transformation (9, 10). However, B-ALL cannot be recapitulated in pre-clinical models that only integrate the fusion, suggesting that additional alterations are necessary for leukemogenesis (11, 12).

The rarity of many of the fusions in B-ALL and their apparent links to functionally distinct pathways complicate their interpretation and the identification of effective therapies. On the other hand, there is increasing evidence for convergent molecular signatures in B-ALL that indicate similar disease progression patterns and common therapeutic vulnerabilities, despite presenting different genetic alterations. For instance, a BCR-ABL1-like B-ALL subtype was described that shows a gene expression profile and a therapeutic vulnerability similar to *BCR-ABL1* patients, despite not presenting the *BCR-ABL1* fusion (13). For *KMT2A-r* fusions, the functional impacts of the different fusion gene pairs have been linked to common downstream mechanisms of chromatin and transcription dysregulation (12). The occurrence of *KMT2A-r* across different ages and lineages and their general association with poor prognosis suggests that a convergent phenotype might be potentially identified in B-ALL associated with poor prognosis. We hypothesized that this phenotype might be captured in the transcriptome and present in high-risk B-ALL tumors independently of the fusion background.

Here we describe a comprehensive multi-cohort, infant and child-focused characterization of B-ALL transcriptomics signatures of high risk. We identified a predictive gene expression signature of high risk independent of the fusion background. This signature was mainly composed of ribosome biogenesis and RNA processing regulators, including the splicing factor *SRRM1*. Experiments in B-ALL cell lines provided evidence for the functional role of SRRM1 in cell survival, proliferation, and invasion. Furthermore, we found an alternative splicing program associated with the high-risk signature potentially mediated by splicing factors interacting with SRRM1. Our results provide a new layer of molecular variation that has remained undetected so far and that represents a potential source of novel prognostic markers and therapeutic strategies in B-ALL.

## Results

### RNA sequencing identifies novel gene fusions in B-ALL

We collected RNA sequencing (RNA-seq) from 428 patient samples obtained at diagnosis from five different B-ALL studies (14–18) (Supp. Table 2 and Data file 1). Patients’ age distribution peaked around one year old, with most cases being classified as infants or young children (Fig. 1a). We applied a comprehensive pipeline to study gene fusions, expression, and RNA splicing (Supp. Fig. 1) (Methods). To identify fusions likely to be associated with the B-ALL phenotype, we removed fusion candidates that we had also detected in non-cancer tissues and normal hematopoietic cells, as well as potential artifacts. We kept all fusion candidates reported in the clinical data of the studied cohorts or involving genes with acute leukemia mutations in COSMIC (19). Additionally, only candidates appearing in 5 or more samples and across different projects were considered. Starting from 1,825 unique candidate fusions, these filters resulted in 158 unique high-confidence fusions (Supp. Fig. 2) (see Methods for details).

**Figure 1.**
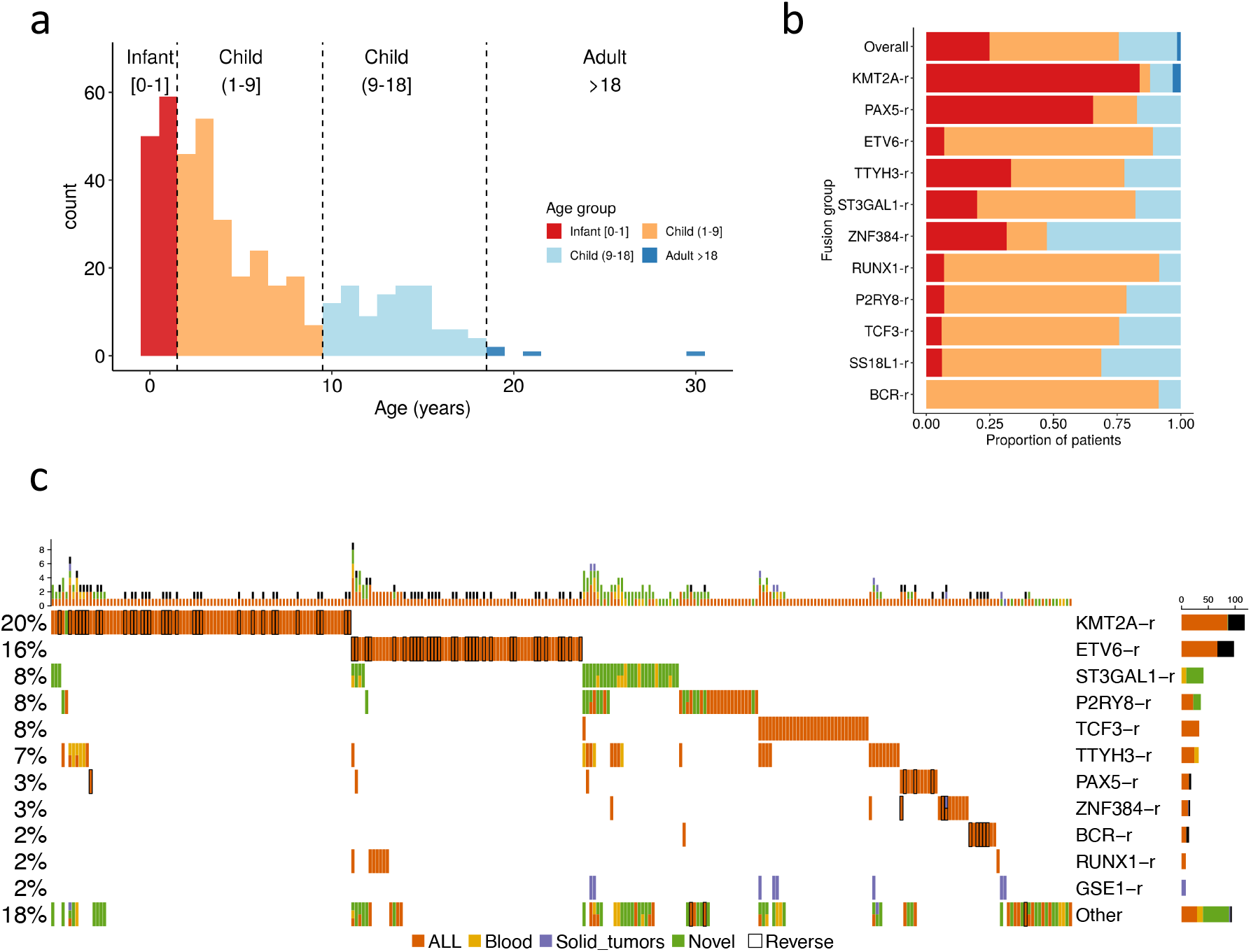
Multicohort identification of gene fusions in B-ALL. **(a)** Age distribution of the B-ALL patients studied. **(b)** The proportion of the gene fusion groups in each age group from (a). **(c)** Fusion oncoprint. The plot shows the most frequent fusions (rows) detected in patients (columns). Fusions are grouped by the most frequent gene in the fusion gene pairs. The bar plot above shows the number of fusions detected per patient. The specific gene fusion pairs in each patient are given in (Supp. Data file 2). A black border line indicates that the reverse fusion was identified in that patient. Cell colors indicate whether the fusions were detected before in B-ALL (orange), in a blood cancer (yellow), in a solid tumor (purple), or whether it was not reported in any cancer before (green). A patient with two different fusions detected in the same gene is depicted with two colors, and they were only kept if they had been previously reported in a tumor and occurred in at least five patients.

To prioritize the relevance of the identified fusions, we further calculated a fusion allele frequency (FAF) (20), defined as the proportion of the gene expression corresponding to a fusion averaged over the two genes participating in the fusion, and which represents a proxy for the fusion clonality (Supp. Fig. 3). We then considered only those fusions with a median FAF of at least 0.1 and having none of its genes with a median individual FAF below 0.1. These analyses resulted in our final list of 87 different fusions (Data file 2) (See methods for details).

Using these fusions, all 5 analyzed cohorts presented a similar distribution of fusions per patient (Supp. Fig. 4). Our analyses of the RNA-seq data recovered 81.35% of the most frequent fusions detected in the same samples with independent experimental methods (Supp. Figs. 5a and 5b). Although most of the identified fusions had been observed previously in ALL, we also identified fusions that had been reported before in other blood cancers or in solid tumors, as well as novel fusions (Supp Fig. 5c). Moreover, we identified known B-ALL fusions in 43 patients that did not have any fusion annotated in the published clinical information and determined the fusion partner in 7 cases that were only annotated as *KMT2A-r* (Data file 2).

We found that some of the identified fusions are overrepresented in specific age groups (Fig. 1b). We recovered the known enrichment of *KMT2A-r* in infant cases and *ETV6-r* in children and young adults (1-18 years). Fusion groups such as *BCR-r* and *P2RY8-r* presented a bimodal or extended age distribution, including infant and child. Similarly, *PAX5-r* appeared in infants and at the upper extreme of childhood cases (Supp. Fig. 6). Despite these associations, age distributions and fusion frequencies may not represent the actual distribution in the population of leukemia patients, as there may have been sample collection biases in each cohort.

Overall, high-confidence fusions were detected for 70% of the samples at diagnosis. Grouping the fusions by the most frequent gene in the fusion pairs, we could identify six major fusion groups: *KMT2A-r* (20%), *ETV6-r* (16%), *ST3GAL1-r* (8%), *P2RY8-r* (8%), *TCF3-r* (8%), and *TTYH3-r* (7%) (Fig. 1c). In the cohorts analyzed, we also found *PAX5-r* (3%), *ZNF384-r* (3%), *BRC-r* (2%), *RUNX1-r* (2%), and *GSE1-r* (2%), as well as a group of low-frequency fusions that appeared in 18% of the patients. The calculated fusions in diagnostic samples showed a clear pattern of mutual exclusions and a frequent co-occurrence of the inverse fusion for *KMT2A-r, ETV6-r, PAX5-r*, and *BCR-r*, but not for other fusions (Fig. 1c). Apart from known fusions in B-ALL and hematological malignancies we described fusions previously observed in solid tumors, *GSE1-SLC7A5* and *CBFA2T3-PIEZO1*. Among the fusions that have not been previously described in cancer, the majority can be attributed to new partners of already known fusion genes in B-ALL or other hematological malignancies such as *ST3GAL1* or *P2RY8* (Fig. 1c) (Data file 2).

### Different gene fusions impact similar functional pathways

We calculated the breakpoints for each of the fusions detected from RNA-seq reads, either as the exon position where the fusion breakpoint was found or using the boundaries of the exons flanking the intron where the breakpoint was assumed to fall. Fusion genes presented multiple breakpoints (Fig. 2a) (Supp. Fig. 7). Moreover, these breakpoints appeared in positions that potentially disrupted the protein domain content. This raised the question of whether different breakpoints in the same or different fusion genes may lead to similar functional impacts. To determine this, we calculated the domains that are maintained or lost in each fusion gene according to the identified breakpoints. In terms of the specific domains kept or lost in a fusion, we observed little overlap between fusions (Supp. Fig. 8). However, when we grouped the protein domains according to their functional ontologies, multiple similarities appeared, such as the loss of DNA-binding domains in *KMT2A-r* and *ETV6-r* and the loss of signal transduction domains in *P2RY8-r* and *BCR-r* (Fig. 2b).

**Figure 2.**
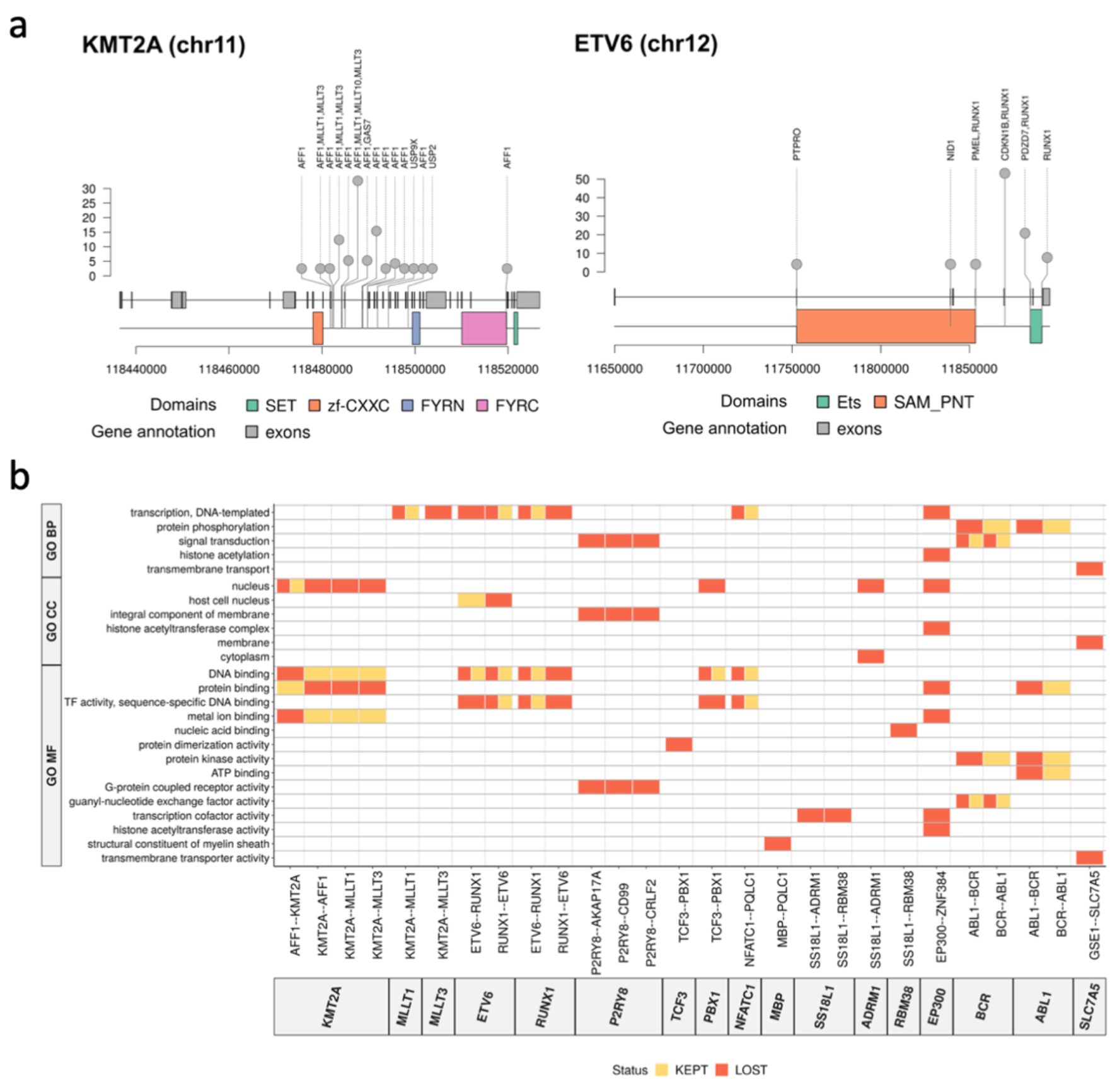
Functional impact assessment of gene fusions. **(a)** Identified breakpoints in the fusions involving *KMT2A* (left panel) and *ETV6* (right panel). Along the genomic locus (x-axis), we indicate the detected breakpoint’s position, defined by an exonic position if the breakpoint occurred inside an exon, or by the last base of the last exon included in the fusion otherwise. The plot also shows the span of the functional domains in genomic coordinates. **(b)** Domains kept (orange) and lost (red) through gene fusions (x-axis) labeled according to gene ontologies (GO) (y-axis): Biological Process (BP), Cellular Compartment (CC), and Molecular Function (MF). The specific functional domains kept or lost in these fusions are given in Supplementary Figure 8.

### Common patterns of gene expression across diverse gene fusion backgrounds

To further characterize the cellular processes associated with the identified fusion groups, we calculated the differential expression patterns among patient groups. Despite the heterogeneity of samples used in the comparison, there were many significant expression changes associated with *KMT2A-r, ETV6-r, ST3GAL1-r, ZNF384-r*, and *TCF3-r* in comparison with the other fusion groups and with patients without fusions (Supp. Fig. 9) (Data file 3). These patterns included the HOXA overexpression characteristic of *KMT2A-r* patients (Supp. Fig. 10) (16). Interestingly, the differentially expressed genes associated with each fusion group did not generally overlap (Supp. Fig. 11), except for the *KMT2A-r* and *ETV6-r* groups, which showed opposite expression patterns (Supp. Fig. 12). This suggested that the functional alterations specific to *KMT2A-r* are reversed in *ETV6-r* tumors.

To investigate this possibility, we studied the pathways enriched or depleted in each fusion group. Each group tended to cluster independently, except for *ST3GAL1-r, PQLC1-r*, and *TTYH3-r*, which presented similar pathway enrichments, and *P2RY8-r, RUNX1-r*, and *PAX5-r*, which were similar to the cases with no fusions (Supp. Fig. 13). Moreover, genes overexpressed in KMT2A-r patients were strongly enriched in MYC targets, ribosome biogenesis, and RNA processing, including splicing and translation regulation (Supp. Fig. 13). Interestingly, MYC targets were upregulated in *KMT2A-r* patients but depleted in *ETV6-r* (Supp. Fig. 13), and transforming growth factor beta (TGF-b) signaling, which antagonizes MYC (21), was depleted in *KMT2A-r* patients. Furthermore, *MYC* expression was higher in *KMT2A-r* patients relative to the other patient groups and normal fetal-liver B-cells (Supp. Fig. 14), whereas *ETV6-r* patients showed *MYC* expression below normal fetal liver B-cells (Supp. Fig. 14). This suggested a gene expression pattern linked to *MYC*, in association with *KMT2A-r*, and reversed in *ETV6-r*.

### A gene expression signature of high-risk B-ALL independent of the gene fusion background

B-ALL cases with *KMT2A-r* are generally considered to have a poor prognosis, whereas *ETV6-r* are deemed to be of good prognosis (3). We thus reasoned that the observed opposite expression patterns might be linked to these risk phenotypes. To investigate this possibility, we calculated the genes differentially expressed between patients harboring only *KMT2A-r* or *ETV6-r*, without any other detected fusion (87 *KMT2A-r* and 68 *ETV6-r*). All RNA-seq used was obtained at the diagnostic stage before treatment. Gene set enrichment analysis on the 1405 genes with significant differential expression (FDR < 0.01 and |logFC| > 1) (Fig. 3a) (Data file 4) revealed two main pathways, one associated with immune response and another related to MYC, translation regulation, and ribosome biogenesis (Fig. 3b). As RNA processing is tightly controlled during normal hematopoiesis and is commonly dysregulated in hematological malignancies (22–24), we hypothesized that genes related to these pathways may separate patients according to their risk of relapse. Feature selection was conducted to extract a list of 39 genes from the target pathways showing overexpression in *KMT2A*-r. *RBM24* and *RNU6-1* were discarded from these genes as they showed high variability across the studied cohorts, resulting in a final signature of 37 genes (Fig 3c) (Supp. Fig. 15).

**Figure 3.**
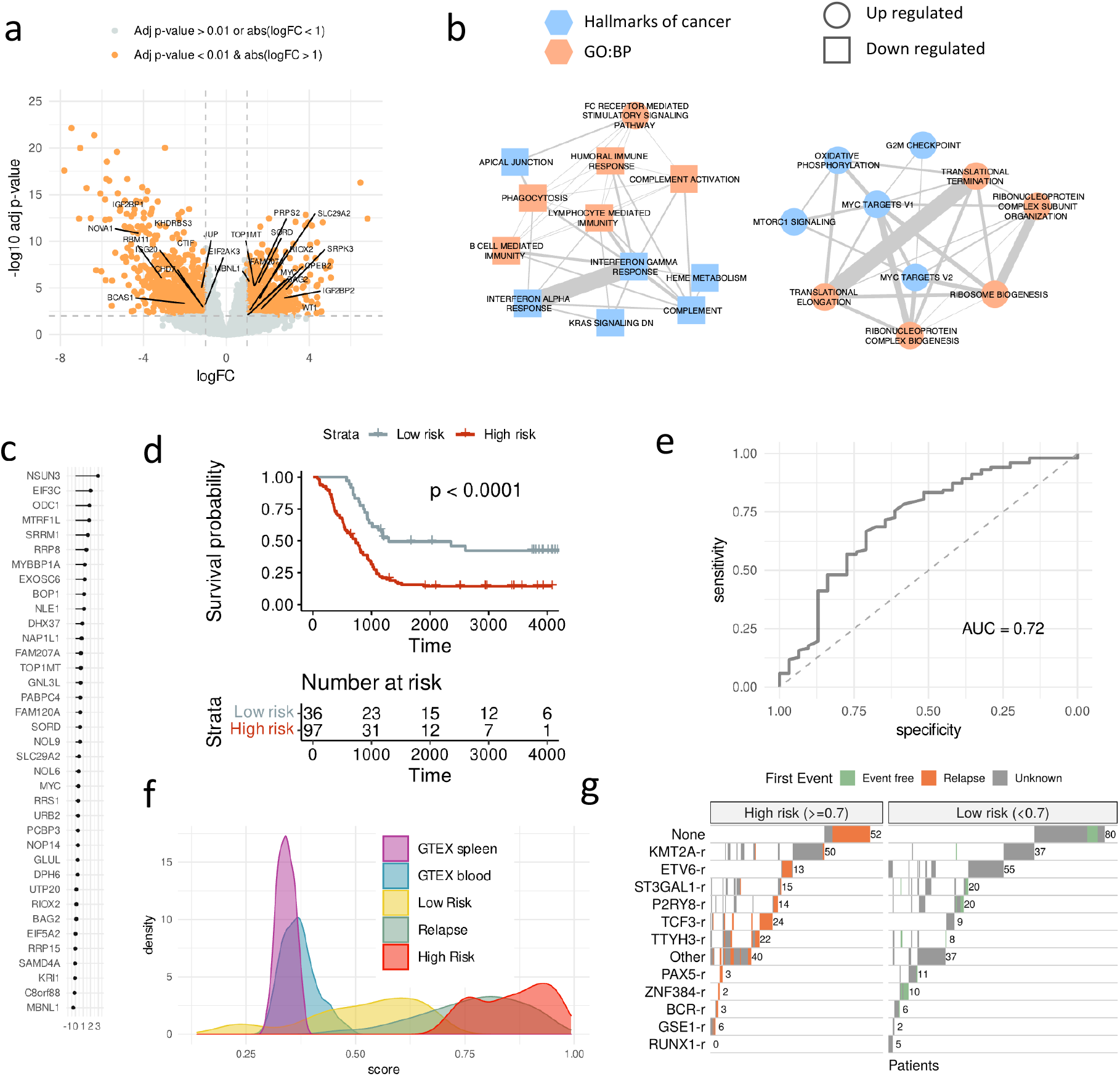
Gene expression signature of high risk. (**a**) Differential gene expression between a subset of patients with only *KMT2A-r* or only *ETV6-r*. The volcano plot shows for each gene the log2(fold-change) (x-axis) and the -log10(corrected p-value) (y-axis). We indicate the genes involved in RNA processing, RNA translation, and Ribosome biogenesis. **(b)** Enriched Cancer Hallmarks and Biological Process Gene Ontologies (GO:BP) of the genes significantly up or down-regulated in the comparison in (a). **(c)** Ranking of the 37 genes of the predictive model of high risk according to their relevance in the leave-one-out test (x-axis). Relevance is defined as the median of the accuracy per gene. **(d)** Kaplan-Meyer plot of the patients separated as high risk (red) (risk score >= 0.7) or low risk (grey) (risk score < 0.7) in a leave-one-out test. **(e)** Average receiving operating characteristic (ROC) curve and area under the ROC curve (AUC) from the leave-one-out test for classifying patients into low and high risk. The ROC curve shows the specificity (x-axis) and sensitivity (y-axis) for the entire range of possible model score threshold values. Sensitivity is defined as the proportion of high-risk cases that are correctly predicted. In contrast, specificity is defined as the proportion of low-risk cases correctly predicted as low risk. **(f)** Risk score distribution for various sample groups: diagnostic samples predicted as high risk, diagnostic samples predicted as low risk, samples obtained at relapse, and blood and spleen samples from GTEX. **(g)** Classification of all the B-ALL diagnostic samples from each fusion subgroup (y-axis) into high (left) or low (right) risk according to our risk score. Patients with a clinical record indicating that they had relapsed are indicated in orange, whereas patients annotated with no relapse are indicated in green (Event free). Patients with no clinical information are indicated in grey (unknown).

We trained and tested a predictive model for risk using the 133 patients from TARGET, a high-risk cohort with clinical follow-up information (Data file 1), using as endpoint the event-free survival and as the prognostic event the first relapse. Using a random forest leave-one-out cross-validation combined with a cox-regression, we obtained a significant separation of patients according to the risk of relapse (log-rank test p-value < 0.0001) (Fig. 3d) and overall accuracy of 0.72, measured as the area under the receiver operating characteristic (ROC) curve (AUC) (Fig. 3e). Importantly, even though the genes in our model were selected from the comparison of *KMT2A-r* and *ETV6-r* patients, only 11% of the 133 patients had KMT2A-r or ETV-6 fusions. Based on the observed values of sensitivity, specificity, and accuracy (Supp. Fig. 16a), we chose the decision boundary at score 0.7 (>=0.7 for high risk and <0.7 for low risk). However, the same accuracy values were maintained with score thresholds between 0.5 and 0.7 (Supp Fig. 16a).

To further evaluate our predictive signature, we trained a single model with all the 133 TARGET samples (Supp. Fig. 16b). We applied this model to normal blood and spleen samples from GTEX, all other B-ALL samples, and to 82 additional samples obtained at relapse, none of which were included in any of the analyses above. Remarkably, samples obtained at relapse showed a distribution similar to the high-risk samples and higher than the normal samples (Fig. 3f). Furthermore, this model separated TARGET patients with and without relapse and detected other high-risk patients from the other cohorts (Fig. 3g) (Supp. Fig. 16c). Our predictions showed that patients with fusions *KMT2A-AFF1, AFF1-KMT2A*, and *TCF3-PBX1* were more frequently in the high-risk group. In contrast, patients with *ETV6-RUNX1, RUNX1-ETV6*, and *ABL1-BCR* were more often classified as low risk, including the ability to stratify patients with higher risk inside every group of fusion (Fig. 3g) (Supp. Fig. 16c) (Data file 5).

We further applied our model to an independent cohort of 188 B-ALL patients (25). Since the gene expression values for this dataset were only available in terms of regularized log (rlog) values, we rebuilt our model using the rlog expression values calculated from the TARGET samples. Using a leave-one-out cross-validation, this model maintained the same classification power as the previous model built with logCPM values (Supp. Fig. 17a), and the scores given by both models showed a high correlation (R=1, p-value<2.2e-16) (Supp. Fig. 17b). To test our predictions, we used the clinical classification of patients published by the independent study into three different risk groups (high, standard, and low) (25). Our model separated these three groups into significantly different score distributions, with the high-risk group showing the highest scores and the low-risk group showing the lowest signature scores (Supp. Fig. 17c). These analyses provide strong support for the ability of our model to predict the potential for relapse in B-ALL independently of the gene-fusion background. To make this model readily available to evaluate the risk on new sets of patients from expression data and to explore the features of the TARGET cohort, we integrated the predictive model into an interactive web resource available at https://github.com/comprna/risk_model_app.

### SRRM1 as a candidate driver of progression and poor prognosis in B-ALL

One of the genes with the highest predictive value in our risk-prediction signature was *SRRM1*, which showed the highest correlation with the risk score (Pearson r= 0.51 p-value = 8.18e-30) (Data file 5) and has been associated before with poor prognosis in prostate cancer (26). Confirming this predictive power, patient samples classified as high-risk and relapse samples showed higher *SRRM1* expression than low-risk samples and normal samples from spleen and blood (Fig. 4a). To further understand the functional transformations associated with *SRRM1* expression, we analyzed multiple samples from progenitor and mature B-cells. *SRRM1* was significantly downregulated in mature B-cells compared with B-cells progenitors (Fig. 4b). Consistent with this, the risk score was significantly higher (0.50-0.73) in undifferentiated B-cells compared to differentiated B-cells (0.38-0.46) (p-value < 0.001) (Supp. Fig. 18) (Data file 6). These results suggest a relevant role for *SRRM1* driving a highly proliferative phenotype.

**Figure 4.**
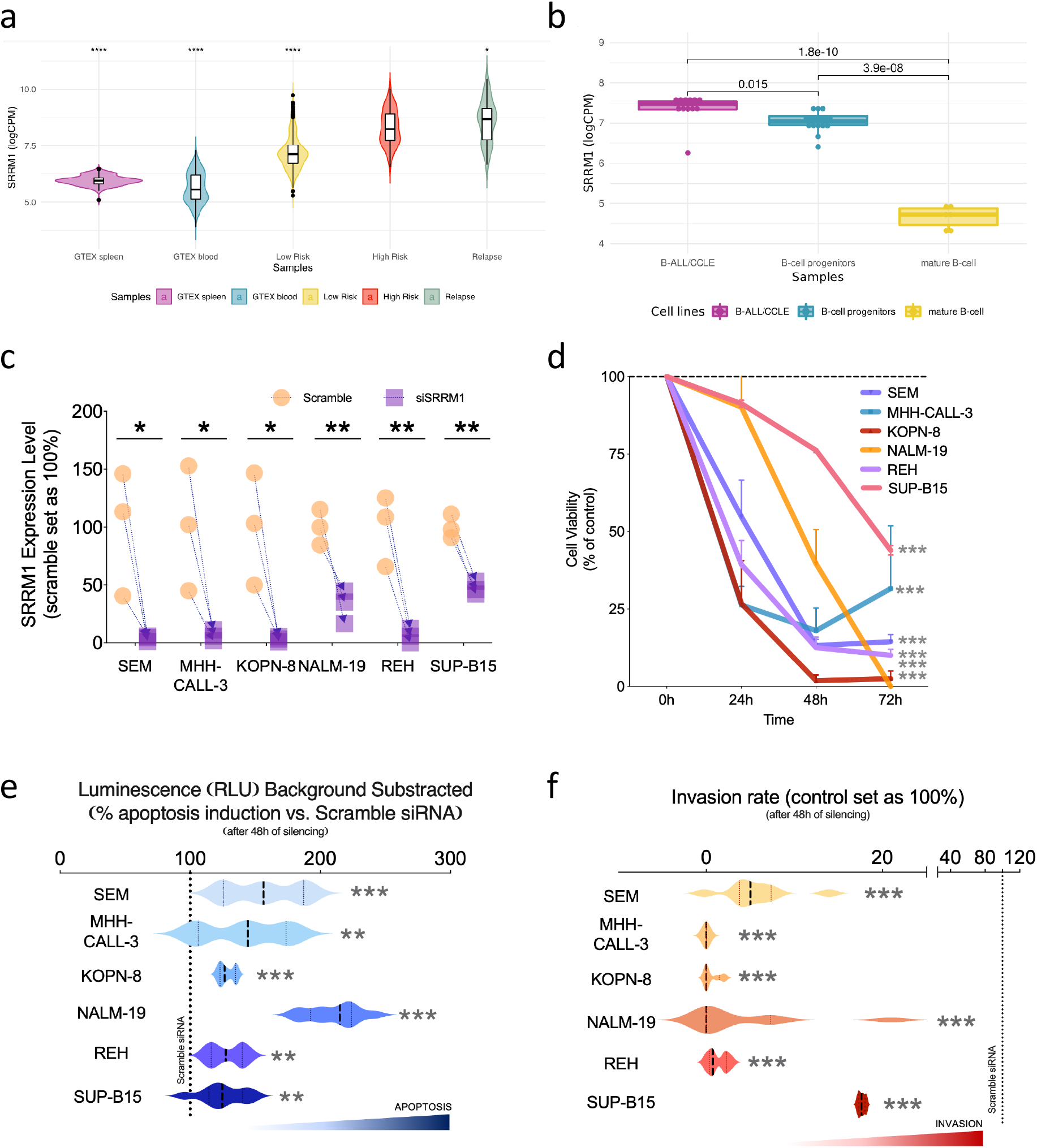
SRRM1 as a potential major driver of high-risk B-ALL. **(a)** Distribution of expression values for SRRM1 in log10 counts per million (logCPM) units (y-axis) in various sample groups: B-ALL diagnostic samples predicted to be high risk (score > 0.7), B-ALL diagnostic samples predicted to be low risk (score < 0.7), B-ALL relapse samples, and blood and spleen samples from GTEX (t-test applied to compare all the groups versus the high-\isk group, **** -> p < 2.2e-16, * -> p = 0.023). **(b)** Normalized log2CPM expression of *SRRM1* in B-ALL cell lines from the cancer cell line encyclopedia (CCLE), B-cell progenitors, and a set of GM12878 biological replicates. P-values were obtained from a t-test mean comparison. **(c)** Validation of *SRRM1* silencing (y-axis) in cells (x-axis) by qPCR (n=3). **(d)** Proliferation rate (y-axis) in response to *SRRM1* silencing in leukemia cell lines at different time points (x-axis) (n=3 per time point). **(e)** Apoptosis rate (x-axis) in response to *SRRM1* silencing in leukemia cell lines (y-axis) (n=3). **(f)** Invasion rate (x-axis) in response to *SRRM1* silencing in human leukemia cell lines (y-axis). The dotted lines represent the control condition (scramble transfected) 100%. Data is presented using the mean ± standard error of the mean. Significant differences from control conditions were indicated as * for P-value < 0.05, ** for P-value < 0.01, *** for P-value <0.001.

*SRRM1* has been described as an essential gene, but a knockdown is known to produce a reduction of the cell viability without killing the cell (Supp. Fig. 19), opening the door to *SRRM1* expression modulation as a potential strategy to reduce the aggressive phenotype of leukemia cells. To test this strategy, we determined proliferation, apoptosis, and invasion rate, after silencing *SRRM1* in six human B-ALL leukemia cell line models bearing distinct functional and phenotypic features: SEM, MHH-CALL-3, KOPN-8, NALM-19, REH, and SUP-B15. The silencing of *SRRM1* expression was successful in all cell models (Fig. 4c) and resulted in a significant decrease in proliferation rate in a time-dependent and cell line-dependent manner (Fig. 4d). Furthermore, using the capase3/7 assay revealed that *SRRM1* silencing significantly induced apoptosis in all human leukemia cells (Fig. 4e).

Moreover, using a trans-well assay to evaluate the invasion capacity revealed that *SRRM1* silencing could potentially impair the capacity of these cells to invade surrounding tissues in all cell lines tested (Fig. 4f). *SRRM1* expression analysis revealed that the cell lines had different basal expression patterns (from high to low levels: KOPN-8>SEM>REH>MHH-CALL-3>NALM-19>SUP-B15) and presented differences in the basal proliferation (Supp. Fig. 20). Furthermore, the proliferation rate with the siRNA at 48h was inversely correlated with the *SRRM1* basal expression and the silencing effectiveness (Supp. Fig. 20), supporting a tight association of *SRRM1* expression with this essential cellular function.

### A candidate SRRM1-dependent splicing program associated with high-risk B-ALL

SRRM1 is a Serine-Arginine rich factor that is associated with splicing complexes and affects splicing through interactions with SR proteins (27) (Fig. 5a). We thus decided to study the alternative splicing events potentially linked with SRRM1 and their association with poor prognosis. Analysis of differential RNA processing events between patients predicted as high and low risk identified a total of 422 events, with 342 of them affecting coding transcripts and showing enrichment of alternative first (AF) and skipping exons (SE) (Fig. 5b) (Data file 7). Those 342 events occurred in 271 genes, none of which were differentially expressed between the same conditions. expressed genes. Moreover, out of the 342 events with differential splicing, only 27 (7.8%) showed a correlation higher than 0.5 (Pearson R) between their PSI and the expression level of the host gene across the samples. This indicates that the splicing modulation identified is largely independent of the expression changes. Moreover, these significant events separated high and low risk patients independently of the cohort (Fig. 5b) (Supp. Fig. 21. Furthermore, events that differentiate the high and low-risk patients, as well as the genes where they occur, had a small overlap with those that differentiate between KMT2A-r and ETV6-r patients (Supp. Fig. 22) (Data file 8). This suggests that similar to the expression signature described above, there may be a splicing signature associated with risk that is independent of the fusion background.

**Figure 5.**
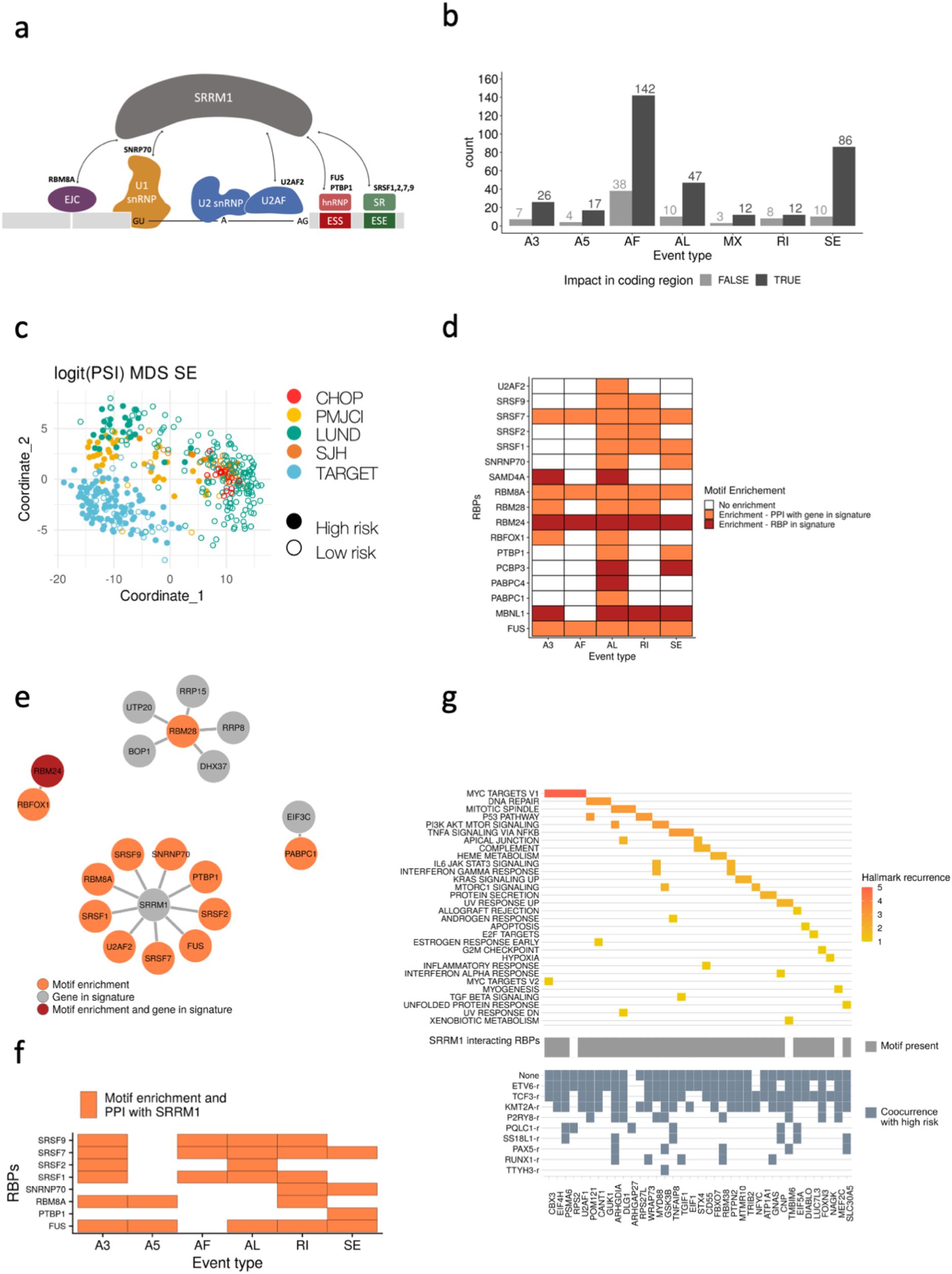
Splicing signature associated with high-risk. **(a)** SRRM1 is a known interactor of multiple splicing factors and RNA binding proteins (RBPs). **(b)** Identified significant splicing changes between high and low-risk patients. The counts are separated according to whether the splicing change impacts a coding region (black) or not (grey), and by event type: alternative 3’ (A3) and 5’ (A5) splice site, alternative first (AF), and alternative last (AL) exon, mutually exclusive exons (MX), retained intron (RI) and skipping exon (SE). **(c)** Multidimensional scaling (MDS) plot of the analyzed samples using only the significant SE events. The color indicates the patient cohort, and the full (empty) circle indicates if the sample is predicted to be of high (low) risk. Other event types are shown in Supplementary Fig. 22. **(d)** Enrichment (z-score > 1.5) of binding motifs for RNA binding proteins (RBP) (y-axis) in each event type (x-axis). The plot indicates whether the RBP is part of the high-risk signature (red) or whether it has a reported protein-protein interaction (PPI) with a gene from the signature (orange). Event types not showing on the figure means that there are no binding motifs of RBP associated with the events. **(e)** Protein-protein interaction networks for the RBPs in the predictive signature and/or with binding motifs enriched in the events differentially spliced between high and low-risk patients. The color indicates whether the RBP motif was enriched (orange), whether the RBP was part of the high-risk signature (gray), or both (red). **(f)** RBPs and splicing factors with associated motifs enriched in events differentially spliced between mature B-cells and B-cell progenitors. **(g)** Upper panel shows cancer hallmarks associated with genes with differentially spliced events that co-occur with high risk in at least one fusion group, colored by its recurrence. In grey, the middle panel indicates the presence of binding motifs for any of the RBPs that interact with SRRM1. Lower panel highlights in blue the fusion groups in which the differentially spliced events co-occur with high risk.

To evaluate the SFs/RBPs potentially associated with these RNA processing changes, we performed a motif enrichment analysis. Specifically, we identified motifs for RBPs that were part of our high-risk signature: *SAMD4A* in A3 and AL events, *RBM24* in all the different types of events, *PCBP3* in AL and SE events, *PABPC4* in AL, and *MBNL1* in A3, AL, RI, and SE events. We also recovered motifs for other RBPs, including the SR protein genes *SRSF1, SRSF2, SRSF7*, and *SRSF9*. Interestingly, these and other SFs/RBPs with motifs enriched in events showed protein-protein interactions with our risk signature members (Fig. 5d) (Supp. Fig. 23). We identified an enrichment in motifs for RBM28, an interactor of 5 of the proteins encoded by genes of our high-risk signature: *RRP8, RRP15, UTP20, BOP1*, and *DHX37* (Fig. 5d). Similarly, we found an enrichment of motifs for *RBFOX1*, which interacts with *RBM24*, also part of the signature (Fig. 5e). Remarkably, there were 9 RBPs with motif enrichment in the events changing with the risk that had protein-protein interactions with the splicing factor SRRM1 (Fig. 5e). Moreover, SRRM1 expression correlated with the inclusion of events differentially spliced between high and low-risk patients, for events harboring binding motifs for RBPs with evidence of protein-protein interaction with SRRM1 (Supp. Fig.24). Such correlation was not always observed for RBPs with enriched motifs and interacting with SRRM1. Moreover, SRRM1 was not among the splicing factors with the highest number of potential interactors (Supp. Fig. 25). These analyses suggest that a high-risk phenotype may be associated with a change in the expression of multiple SFs and RBPs that impact RNA processing, with a potentially prominent role for *SRRM1*.

We observed before that *SRRM1* expression as well as our risk signature score were higher in B-cells progenitors compared with mature B-cells. We thus next tested whether SRRM1 could play a role in the splicing changes across B-cell differentiation, we calculated the differentially spliced events and performed motif enrichment between progenitors and mature B-cells. Importantly, differentially spliced events between progenitors and mature B-cells contained motifs for the same SRRM1-interacting RBPs and splicing factors that we observed for leukemia patients (Fig 5f). Additionally, the genes with differentially spliced events were enriched in the same pathways obtained in the comparison between high and low risk patients, which were different from the pathways enriched in genes with splicing differences between *KMT2A-r* and *ETV6-r* (Supp. Fig 26) (Data file 9). Interestingly, the expression level of the *KMT2A-AFF1* and *ETV6-RUNX1* fusions, the most abundantly observed in the *KMT2A-r* and *ETV6-r* groups, did not show any correlation with the risk score or with the *SRRM1* expression levels (Supp. Fig. 27).

To further investigate the possible mechanisms linking *SRRM1* with high risk, we calculated the subset of splicing events associated with risk that was also significantly associated with risk within each fusion group. Most of these events co-occurred with patients with *KMT2A-r, ETV6-r, TCF3-r*, as well as patients with no fusions (Fig. 5g, lower panel) (Supp. Fig. 28); and presented binding motifs for RBPs interacting with *SRRM1* (Fig. 5g, middle panel). Moreover, the genes harboring those events were significantly associated with cancer-related pathways, such as MYC targets, DNA repair, and the p53 pathway (Fig 5g, upper panel). One of these genes was *EIF4H*, a translation initiation factor that is key for translational control. Overexpression of *EIF4H* has been associated before with cell proliferation and increased chemoresistance in lung cancer (28). Our analyses indicated that one of the EIF4H isoforms (ENST00000265753.12) decreased expression in the low-risk group, while a second isoform (ENST000000353999.6) had stable expression across all patients (Supp. Fig. 29).

Taken together, our results provide suggestive evidence that there is a molecular signature of expression and splicing changes, possibly driven by *SRRM1* that is predictive of bad prognosis in B-ALL and that is independent of the fusion background.

## Discussion

In this study, we performed a multicohort age-agnostic analysis of B-ALL cases focused on the differences of risk outcome independent of the fusion background. We identified an expression pattern involving MYC targets, translational regulators, and splicing factors associated with an increased probability of relapse. This is consistent with previous results showing that translation is tightly controlled during normal hematopoiesis (22) and is commonly deregulated in cancers, including hematological malignancies (29, 30). Overexpression of *MYC* promotes expression of the translational machinery, increasing ribosome production and activity, leading to increased cell growth (23). Additionally, *MYC* overexpression plays a role in the dysregulation of the splicing machinery during lymphomagenesis (24). Our findings suggest that alterations in RNA processing could be involved in driving tumor progression and resistance to current therapies in B-ALL. This is consistent with recent work showing that aberrant splicing is directly implicated in the development of therapy resistance in B-ALL (31, 32)

We summarized our findings in a 37-gene signature that showed prognostic value on B-ALL patients samples. Our signature classified patients in high and low risk with high accuracy and independently of the fusion background. The same signature separated high-risk patients from normal blood and spleen samples, and B-ALL cell lines and B-cell progenitors from mature B-cells. These results provide evidence that high-risk B-ALL cases recapitulate a gene expression pattern independently of the gene fusion background. In further support of our findings, analysis of an independent cohort (25) showed that our signature provided a significant separation of patients according to their relapse (log-rank test p-value 0.00041) (Supp. Fig. 30a).

Our findings suggest that gene fusions operate mainly as initiating events, and an independent convergent mechanism defines high risk in B-ALL. This would agree with results using current murine and humanized models of *KMT2A-r* B-ALL (33) showing that they do not faithfully recapitulate the disease pathogenesis and suggesting that *KMT2A-r* alone is insufficient to sustain leukemia (34, 35). Our derived gene expression signature could complement current clinical assessment methods of B-ALL patients. Our signature would be beneficial as a strategy to identify patients with a high risk of relapse when no fusions are detected, or when the presented fusion is of unknown prognosis. Moreover, it is conceivable that a deeper understanding of these genes may provide further mechanistic insight into the common functional underpinnings underlying B-ALL and thus reveal potential novel therapeutic avenues. Our high-risk signature included several splicing factors (SFs) and genes encoding for RNA binding proteins (RBPs), such as *SRRM1, IGF2BP2*, and *MBNL1*, which suggested a pattern of differential RNA processing linked to high risk. MBNL1 is a protein involved in alternative splicing, predominantly regulates intron exclusion, and has been found consistently overexpressed in *KMT2A*-rearranged leukemia. Although inhibition of MBNL1 is linked with selective leukemic cell death, this effect seems to be *KMT2A*-r specific (36). Furthermore, the relative risk predictive power of *MBNL1* in our signature is low, possibly due to our cohorts having many patients with no *KMT2A* fusions, further supporting the specificity of MBNL1 to *KMT2A*-r cases.

Among the genes in our risk signature involved in RNA processing, *SRRM1* showed one of the strongest predictive powers and had the strongest correlation with the risk signature score. Samples at diagnosis from high-risk patients, before treatment, presented the highest *SRRM1* expression, similarly to samples obtained at relapse, and higher than normal blood/spleen samples. Moreover, *SRRM1* was highly expressed in B-ALL cell lines and B-cell progenitors compared to mature B-cells, further linking SRRM1 to a potential proliferative cellular state. Our analysis also showed that most of the splicing events changing between high and low-risk patients showed a correlation of their inclusion levels with SRRM1 expression and contained binding motifs for SFs/RBPs that have evidence of protein-protein interactions with SRRM1. Moreover, these events occurred in genes involved in cancer pathways. Importantly, analysis of an independent cohort (25) provided additional validation of the potential association of SRRM1 and its interactors with these splicing changes. Indeed, the splicing events changing between high and low-risk patients showed the same correlation patterns with SRRM1 and its RBP interactors in this new independent cohort (Supp. Fig. 30b).

Overexpression of *SRRM1* has been associated previously with poor prognosis in prostate cancer (26) and silencing of *SRRM1* was shown to reduce cell proliferation through a reduction of AKT phosphorylation levels and an increased expression of *PTEN*, a well-known tumor suppressor (26). Previous studies have also shown that *SRRM1* overexpression leads to the expression of a *CD44* isoform that acts as a RAS-signaling activator and induces metastatic potential in non-metastatic cells (38). Furthermore, *SRRM1* has been identified as part of a chromatin protein complex that drives B-cell differentiation (39). We showed that silencing of *SRRM1* in B-ALL cell models leads to a significant decrease in proliferation and invasion rates and a significant increase in apoptosis capacity. The effect in response to *SRRM1* silencing was different in all cell lines tested, especially in the decrease in proliferation rate. This appeared to be associated with the variable *SRRM1* expression found in these cell lines rather than with the basal proliferation rate of each cell model. This would indicate that tumors with higher *SRRM1* expression would depend more on *SRRM1* for proliferation. In contrast, tumors with lower *SRRM1* expression would rely on other proliferation mechanisms. Importantly, we analysed data from a recent pan-cancer protein map atlas based on 946 human cancer cell lines (37). This showed that SRRM1 presents the highest protein levels in hematological tumors (Supp. Fig. 31). This provides an additional layer of evidence for the potential role of SRRM1 in the progression of leukemia and, in particular, B-ALL.

In conclusion, we have presented a gene expression signature that predicts poor outcomes in samples at diagnosis independently of the fusion background. This signature is associated with *SRRM1* overexpression and with splicing changes potentially partly driven by SRRM1 interactions with other splicing factors. This leads us to propose that *SRRM1* overexpression may contribute to sustaining tumor malignancy and lead to poor prognosis in B-ALL. Furthermore, *SRRM1* could function as a novel prognostic marker of high-risk B-ALL, and its depletion could be used in combination with standard therapies to achieve more effective treatments in high-risk cases.

## Methods

### Data availability

All samples were obtained from various sources through controlled or public access. The series from St Jude Children’s Research Hospital (SJH) (EGAS00001000246) (18) and Lund University (LUND) (EGAS00001001795) (14) were downloaded from the European Genome-phenome Archive (EGA), and the corresponding clinical information was obtained from the associated publications. The series from Children’s Hospital of Philadelphia (CHOP) (GSE115656) was downloaded from Gene Expression Omnibus (GEO) (15). Samples from TARGET (Therapeutically Applicable Research to Generate Effective Treatments) were downloaded from the TARGET data portal at National Cancer institute (NIH) together with the associated clinical information, corresponding to dbGAP accessions phs000463 (ALL phase 1) and phs000464 (ALL phase 2). Data from patients from the Princess Maxima Center for Pediatric Oncology (PMJCI) from (16) was obtained from the authors. We also analyzed data from 12 B-ALL cell lines coming from the Cancer Cell Line Encyclopedia (CCLE) (40), from normal blood and spleen samples from The Genotype-Tissue Expression (GTEx) project (41), and from 16 B-cell progenitors from CHOP (GSE115656) (15). We also analyzed an independent cohort of B-ALL patient RNA-seq datasets (25), with EGA IDs EGAD00001004461, EGAD00001006609, and EGAD00001007530. All the information related to the clinical annotations and sample extraction is described in detail in their respective publication. A summary of the sequencing platforms used for each cohort used in this study is provided in supplementary table 1. A detailed description of the samples selected from each project with their relevant clinical information is provided in Supp. Tables 2 and data file 1. We only used samples classified as B-cell ALL that had at least 25% of the reads mapping to the genome (GRCh38) and such that these corresponded to at least 5M reads. All the samples were processed with the same pipeline outlined in Supp. Figure 1. FastQC (42) was used for quality control of the FASTQ files. FASTQ files FROM SRA for the TARGET samples were extracted using the SRAToolKit (v 2.9.0) (https://github.com/ncbi/sra-tools).

### Fusion detection

We used STAR-Fusion v.1.4.0 (20) to identify gene-fusions from the RNA-seq data. The index was generated using the Gencode (v27) annotation and the GRCh38 assembly. STAR-Fusion was run for each FASTQ file using the default parameters described at https://github.com/STAR-Fusion/STAR-Fusion/wiki/Home/. We required at least one read count supporting the fusion junction given by the field JunctRC (or JunctionReads in the latest version of the manual) in the STAR-Fusion output, one read count connecting the fusion junction (SpanRC or SpanningFrags field), 0.1 Fusion Fragments Per Million total reads (FFPM), and junction reads that cover at least 25 bases on both sides of the breakpoint (indicated as ‘YES_LDAS’ in the STAR-Fusion output). The fusion allele frequency (FAF) was defined as the average of the allele frequency for both partners of the fusion pair, i.e. FAF = (FAF_L_ + FAF_R_)/2, where each value FAF_i_, for i=L,R was defined as FAF_i_ = F_i_ / (F_i_+WT_i_), where the F_i_ represents the number of reads that support the fusion breakpoint and WT_i_ represents the number reads that support the wild type fragment of the gene, not present in the fusion.

### Fusion filtering and classification

Fusion calls involving pseudogenes were removed from the output, as well as fusions between paralogous genes (genes with 70% or more sequence identity), as they were considered potential artifacts. Fusions between immunoglobulin or hemoglobulin genes were also discarded. Additional filters for possible false positives were applied: A promiscuity filter was used to remove fusions involving genes paired with more than one other gene within the same sample, known to be potential artifacts from the library preparation (43). Moreover, we removed all predicted fusions that occurred only in one project to avoid project biases. We also filtered out fusions previously detected in non-cancerous tissues or cells (44), detected in normal samples from TCGA (45), or seen with STAR-fusion in RNA-seq data from normal blood cell types (46). Additionally, we only kept fusions that appeared in 5 or more patients. Fusions involving genes previously reported to have mutations or fusions in any leukemia were kept independently of these filters, but only if they appeared in 5 or more patients. Finally, the FAF was used to select the most relevant fusions (see Methods section Calculation of Fusion Allele Frequency for details). Based on the FAF distribution across all patients from the different cohorts, each fusion was required to have a median FAF greater than 0.1. The partner with the lowest allele frequency in the fusion was required to have a median of 0.01 for the individual FAF value.

We classified the fusions into four major groups: 1) “ALL”, which indicated those already reported in any of the analyzed ALL datasets or reported previously in the databases COSMIC (19), TCGA (45), or MitelmanDB (47); 2) “blood”, which indicated those fusions known to appear in other hematological malignancies according to the same databases; 3) “solid tumors”, which indicated fusions known in other solid tumors and present in the same public databases; and 4) “novel”, which indicated fusions that were not present in the public databases. Fusions were grouped and labeled according to the most recurrent partner for differential expression analysis, co-occurrence analysis, and visualization purposes. Fusion groups that were not showing a pattern of mutual exclusion with any of the other groups according to a waiting time model for mutually exclusive cancer alterations implemented by the R package TiMEx (48) were grouped as “Other”.

### Analysis of fusion disrupted domains

The same fusion breakpoint given by the RNA-seq reads was used if this occurred inside an exon expressed in the fusion. Otherwise, the positions used were the last base of the last exon from gene 1 included in the fusion, and the first base of the first exon from gene 2 included in the fusion. We defined the breakpoints for gene 1 and gene 2 of the identified fusions using these values. PFAM domains mapping to the proteins encoded by each fusion gene were extracted from Biomart (49), the protein coordinates of the domain span were converted to genomic coordinates and overlapped with the fusion breakpoints of the corresponding genes to establish for each breakpoint whether the domain was kept or lost as a result of the fusion.

### Gene expression and functional enrichment analysis

Transcript level quantification for the Gencode transcriptome release 27 (GRCH38.p10) (50) was obtained in transcripts per million (TPM) units using Salmon (v 0.7.2) (51). Gene level quantification was obtained by transforming transcript TPMs to counts per gene using the *tximport* library function from Bioconductor (52).

For differential expression analyses, we considered the patients with only one identified fusion (after performing all the filtering steps) and for the most frequent fusions: 87 patients for *KMT2A-r*, 68 for *ETV6-r*, 9 for BCR-r, 36 for P2RY8-r, 14 for PAX5-r, 25 for PWLC1-r, 5 for RUNX-r, 35 for ST3GAL1-r, 33 for TCF3r, 33 for TTYH3-r, and 12 for ZNF384-r, plus the 133 patients with no fusions detected. We calculated the differentially expressed genes between pairs of groups. The read counts per gene were transformed to log2 counts per million (logCPM) using edgeR (53), and genes with mean logCPM < 0 were filtered out. The data was normalized with the TMM method from the edgeR package. Differential expression analysis was performed with LIMMA (54) using the function *limma*.*voom* adjusted by SVA with the covariables of sex, project, and tissue (bone marrow or peripheral blood).

Gene set enrichment analysis was performed with GSEA (55) for the list of hallmarks and for the biological process ontology using the pre-ranked enrichment method, sorting all the genes by the value of −*log*_10_(*p* − *value*) *· log*_2_*FC* obtained from the differential expression analysis. In the case of splicing factors and RBPs, as there is no pathway or hallmark gene set associated with them in the available databases, we built a list of genes splicing factor and/or RBP function from previous studies (56, 57) and run a pre-ranked GSEA with the absolute value (Data file 10) (Supp. Fig. 13).

### Gene selection to construct a predictive model of prognosis

We used gene expression data from 133 TARGET patients with complete clinical information about the age of diagnosis and time to the first relapse and calculated a log2 fold-change (logFC) per gene using the normalized logCPM mean expression between patients with relapse and without relapse. Similarly, we calculated a logFC from the gene expression of 140 patients from all other cohorts (SJH, LUND, CHOP and PMJCI) comparing the ones carrying only *KMT2A-r* against the ones with only *ETV6-r*. We only considered genes with a logFC > 0.5 in both comparisons and that were included in at least one of three sets: 1) the GSEA hallmark as MYC targets (v1 and v2), in 2) the Gene Ontology Biological Process of translation (initiation, elongation, or termination), or 3) and a list of genes encoding splicing factors and RBPs (Data file 10). For every gene, we applied a cox regression survival model adjusted by age and gender, selecting only the genes with a p-value < 0.05 according to a Wald test (Supplementary Table 3). This produced a total of 39 overexpressed genes associated with prognosis and with our target biological functions and pathways. From these 39 genes, the expression variability between cohorts was evaluated using mean logCPM expression in each dataset and the maximum logFC between datasets. Genes with the highest variability across datasets (max logFC > 3) were removed to avoid any dataset-related bias, obtaining the final 37 genes to build the predictive model. Training data was restricted to the 133 TARGET B-ALL patients. The model consisted of a random forest, implemented using the randomForest library in R, with a total of 400 trees and 4 variables randomly sampled as a candidate at each split. The leave-one-out strategy was used to evaluate the prediction accuracy, while avoiding overfitting. A numeric k-score between 0-1 obtained from the prediction of the random forest model was used to classify the patients according to risk. A threshold was established on 0.7, according to accuracy measures, to classify the patients as high-risk (k-score >= 0.7) or low-risk (k-score < 0.7). To apply the predictive model to expression values given in terms of regularized-log (rlog) values, we used the TARGET samples as before, applying a regularized log transformation using the package DESeq2 with the *rlog* function (58). We then built and tested the model as before, using the rlog values instead of the logCPM values.

### Differential splicing analysis

SUPPA (59, 60) was used to perform the differential splicing analysis. SUPPA predicts the relative inclusion and differential splicing of the events using isoform-level relative abundances. Using RT-PCR experiments, SUPPA’s accuracy was shown to be comparable to methods based on the direct quantification of event PSI from reads (59, 60). SUPPA *generateEvents* was used to generate alternative splicing events defined from protein-coding transcripts and covering the annotated ORFs. The relative inclusion of each event was calculated as a Percent Spliced In (PSI) value with SUPPA *psiPerEvent* using the transcript abundances in TPM units obtained before. A minimum total expression of the transcripts involved in the event of 1 TPM was required. Events without a defined PSI value in more than 10% of the patients across all cohorts were discarded. These included events that did not pass the transcript expression filter or that had all the transcripts involved in the event with zero expression. The remaining missing PSI values were imputed using nearest neighbor averaging with the *impute*.*knn* function in R from the Impute library (61). To test the significant differential inclusion of the events in the comparisons of high against low-risk patients and in *KTM2A*-r against *ETV6*-r patients, a ΔPSI was calculated as the difference of the mean PSI from each group. We discarded all events with a standard deviation (SD) across groups lower than 0.1. We applied a linear regression model with a logit transformation of the PSI to estimate the significance of the splicing changes and adjusted the p-value by calculating a false discovery rate (FDR), using the same covariables adjustment as in the differential expression analysis described previously. We considered significant all the changes with |ΔPSI| > 0.2 and an FDR corrected p-value < 0.01. Differential splicing between B-cell precursors and GM12878 was calculated with the SUPPA *diffSplice* command with default options.

### Motif Enrichment analysis

We searched for RBP binding motifs on the regions neighboring each splicing event with MoSEA (https://github.com/comprna/MoSEA) (57). MoSEA was run against a database of Position Frequency Matrices (PFM) and k-mers (6-mers) associated with each RBP. Enrichment was assessed by comparing a set of events differentially spliced between conditions with a set of events with no significant change between the same conditions. For each motif, MoSEA calculated a z-score from the comparison of the observed frequency observed in differentially spliced events with the distribution of frequencies in 100 control subsamples of the same size, considering the length distribution and GC content of the differentially spliced events set. We considered those motifs PFMs and 6-mers with z-score > 1.5.

### Retrieving Protein-Protein Interactions

We used the STRING database (62) to retrieve Protein-Protein Interactions with the detailed scores of the links between proteins. Only those with experimental scores different from 0 and a combined score higher than 900 were kept.

### Co-occurrence analysis of differentially spliced events and high risk

An alternative exon was considered included for PSI > 0.5 and absent otherwise for each sample. For every fusion group, a matrix was built with the presence or absence of events in each patient. The co-occurrence of the events with high risk was tested with a probabilistic model of species co-occurrence implemented in the R package co-occur (63).

### Functional enrichment analysis of differentially spliced genes

Genes associated with differentially spliced events were tested for functional enrichment of Gene Ontology Biological Process terms with the R package clusterProfiler from Bioconductor (64). Benjamini-Hochberg (BH) correction was used to calculate adjusted p-values (q-values). Only ontologies with p-value and q-value below 0.05 were selected.

### Leukemia cell lines selection

SEM, MHH-CALL-3, KOPN-8, NALM-19, REH, and SUP-B15 cells were obtained from the Leibniz Institute DSMZ (#ACC546, #ACC339, #ACC552, #ACC522, #ACC22 and #ACC389, respectively) and cultured according to the supplier’s recommendations. These cell lines were previously checked for mycoplasma contamination by PCR as previously reported (65). Results were expressed as a percentage with respect to scramble-transfected controls.

### RNA isolation, real-time qPCR, and customized qPCR dynamic array based on microfluidic technology

Total RNA from leukemia cell lines was extracted with TRIzol® Reagent (ThermoFisher Scientific, #15596026). Total RNA concentration and purity were assessed by Nanodrop One Microvolume UV-Vis Spectrophotometer (ThermoFisher Scientific). For qPCR analyses, total RNA was retrotranscribed by using random hexamer primers and the RevertAid RT Reverse Transcription Kit (ThermoFisher Scientific, #K1691). Thermal profile and qPCR analysis to obtain absolute mRNA copy number/50 ng of sample of selected genes are reported elsewhere (66). To control the possible variations in the efficiency of the retrotranscription reaction, mRNA copy numbers of the different transcripts analyzed were adjusted by *ACTB* expression. Specific primers for human and mouse transcripts including *ACTB* and *SRRM1* genes were specifically designed with the Primer3 software [*SRRM1* (NM_001303448.1) - forward: GTAGCCCAAGAAGACGCAAA, reverse: TGGTTCTGTGACGGGGAG; *ACTB* (NM_001101) -forward: ACTCTTCCAGCCTTCCTTCCT, reverse: CAGTGATCTCCTTCTGCATCCT].

### Silencing of splicing factors by specific small interfering RNA

Pre-designed and validated specific small interfering RNA (siRNA) oligos for knockdown of endogenous *SRRM1* (#s20018; Silencer® Select siRNAs; ThermoFisher Scientific) were used, which is a pre-validated siRNA. Briefly, cells (n = 500 000 cells/well) were transfected with 25 nM of each siRNA individually using Lipofectamine® 3000 Transfection Reagent (ThermoFisher Scientific, # L3000075) according to the manufacturer’s instructions. Silencer® Select Negative Control siRNA (ThermoFisher Scientific, #4390843) was used as a scramble control. After 24 h, cells were collected for validation of the transfection by qPCR and seeded for different functional assays.

### Proliferation rate determination

Cell proliferation in response to *SRRM1* silencing in leukemia cell lines was analyzed using the alamarBlue™ assay (Biosource International, Camarillo, CA, USA), as previously reported (67). Briefly, cells were seeded in 96-well plates at a density of 25,000 cells/well and serum-starved for 24h. Then, proliferation was evaluated every 24h using the FlexStation-III system (Molecular Devices, Sunnyvale, CA, USA) for up to 72h. Results were expressed as a percentage referred to as scramble-transfected controls.

### Apoptosis measurement

Apoptosis induction in response to *SRRM1* silencing in leukemia cell lines (25,000 cells/well onto white-walled multiwell luminometer plates) was performed by using Caspase-Glo^®^ 3/7 Assay (Promega Corporation, #G8091) as previously reported (67). Briefly, Cells were seeded in 96-well white polystyrene microplate flat bottom clearplates at a density of 25,000 cells/well and serum-starved for 24h. Results were expressed as a percentage referred to as scramble-transfected controls.

### Invasion rate determination

The invasion rate in response to *SRRM1* silencing in leukemia cell lines was assessed by using the 96-well cell Trans-well invasion assay (Basement Membrane-8 μm, AssayGenie, #BN01086) according to the manufacturer’s protocol. The top chamber membrane was coated with the basement membrane solution. Afterward, cells were seeded in the top chamber in 0,5% serum media. Then, 10% FBS media was placed in the bottom chamber to promote cell invasion in all experimental conditions. Then, proliferation was evaluated every 24h using the FlexStation-III system (Molecular Devices, Sunnyvale, CA, USA).

### Statistical analysis for the B-ALL cell line experiments

Numerical results were evaluated for statistical differences by t-test, multiple t-tests, and 2-way ANOVA. All statistical analyses were performed using Prism software v.9.0 (GraphPad Software, La Jolla, CA, USA). P-values < 0.05 were considered statistically significant. The plotted data represent the median (interquartile range) or means ± standard error of the mean (SEM). Significant difference from control conditions was indicated as * for P-value < 0.05, ** for P-value < 0.01, *** for P-value <0.001.

## Data availability

All samples were obtained from various sources through controlled or public access. The series from St Jude Children’s Research Hospital (SJH) (EGAS00001000246) (18) and Lund University (LUND) (EGAS00001001795) (14) were downloaded from the European Genome-phenome Archive (EGA), and the corresponding clinical information was obtained from the associated publications. The series from Children’s Hospital of Philadelphia (CHOP) (GSE115656) was downloaded from Gene Expression Omnibus (GEO) (15). Samples from TARGET (Therapeutically Applicable Research to Generate Effective Treatments) were downloaded from the TARGET data portal at National Cancer institute (NIH) together with the associated clinical information, corresponding to dbGAP accessions phs000463 (ALL phase 1) and phs000464 (ALL phase 2). Data from patients from the Princess Maxima Center for Pediatric Oncology (PMJCI) from (16) was obtained from the authors. Data from 12 B-ALL cell lines were obtained from the Cancer Cell Line Encyclopedia (CCLE) (40). Data from normal blood and spleen samples were obtained from The Genotype-Tissue Expression (GTEx) project (41), and from 16 B-cell progenitors were obtained from the GEO dataset (GSE115656) (15).

## Acknowledgments

We thank the authors of the different studies used in this article for facilitating access to their datasets: SJH (18), LUND (14), CHOP (15), TARGET, and PMJCI (16).

## Funding

This work was funded by the Spanish Ministerio de Ciencia, Innovación y Universidades with grant BIO2017-85364-R (to E. Eyras). The funding body did not contribute to the design of the study, analysis, or interpretation of data, nor to the writing of the manuscript.

## Description of data files

**Data file 1**. Table with detailed clinical information for all patients obtained from the original sources. *Sample_ID* is the code used to identify the sample. *Patient_ID* is the code used to identify the patient. Project indicates the sample cohort of origin. *Type* indicates the type of leukemia. *Tissue* indicates the origin of the sample. *Blasts* indicates the proportion of blasts in the sample. DX_RL indicates the time when the sample was obtained at diagnosis (DX) or after relapse (RL). *Gender* indicates whether the patient is female (F) or male (M). *Age* indicates the age of the patient in months. *Fusions* indicates the fusion recorded in the clinical information. *Cell*.*of*.*Origin* indicates if the samples were obtained from B-cells or B-cells precursors. *First*.*event* indicates if the patient had a relapse or not. *Event*.*Free*.*survival*.*time*.*in*.*days* is the time in days when the first event appeared. *Vital*.*Status* indicates if the patient is dead or not. *Overall*.*Survival*.*Time*.*in*.*Days* is the time until the patient’s death or the last time of follow-up.

**Data file 2**. Table with all fusions detected. Columns indicate sample code (*Sample*), patient ID (*ID*), cohort, source of the fusion calling (Star-fusion, Clinical information, or both), cancer type in which the fusion has been observed before (*Cancer_category*), fusion group in which the fusion has been assigned (Group), number of reads supporting the fusion junction (*JunctRC*), number of spanning reads supporting the fusion (*SpanRC*), expression of the fusion in Fusion Fragments per Million (*FFPM*), Ensembl gene identifier of the first fusion gene (*ENG1*), breakpoint coordinates of the first fusion gene (*BP1*), Ensembl gene identifier of the second fusion gene (*ENSG2*), breakpoint of the second fusion gene (*BP2*), type of chromosomal rearrangement (annotation), FAF of the first fusion gene (*FAF_gene1*), FAF of the second fusion gene (*FAF_gene2*), and combined FAF (*FAF*).

**Data file 3**. Tables with all differential expression analyses between fusion groups. The comparisons and labels correspond to those shown in supplementary figure 9. The GeneID column indicates the gene name. LogFC is the log2-fold-change between the two conditions. AveExpr is the average log2-expression for the gene over all samples. T is the moderated t-statistic. P.Value is the raw p-value. Adj.P.Val is the adjusted p-value using the BH method. B is the log-odds that the gene is differentially expressed.

**Data file 4**. Differential expression analysis between *KMT2A*-r and *ETV6*-r patients. The GeneID column indicates the gene name. LogFC is the log2-fold-change between the two conditions. AveExpr is the average log2-expression for the gene over all samples. T is the moderated t-statistic. P.Value is the raw p-value. Adj.P.Val is the adjusted p-value using the BH method. B is the log-odds that the gene is differentially expressed.

**Data file 5**. Pearson correlation for the signature genes with the risk score. Column *statistic* is the value of the test statistic, *p*.*value* is the p-value of the test and *estimate* is the R of the correlation test.

**Data file 6**. Signature scores for all the patients and cell lines studied. Column *Sample* indicates the sample code. Column *None* indicates the score associated with not having a relapse. Column *Relapse* indicates the score related to not having a relapse. Column *Sample_type* indicates if the origin of the sample is a patient at the diagnostic stage (Diagnostic), a patient at relapse stage (Relapse), a cancer cell line (CCLE), or a normal sample (GM12878 cell line, B-cell progenitors).

**Data file 7**. Differential splicing events between high and low-risk patients. Columns indicate the event ID, dPSI, p-value, and FDR-adjusted p-value as calculated with the linear model (see Methods), HGNC gene symbol, transcripts supporting the inclusion of the event (Alt 1), transcripts supporting the exclusion of the event (Alt 2), column indication whether any of Alt 1 or Alt 2 transcripts are annotated as coding (transcript_type), cancer hallmarks associated to the differentially spliced genes (hallmarks), fusions in which the event shows co-occurrence with high risk (high_risk_cooccur) and presence of motifs for SRRM1 interacting RBPs(SRRM1_RBPs_motifs).

**Data file 8**. Differential splicing events between *KMT2A*-r and *ETV6*-r patients. Columns indicate the event IDs, dPSI, p-value, and FDR-adjusted p-value as calculated with the linear model (see Methods).

**Data table 9**. Differential splicing events between B-cell precursors and GM12878 cells. Columns indicate the event ID, dPSI, and p-value provided by SUPPA. Events for which the method could not retrieve a dPSI value were omitted (see Methods).

**Data file 10**. A manually curated list of RBPs and splicing factors used for the analysis.

## Notes

https://github.com/comprna/risk_model_app

### Competing Interest Statement

The authors have declared no competing interest.

### Summary of Updates

We have provided further validation of our results using an independent data cohort.

