## Supplementary Figures and Tables for "A convergent malignant phenotype in B-cell acute lymphoblastic leukemia involving the splicing factor SRRM1"

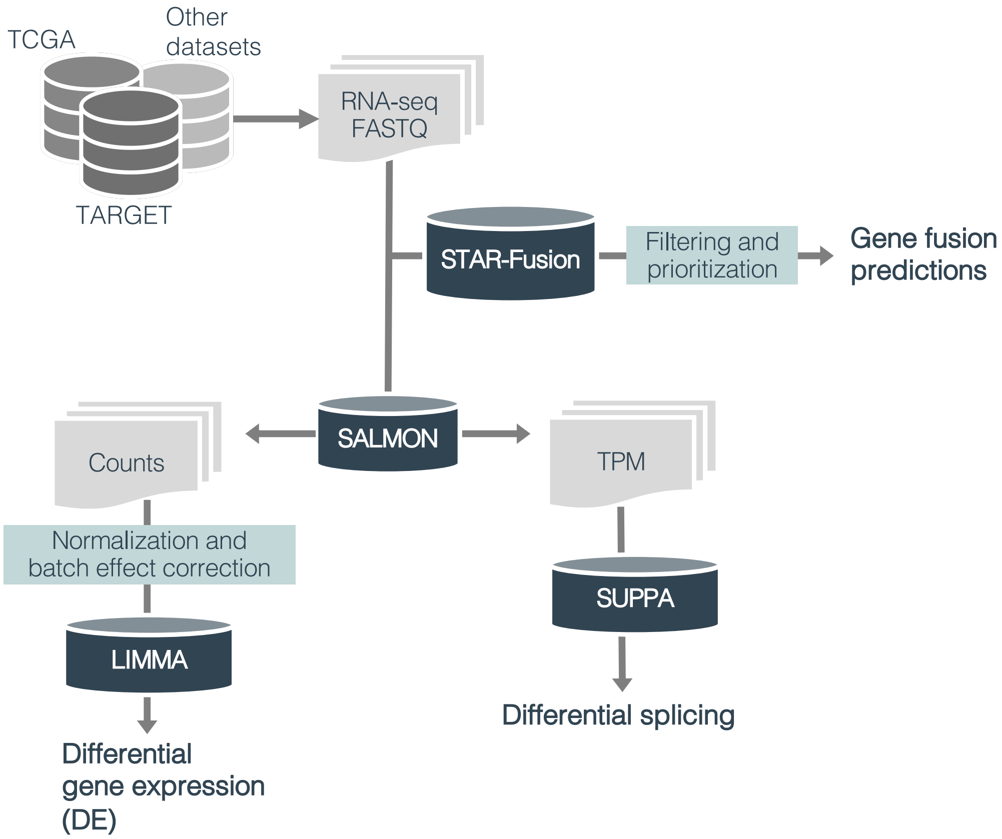


**Supplementary Figure 1. RNA sequencing (RNA-seq) analysis pipeline.** The figure depicts the analyses performed on the short-read RNA-seq samples from Data Table 1. The description of the methods used can be found in the Methods section.

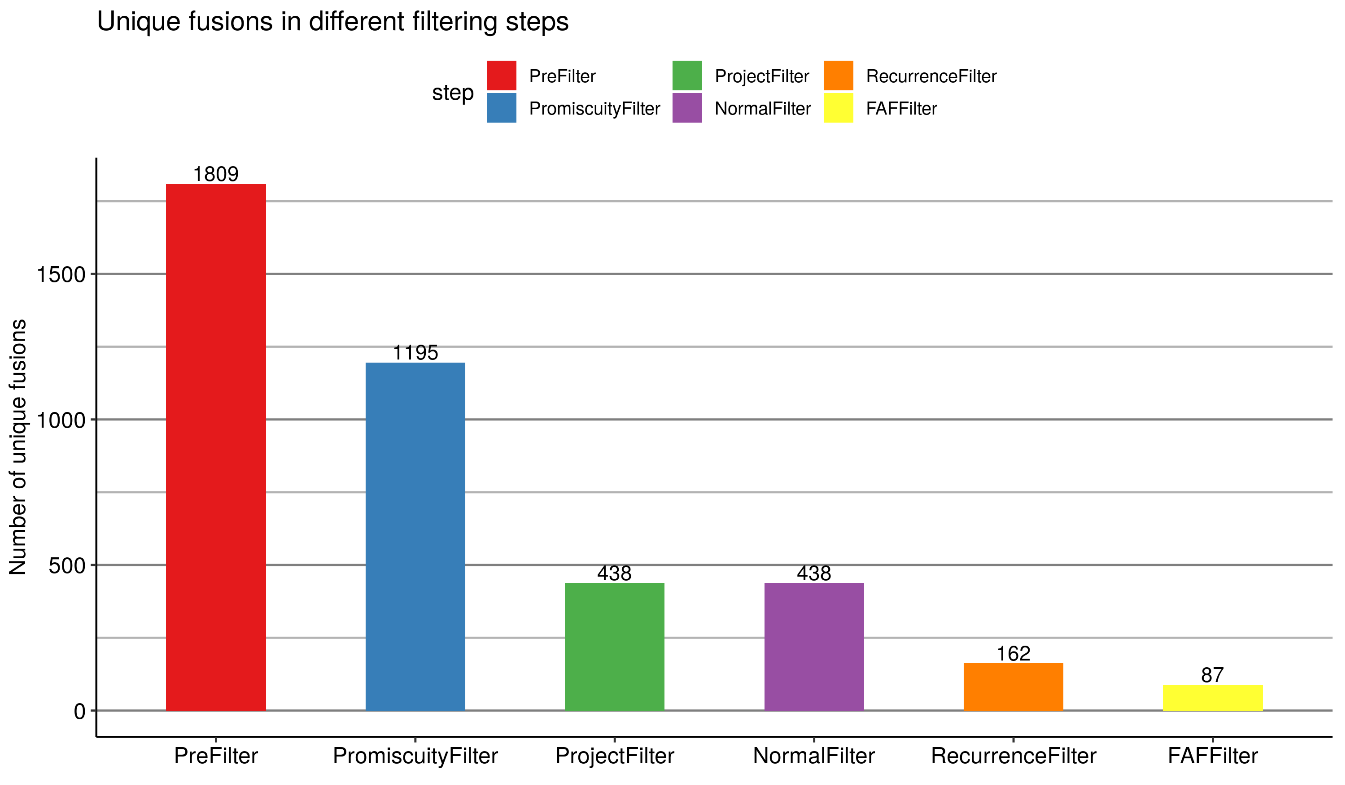


**Supplementary Figure 2. Detection of gene-fusions.** Bar plot showing the number of unique fusions identified after each filtering step was applied. We started with 1825 fusion candidates. On top of each bar, we indicate the number of fusions left after applying each filter consecutively, from left to right. PreFilter (red): fusions involving Ig genes, Hb genes, pseudogenes and paralogous genes were removed. PromiscuityFilter (blue): fusions involving genes with multiple partners in the same sample were removed, except if the fusions were previously observed in cancer. ProjectFilter (green): fusions that appeared in only one of the cohorts were removed. NormalFilter (purple): fusions appearing in normal samples were removed. RecurrenceFilter (orange): fusions occurring in fewer than 5 patients were removed, except for fusions involving genes that were observed before in other fusions or mutated in leukemia. FAFFilter (yellow): cases with low fusion allele frequency (FAF) were removed.

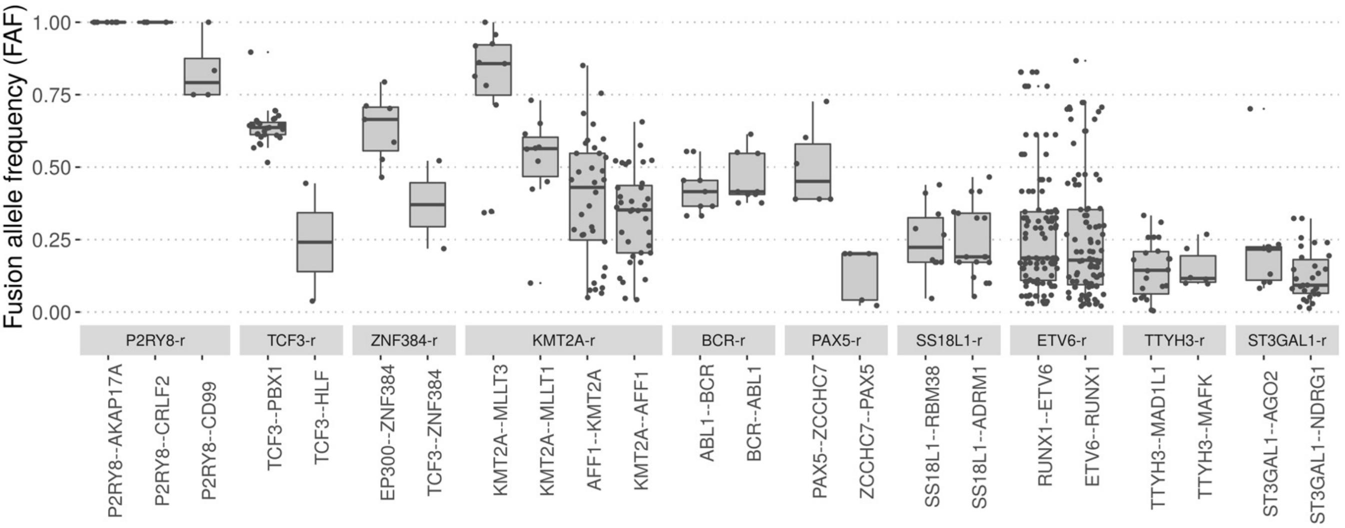


**Supplementary Figure 3. Fusion allele frequency (FAF) values.** For each of the fusion groups depicted in Figure 1, we give the distribution of the FAF values (y axis) for the most common gene fusion pairs (x axis).

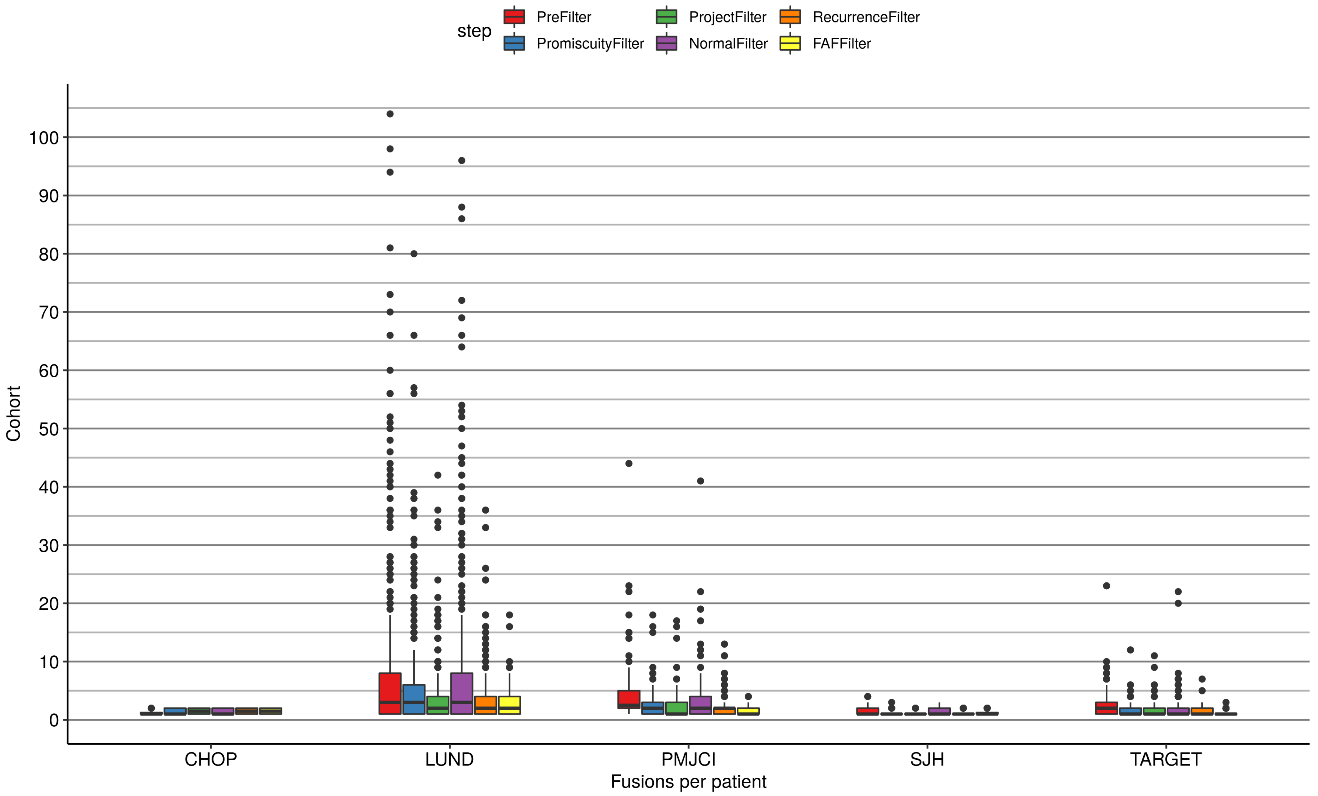


**Supplementary Figure 4. Number of fusions per patient.** Distribution of the number of fusions in each cohort after applying the filters described in Supp. Fig. 2. The fusions used in this work correspond to the yellow distributions, resulting from applying all filters.

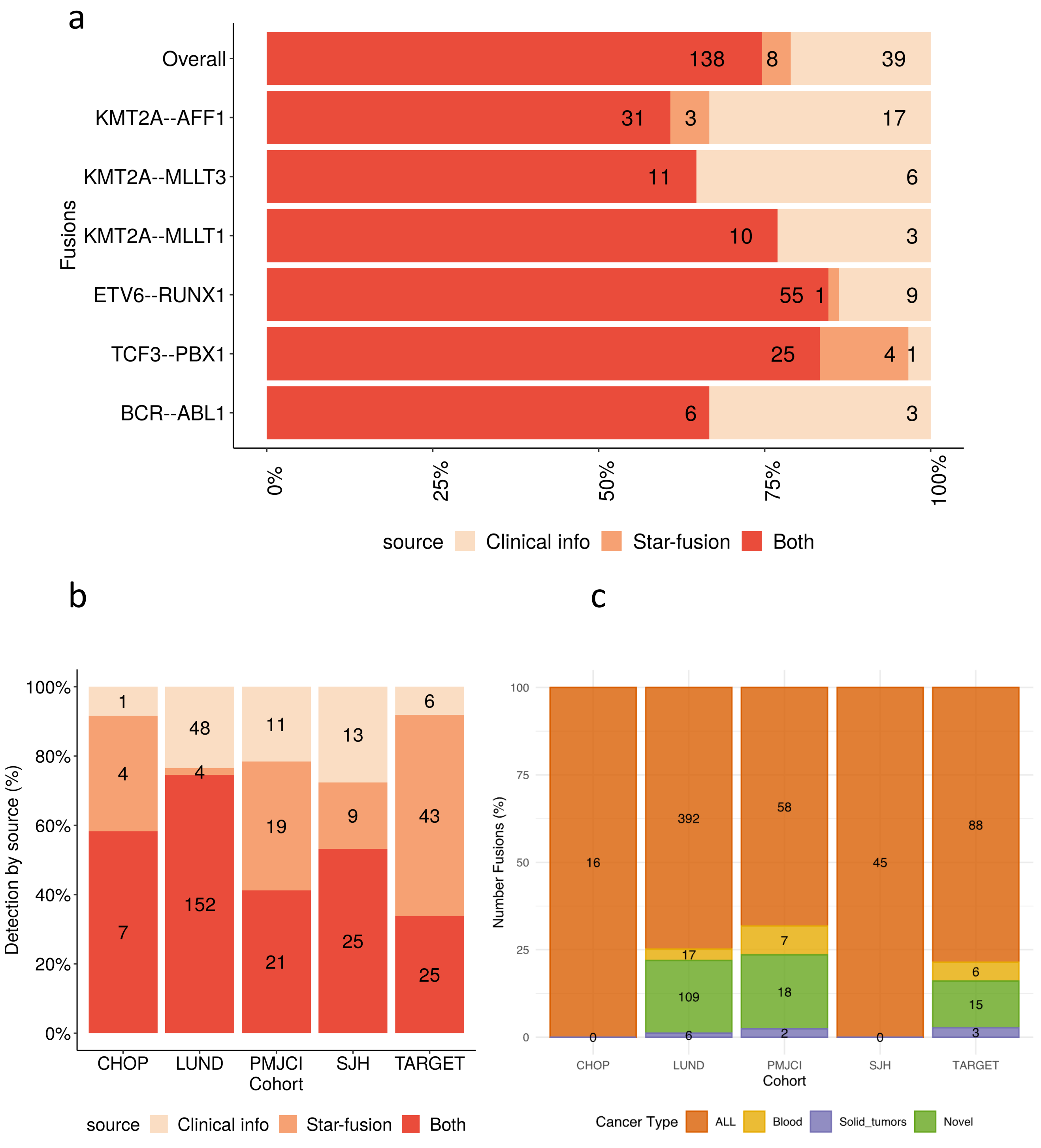


**Supplementary Figure 5. Detection of annotated fusions measured by independent experimental methods.** **(a)** For each fusion, we show the proportion of patients in which 1) the fusion was detected from RNA-seq and was annotated in the clinical information based on independent experimental methods (Both), 2) the fusion was only detected from RNA-seq (Star-fusion), and 3) the fusion was annotated in the clinical information but not detected in RNA-seq (Clinical info). **(b)** Proportion of fusions in each of the B-ALL cohorts separated as in (a). **(c)** Proportion of fusions by project colored according to whether the fusion has been previously described in ALL, other blood cancers (Blood), solid tumors, or has not been previously described in any cancer type (Novel).

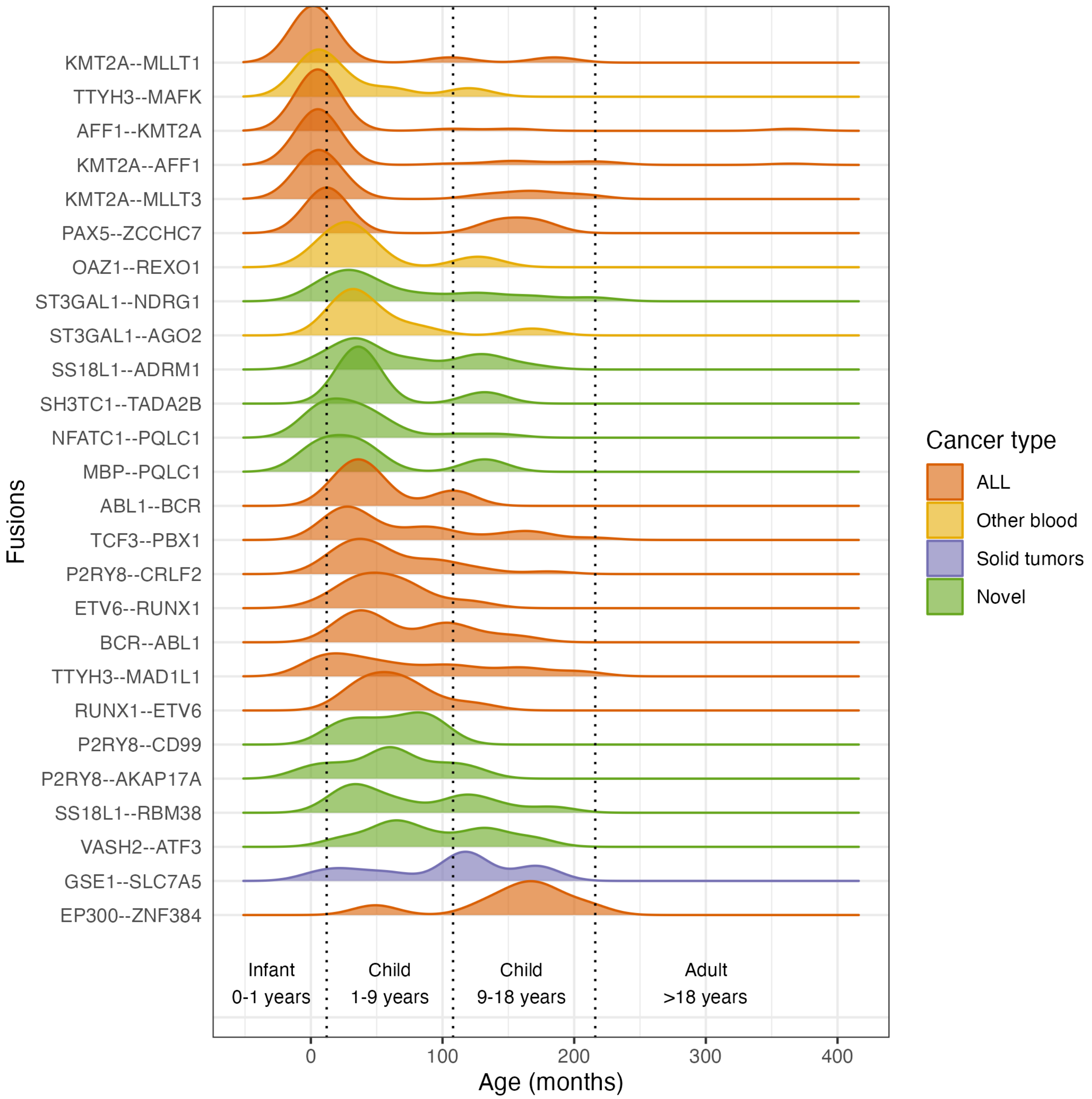


**Supplementary Figure 6. Fusion range distribution by age.** Age distribution of the patients (x axis) by fusion (y axis), colored according to whether the fusion has been previously described in ALL, other blood cancers (Other blood), solid tumors, or has not been previously described in any cancer type (Novel).

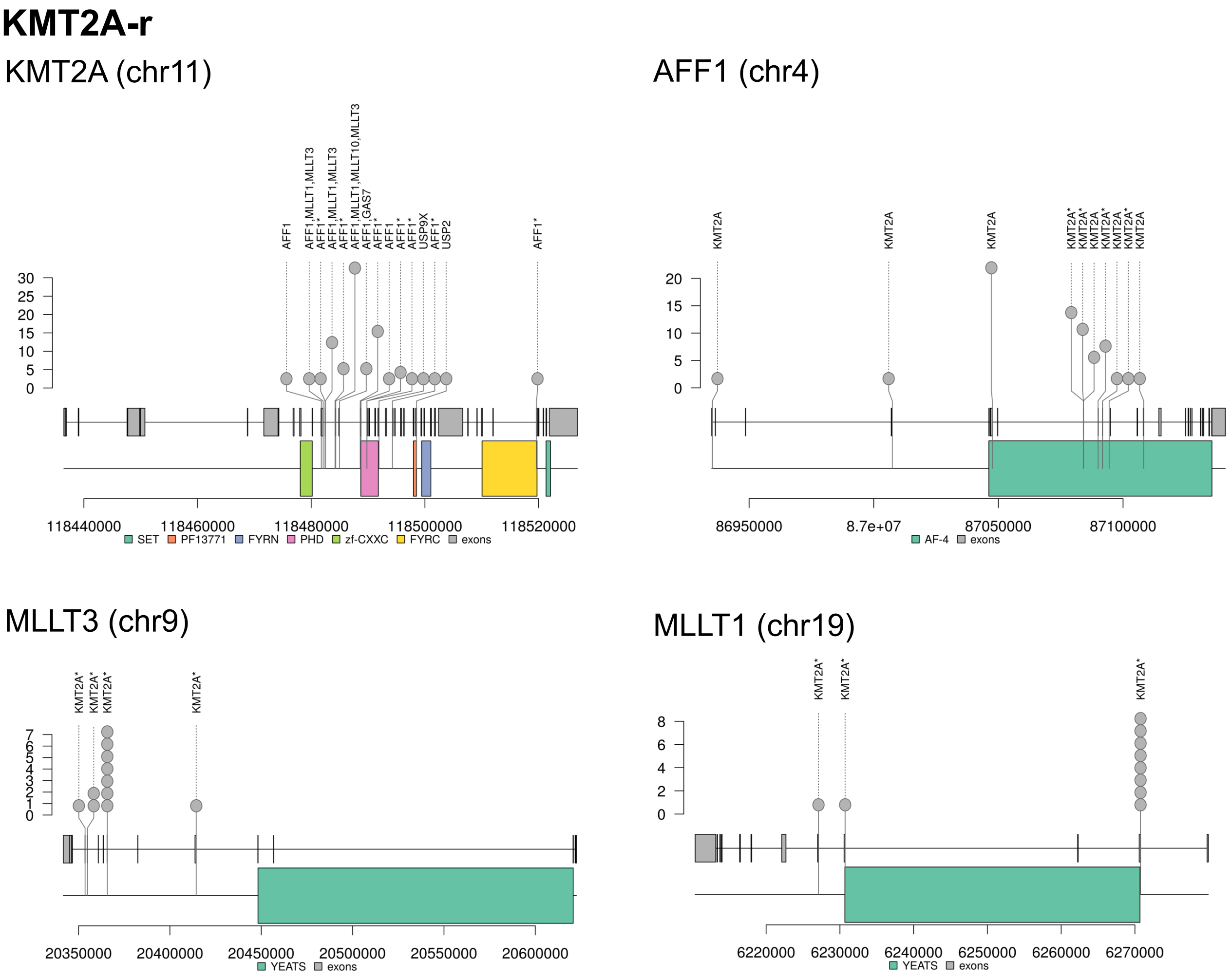


**Supplementary Figure 7. Fusion breakpoints.** Breakpoints of detected fusion genes for fusions present in at least 5 patients. Along the gene models (exons in grey boxes and genomic extent of PFAM domains, when available, in coloured boxes), we indicate the position of the fusion breakpoint detected (grey dots). The fusions partners are written next to each breakpoint. A fusion partner is indicated with * if it is the first partner of the gene. Otherwise, the depicted gene is the first partner of the fusion. Partners with recurrence lower than 5 are indicated but not depicted. The plots show the actual breakpoints only if they occur inside an exon. Otherwise, the given position is the last base of the last exon from the first gene included in the fusion, and of the first base of the first exon from the second gene included in the fusion. The x axis indicates the genomic positions and the y axis indicates the recurrence of the breakpoint. Fusions are grouped according to Figure 1b, genes present in more are depicted only once.

### ETV6-r

#### ETV6 (chr12)

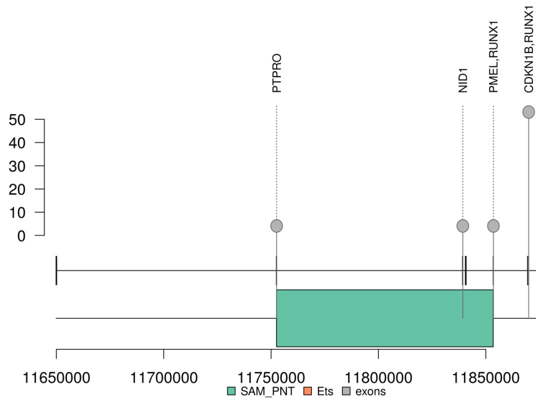


#### RUNX1 (chr21)

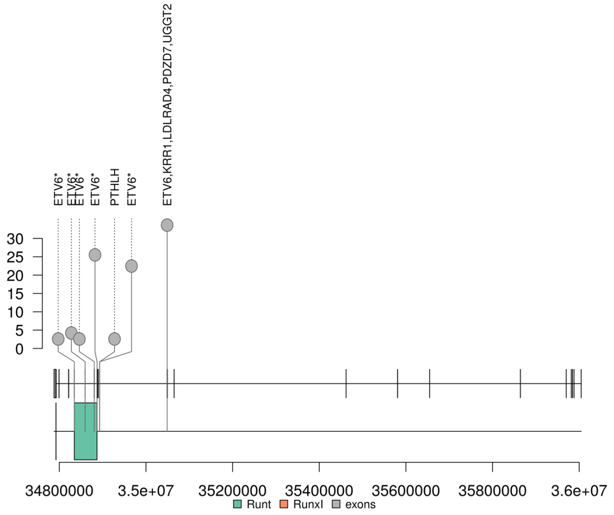


### ST3GAL1-r

#### ST3GAL1 (chr8)

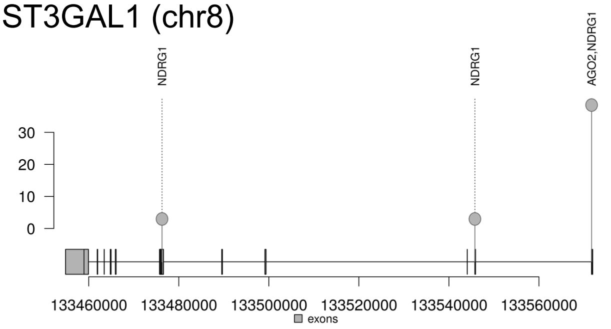


#### NDRG1 (chr8)

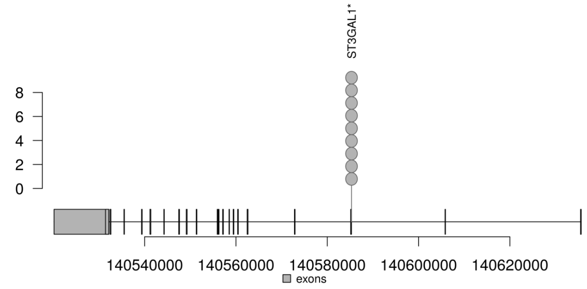


#### AGO2 (chr8)

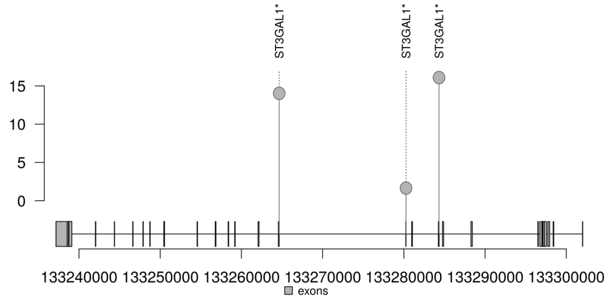


**(Supplementary Figure 7 cont.)**

## P2RY8-r

P2RY8 (chrX)

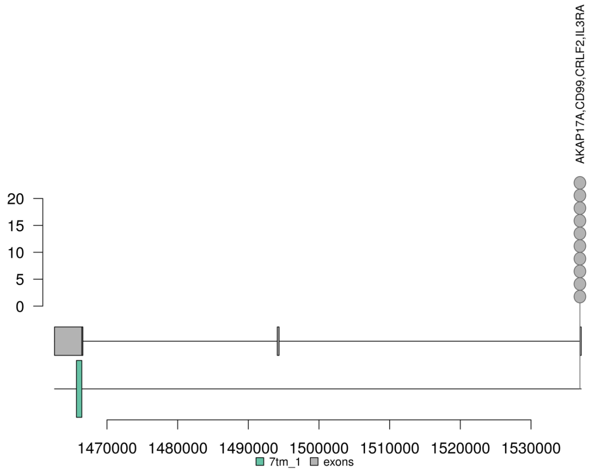


CRLF2 (chrX)

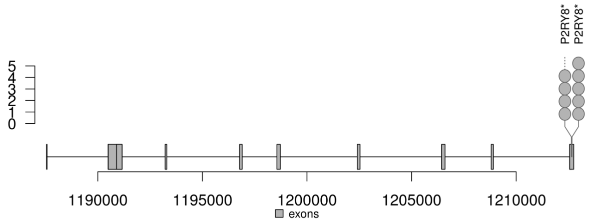


CD99 (chrX)

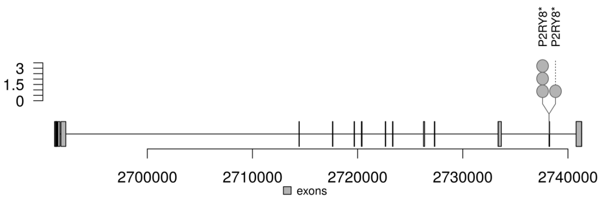


AKAP17A (chrX)

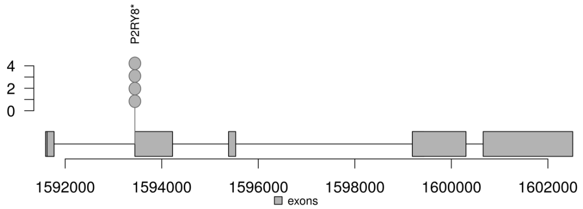


#### TCF3-r

TCF3 (chr19)

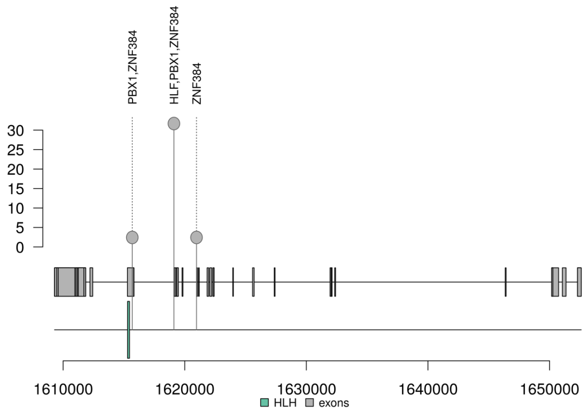


PBX1 (chr1)

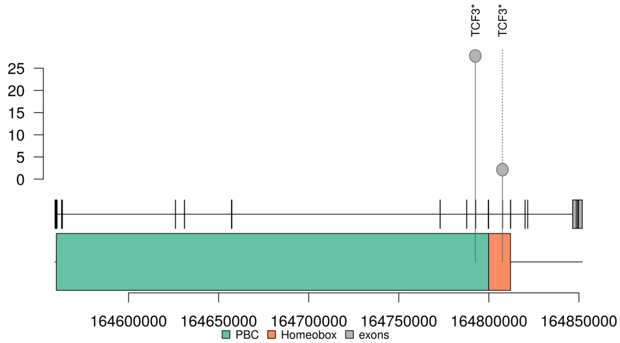


**(Supplementary Figure 7 cont.)**

### TTYH3-r

TTYH3 (chr7)

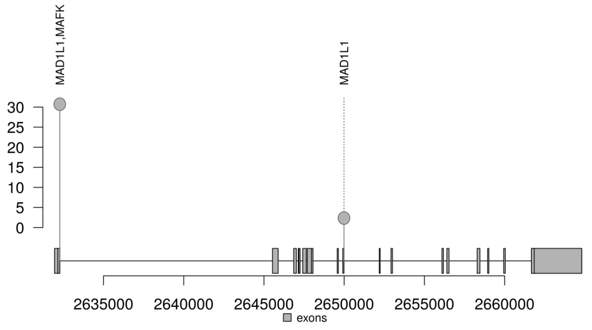


MAD1L1 (chr7)

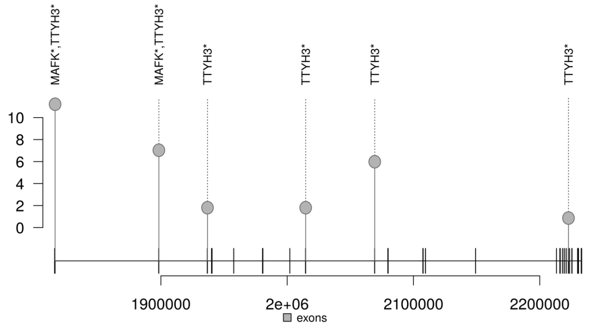


MAFK (chrX)

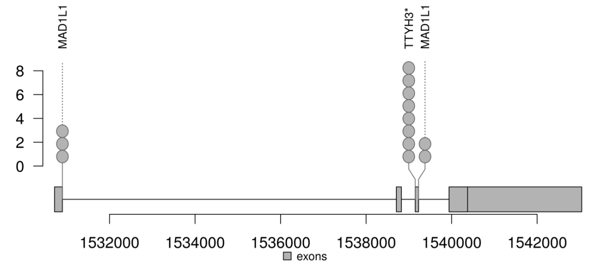


### ZNF384-r

ZNF384 (chr12)

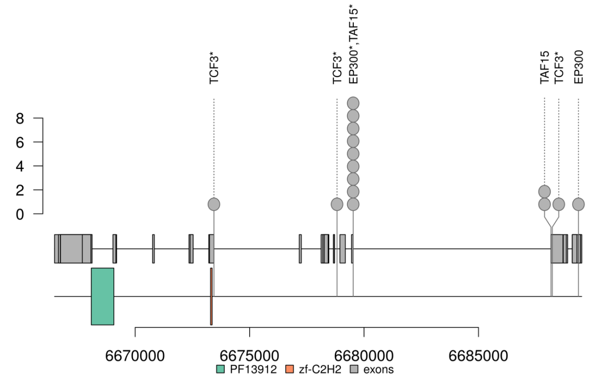


EP300 (chr22)

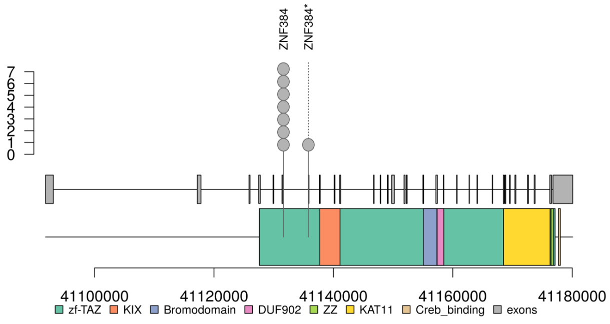


### (Supplementary Figure 7 cont.)

#### BCR-r

BCR (chr22)

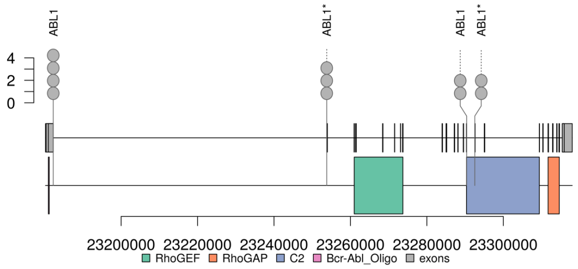


ABL1(chr9)

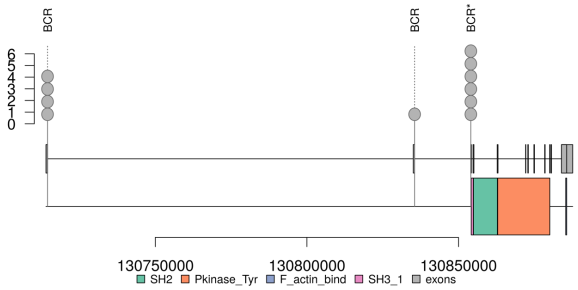


#### GSE1-r

GSE1 (chr16)

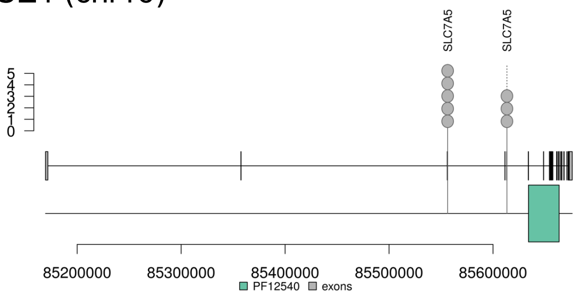


SLC7A5 (chr16)

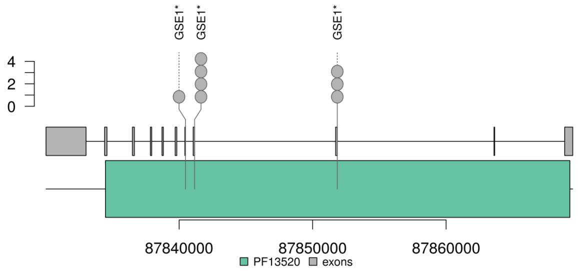


**(Supplementary Figure 7 cont.)**

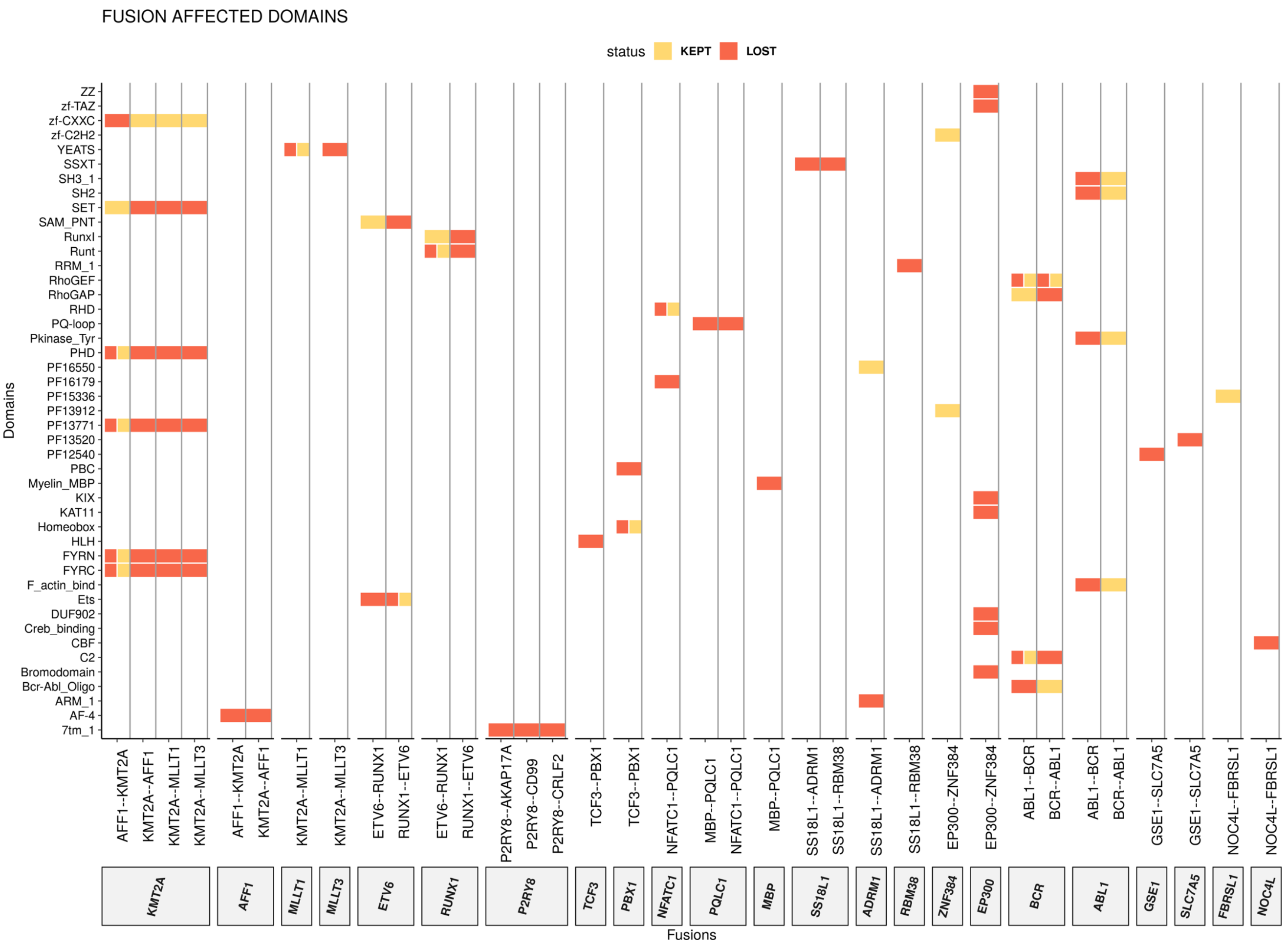


**Supplementary Figure 8. Domains kept and lost through gene fusions.** For each fusion (x axis), we indicate the domain (y axis) that is kept (yellow) or lost (red) in the fusion gene indicated in the box below the fusions.

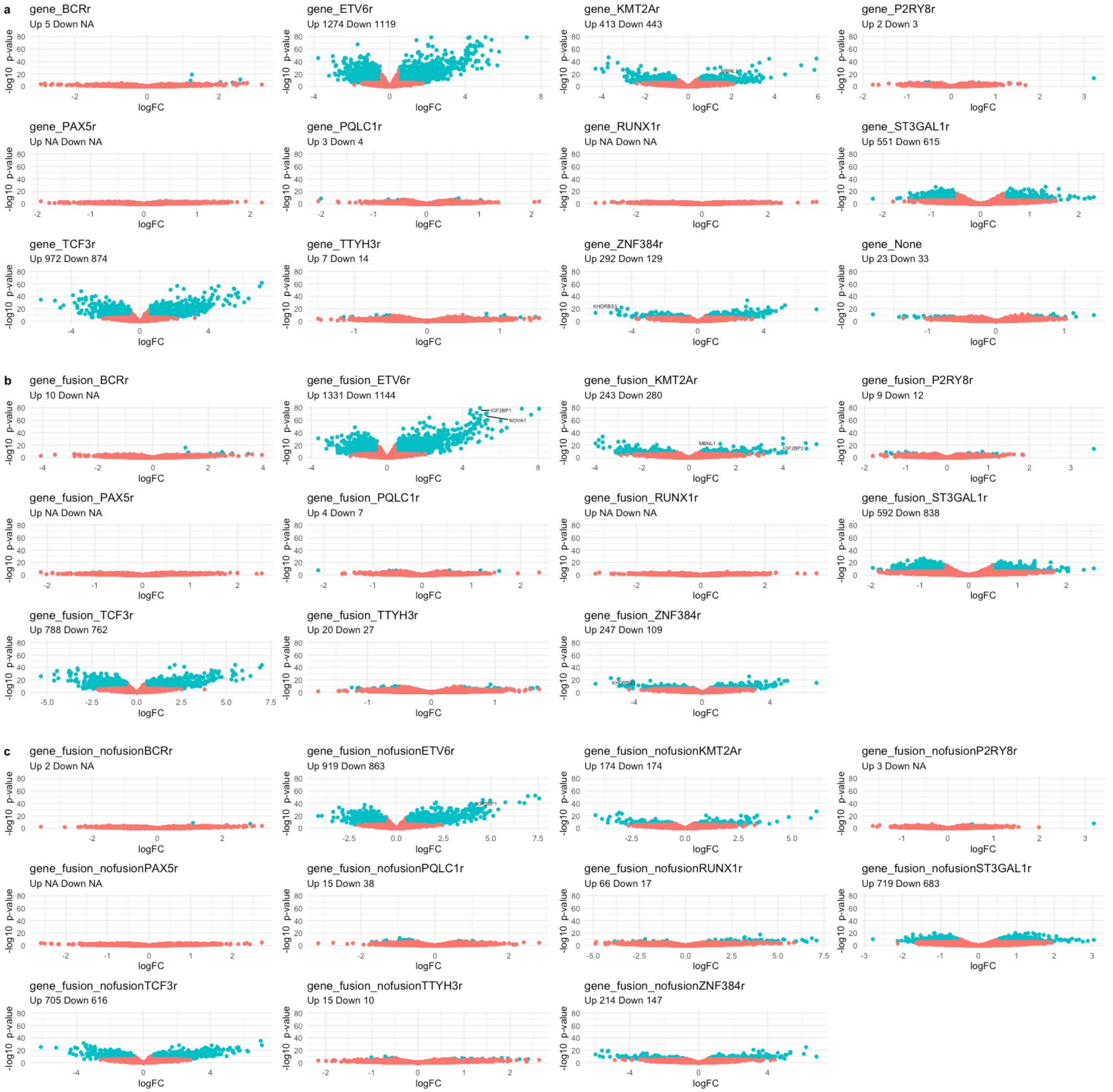


**Supplementary Figure 9. Systematic differential expression analysis between fusion groups.** For each fusion group (Supp. Table 2), we show the volcano plots for the differential expression analysis in three comparisons: **(a)** the fusion group vs. all leukemia patients (12 comparisons), **(b)** each fusion group vs. patients with some other fusion (11 comparisons), and **(c)** the fusion group vs. leukemia patients without fusion (11 comparisons). Patients that fall in more than one fusion group, because they have multiple fusions classified in different fusion groups, were removed from the comparisons. Volcano plots show on the x axis the log2 fold-change and on the y axis the -log10(corrected p-value). On each subtitle appear the total number of up and down regulated genes per comparison after applying a Bonferroni filter with an absolute log2 fold-change > 0.5.

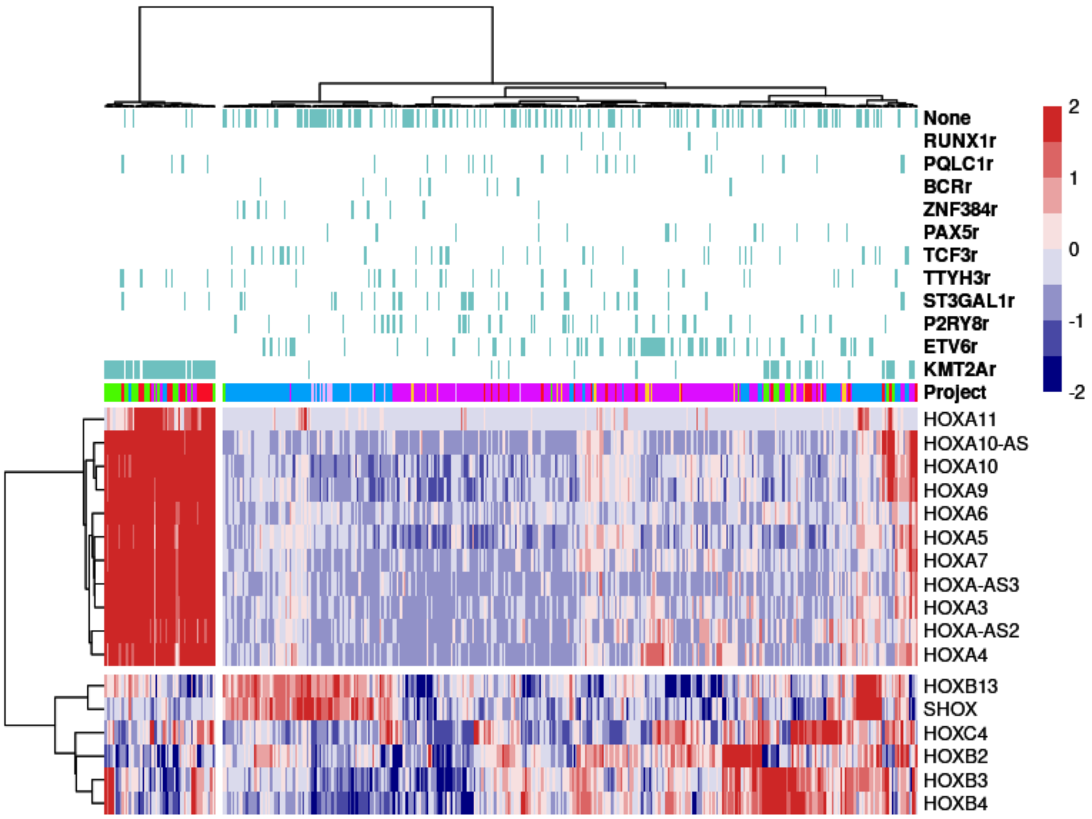


**Supplementary Figure 10. Expression of HOX genes.** We cluster all patients according to the expression of the HOX genes. Expression is given as the z-score of the log2(CPM) (CPM: counts per million). For each patient, we indicate above their cohort (Project) and fusion group. We indicate with "None" those patients without any recorded fusion. The Euclidean distances and Ward clustering method were used to generate this heatmap.

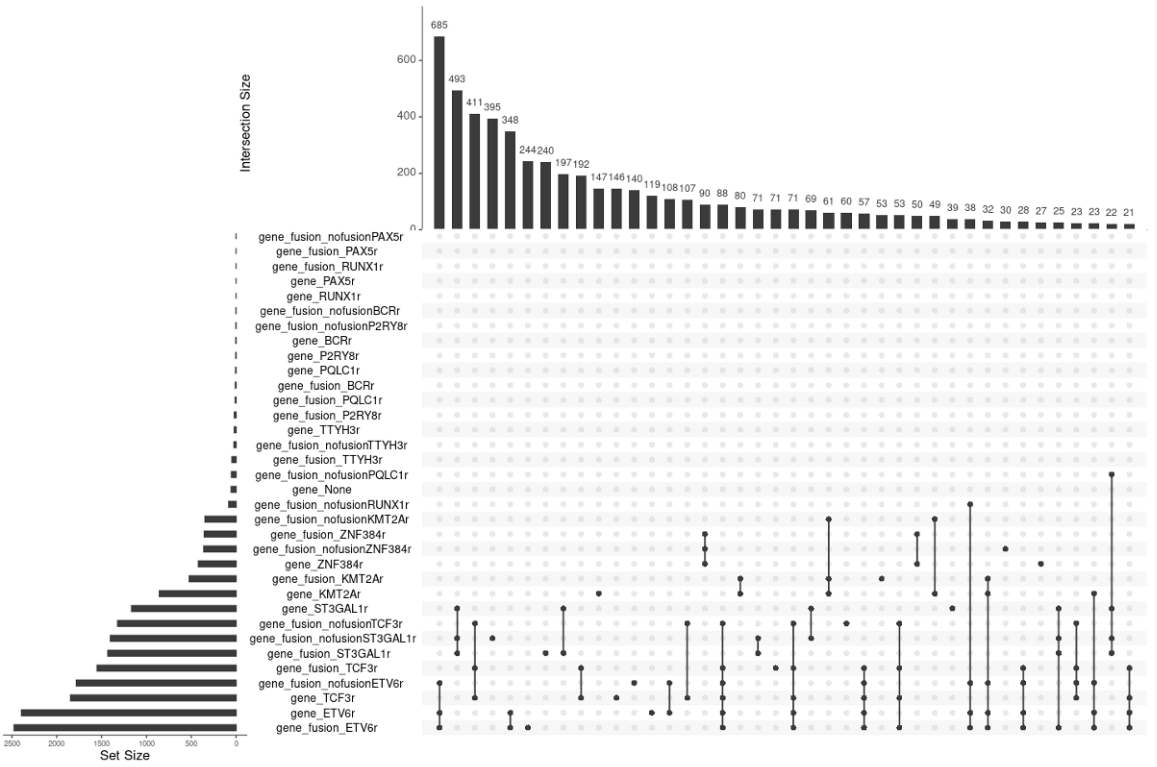


**Supplementary Figure 11. Overlaps between differentially expressed genes in each fusion group.** Intersection of genes differentially expressed from each comparison in Supp. Fig. 9. For each comparison on the left, the horizontal bar plot indicates the number of differentially expressed genes, and the vertical bar plot indicates the intersection of genes in two or more of these comparisons (indicated by bullet points). The intersections did not consider the direction of change.

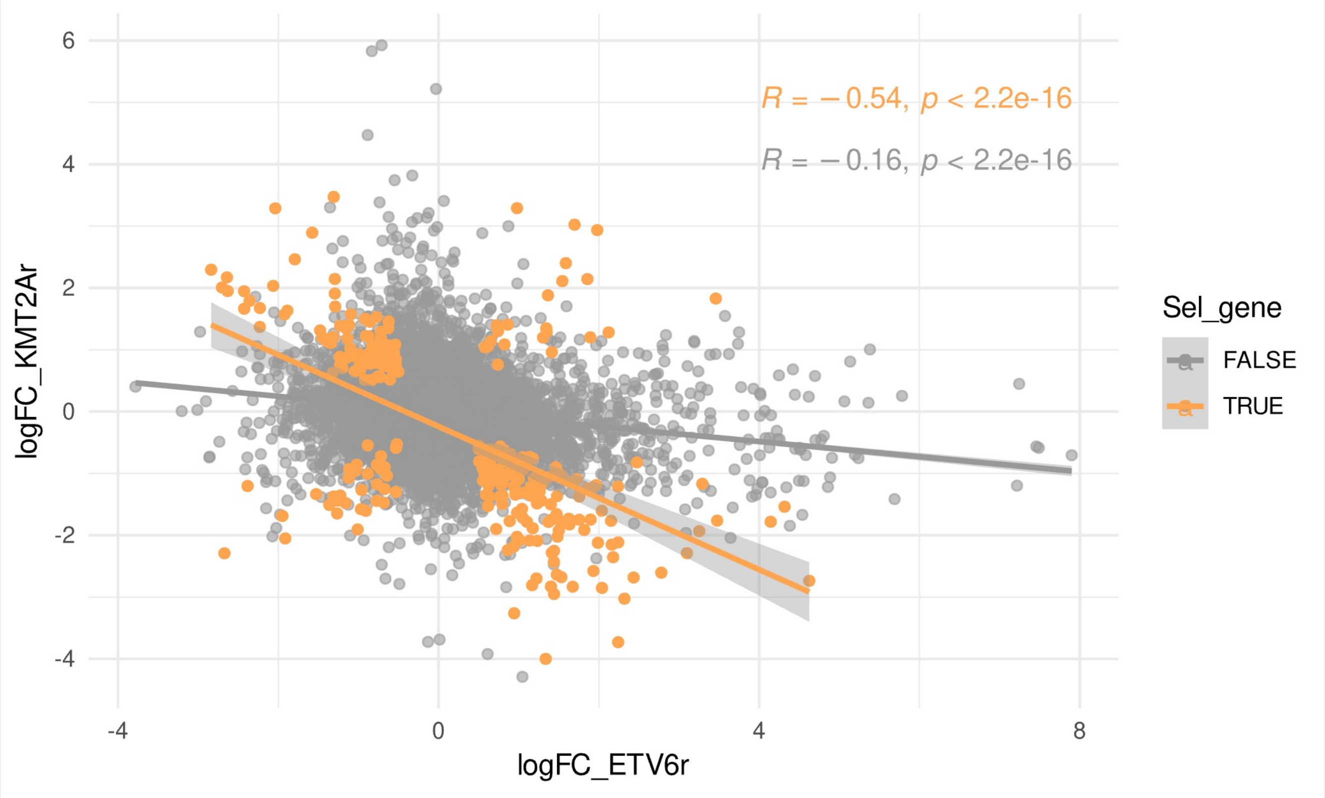


**Supplementary Figure 12. Overlap of genes that are differential expressed between *KMT2A*-r or *ETV6*-r and the other groups of leukemia patients**. The scatterplot shows the log2 fold-change (logFC) from comparing *ETV6*-r with the rest of leukemia patients (x axis), and from comparing *KMT2A*-r with the rest of leukemia patients (y axis). In orange we show the genes that were significant in both comparisons, in grey all the genes that showed no significance in any of the comparisons. R indicates the Pearson correlation value.

a

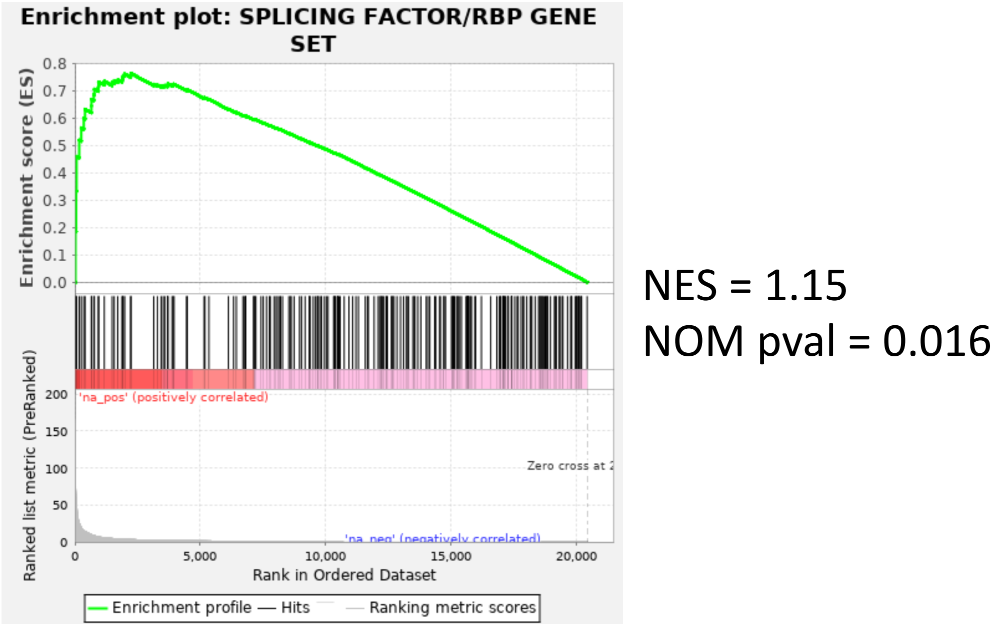

Enrichment plot: SPLICING FACTOR/RBP GENE SET
Enrichment score (ES)
0.8
0.7
0.6
0.5
0.4
0.3
0.2
0.1
0.0
Ranked list metric (PreRanked)
200
150
100
50
0
'na_pos' (positively correlated)
Zero cross at 2
'na_neg' (negatively correlated)
0
5,000
10,000
15,000
20,000
Rank in Ordered Dataset
Enrichment profile — Hits — Ranking metric scores
NES = 1.15
NOM pval = 0.016


b

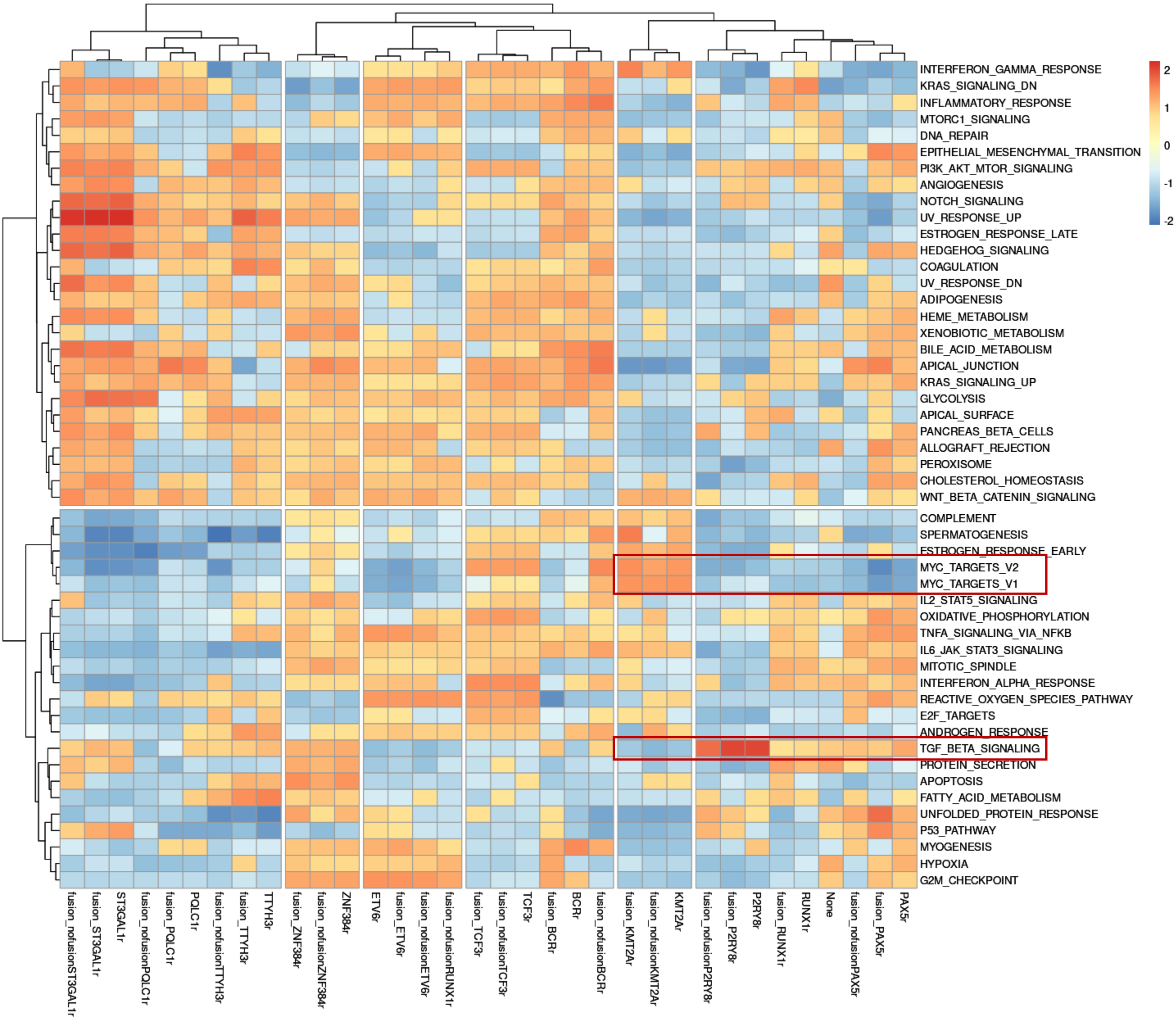

2
1
0
-1
-2
INTERFERON_GAMMA_RESPONSE
KRAS_SIGNALING_DN
INFLAMMATORY_RESPONSE
MTORC1_SIGNALING
DNA_REPAIR
EPITHELIAL_MESENCHYMAL_TRANSITION
PI3K_AKT_MTOR_SIGNALING
ANGIOGENESIS
NOTCH_SIGNALING
UV_RESPONSE_UP
ESTROGEN_RESPONSE_LATE
HEDGEHOG_SIGNALING
COAGULATION
UV_RESPONSE_DN
ADIPOGENESIS
HEME_METABOLISM
XENOBIOTIC_METABOLISM
BILE_ACID_METABOLISM
APICAL_JUNCTION
KRAS_SIGNALING_UP
GLYCOLYSIS
APICAL_SURFACE
PANCREAS_BETA_CELLS
ALLOGRAFT_REJECTION
PEROXISOME
CHOLESTEROL_HOMEOSTASIS
WNT_BETA_CATENIN_SIGNALING
COMPLEMENT
SPERMATOGENESIS
ESTROGEN_RESPONSE_EARLY
MYC_TARGETS_V2
MYC_TARGETS_V1
IL2_STAT5_SIGNALING
OXIDATIVE_PHOSPHORYLATION
TNFA_SIGNALING_VIA_NFKB
IL6_JAK_STAT3_SIGNALING
MITOTIC_SPINDLE
INTERFERON_ALPHA_RESPONSE
REACTIVE_OXYGEN_SPECIES_PATHWAY
E2F_TARGETS
ANDROGEN_RESPONSE
TGF_BETA_SIGNALING
PROTEIN_SECRETION
APOPTOSIS
FATTY_ACID_METABOLISM
UNFOLDED_PROTEIN_RESPONSE
P53_PATHWAY
MYOGENESIS
HYPOXIA
G2M_CHECKPOINT
fusion_nofusionST3GAL1r
fusion_ST3GAL1r
ST3GAL1r
fusion_nofusionPQLC1r
fusion_PQLC1r
PQLC1r
fusion_nofusionTTYH3r
fusion_TTYH3r
TTYH3r
fusion_ZNF384r
fusion_nofusionZNF384r
ZNF384r
ETV6r
fusion_ETV6r
fusion_nofusionETV6r
fusion_nofusionRUNX1r
fusion_TCF3r
fusion_nofusionTCF3r
TCF3r
fusion_BCRr
BCRr
fusion_nofusionBCRr
fusion_KMT2Ar
fusion_nofusionKMT2Ar
KMT2Ar
fusion_nofusionP2RY8r
fusion_P2RY8r
P2RY8r
fusion_RUNX1r
RUNX1r
None
fusion_nofusionPAX5r
fusion_PAX5r
PAX5r

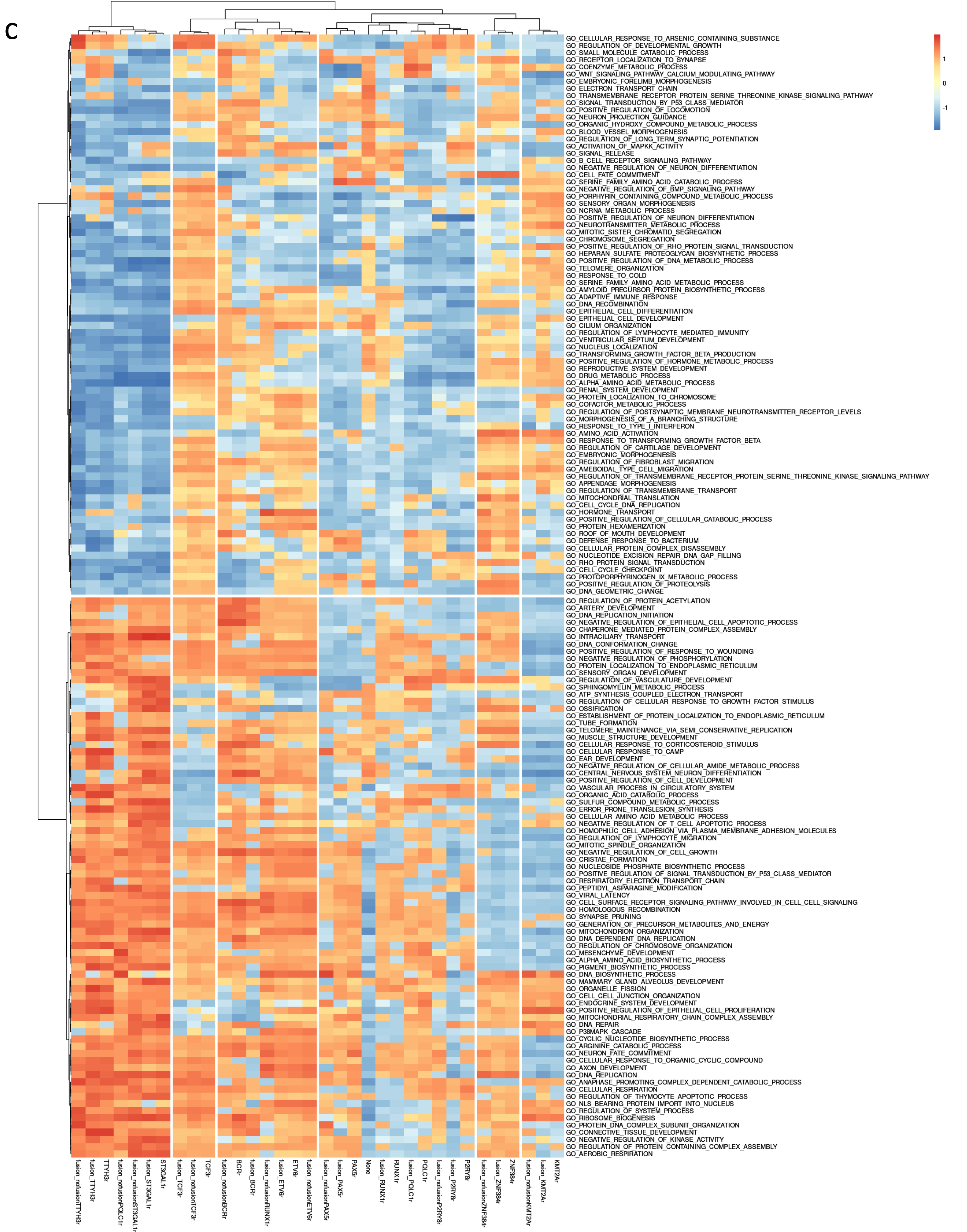


**Supplementary Figure 13. Gene set enrichment across fusion groups**. **(a)** Pre-ranked GSEA using a manually curated list of splicing factors and RBP. In this case, we compared the KMT2A-r versus the ETV6-r samples. In the other plots, we show the similarities between the differential expression (DE) patterns associated to each fusion group according to cancer hallmarks **(b)** and gene ontologies (Biological Process) **(c)**. For each DE analysis (x axis) we calculated the enriched hallmarks and ontologies. We used the top 10 up and down regulated in each comparison and clustered the groups using the normalized enriched score (NES), defined as log10(fold-change) x (-log10(corrected p-value)). The color indicates enrichment (NES > 0) or depletion (NES<0). *KMT2A-r* and *ETV6-r* show common hallmarks with opposite patterns. The Euclidean distances and Ward clustering method were used to generate this heatmaps.

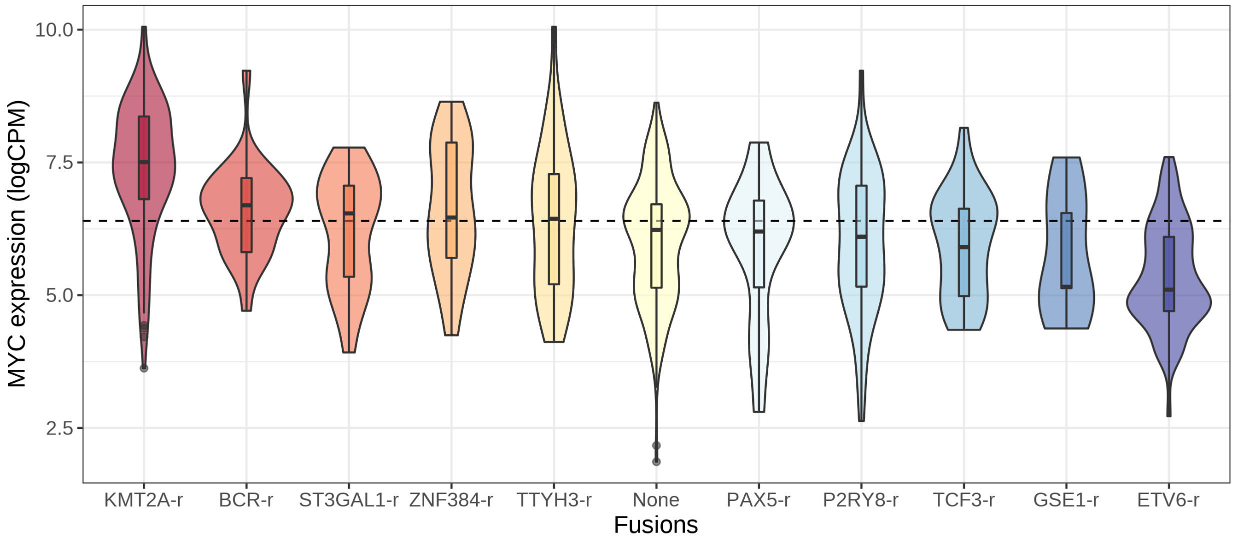


**Supplementary Figure 14.** Distribution of MYC expression values (y axis) for each fusion group compared to the mean expression in normal Feta-liver B-cells (horizontal dashed line). Expression is represented in log2(CPM), where CPM: counts per million.

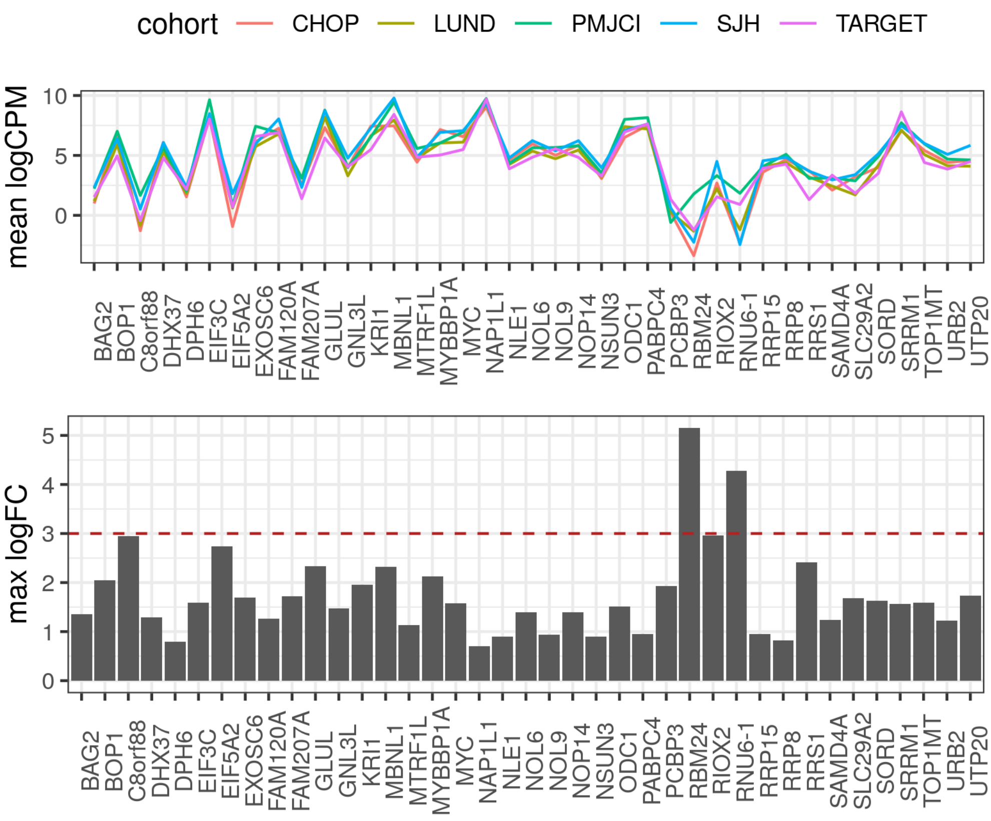


**Supplementary Figure 15. Inter cohort gene variation from 39 gene candidates.** Top panel show the mean logCPM expression for every gene by cohort. Bottom panel show the maximum logFC between the mean logCPM expression by cohort.

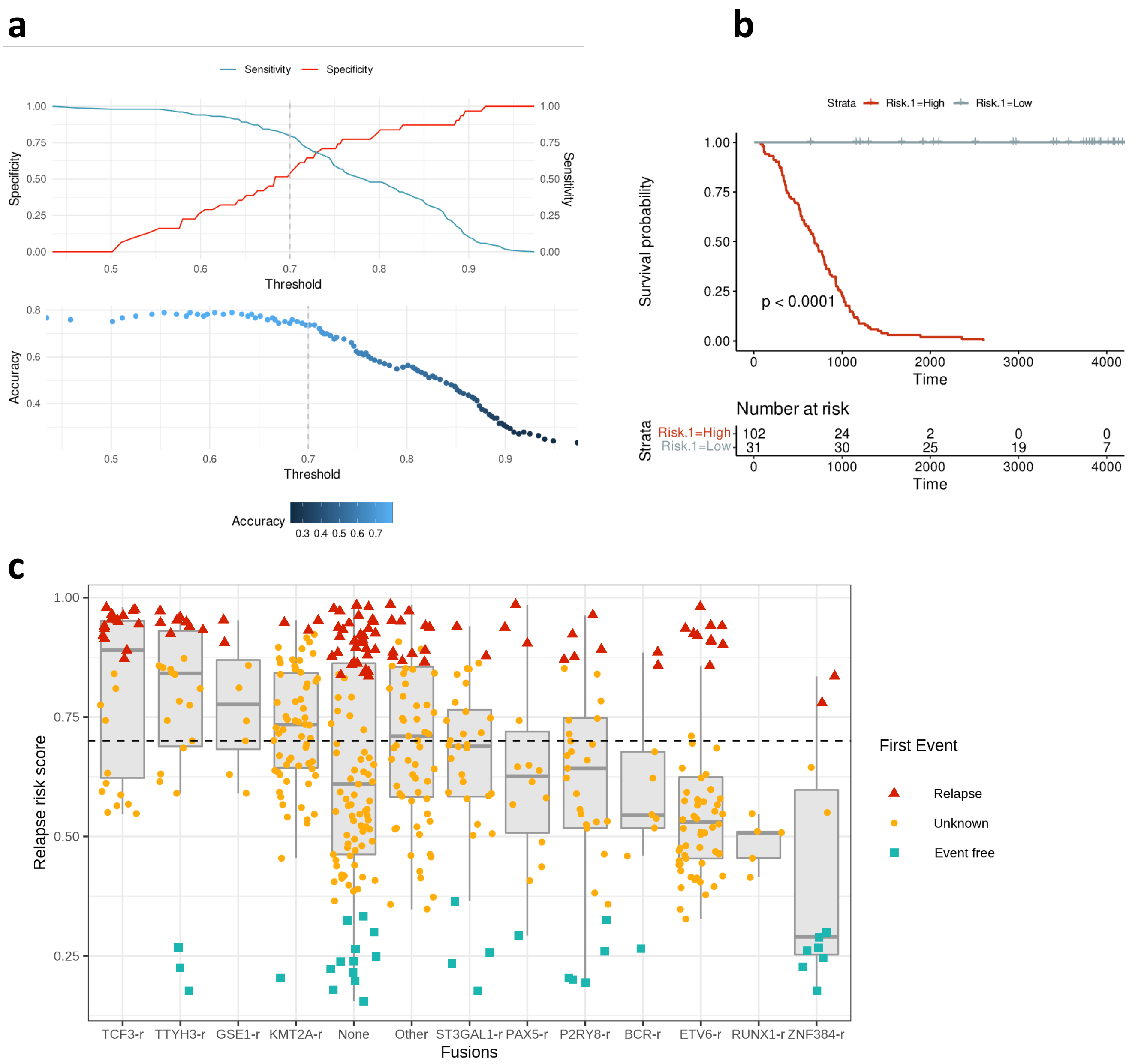


**Supplementary Figure 16. Gene expression signature associated to high risk. (a)** Specificity, Sensitivity, and Accuracy of the classification into low and high risk (y axis) as a function of the model score (x axis). We selected the threshold at score = 0.7, which shows a good balance between Specificity, Sensitivity, and Accuracy. Sensitivity is here calculated as the proportion of high-risk cases that are correctly predicted. Specificity is here calculated as the proportion of low-risk cases that are correctly predicted as low-risk. Accuracy is the proportion of correct predictions (high or low risk cases) over the total number of cases. **(b)** Kaplan-Meyer curve of the patients separated by the model, prior to performing leave-one-out benchmarking. **(c)** Box plots with the distribution of the K-score values for each of the fusion groups tested. We indicate the used threshold of 0.7 with a dashed line. Cases with relapse are indicated as triangles, cases with no relapse as squares, and cases without follow up annotation are indicated as circles.

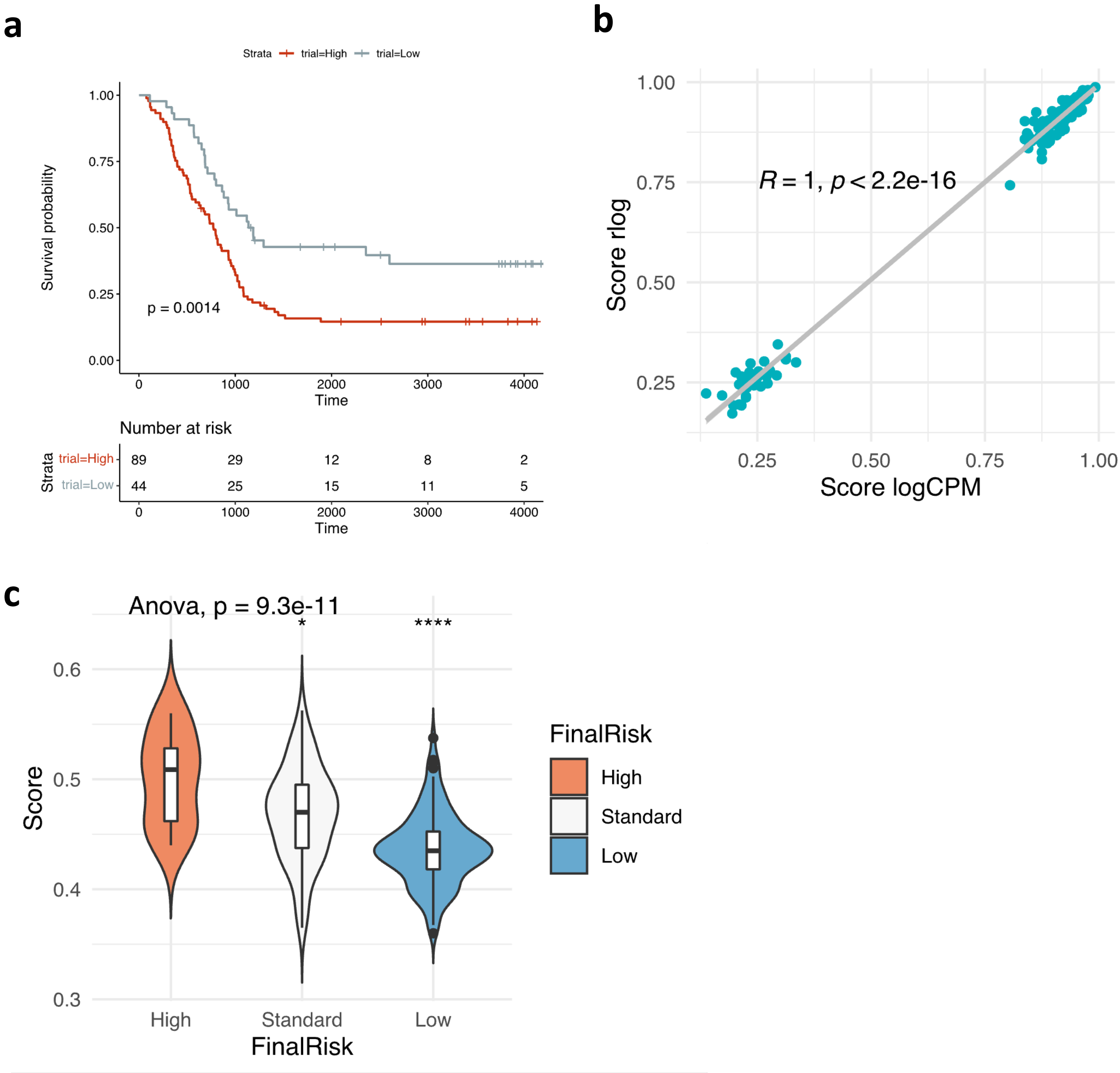


**Supplementary Figure 17. Signature validation on an independent cohort. a)** Kaplan-Meyer plot of the patients separated as high risk (red) (risk score >= 0.7) or low risk (grey) (risk score < 0.7) in a leave-one-out test with the rlog model. **b)** Pearson correlation between target scores using logCPM model and rlog model. **c)** Violin plots with patients scores from an independent cohort using rlog model and separated by the risk groups described on the paper. One-way Anova test for global mean differences. T-test to compare means by group regarding High-risk group (* indicates p <= 0.05, **** indicates p <= 0.00001).

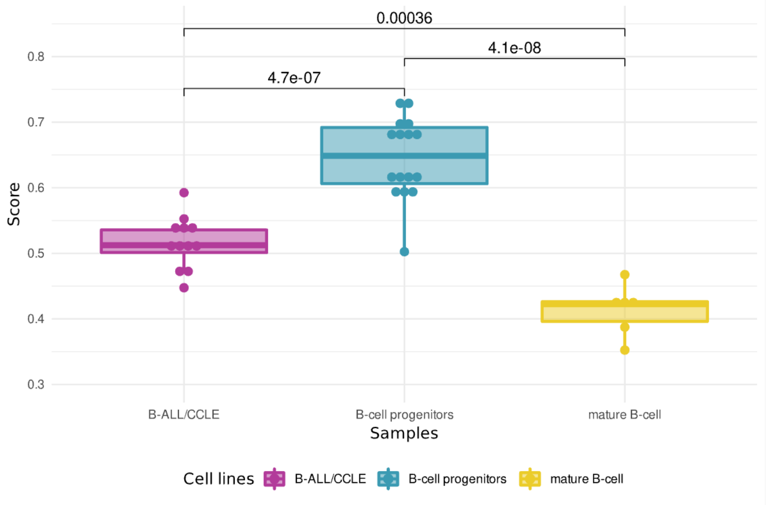


**Supplementary Figure 18. Predictor score boxplot by group of B-cell samples.** Score grouped by B-ALL cell lines from CCLE, B-cell progenitors from CHOP cohort and set of 6 cell lines from GM12878, p-value obtained from a t-test mean comparison.

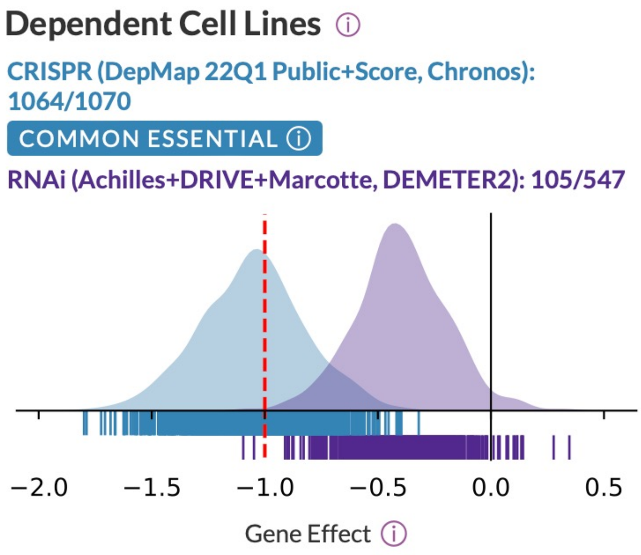


**Supplementary Figure 19. SRRM1 Dependent cell line plot from the depmap portal (https://depmap.org/portal/).** A cell line is considered dependent if it has a probability of dependency greater than 0.5. Gene effect is a score where a lower score means that a gene is more likely to be dependent on a given cell line. A score of 0 is equivalent to a gene that is not essential, whereas a score -1 corresponds to the median of all common essential genes.

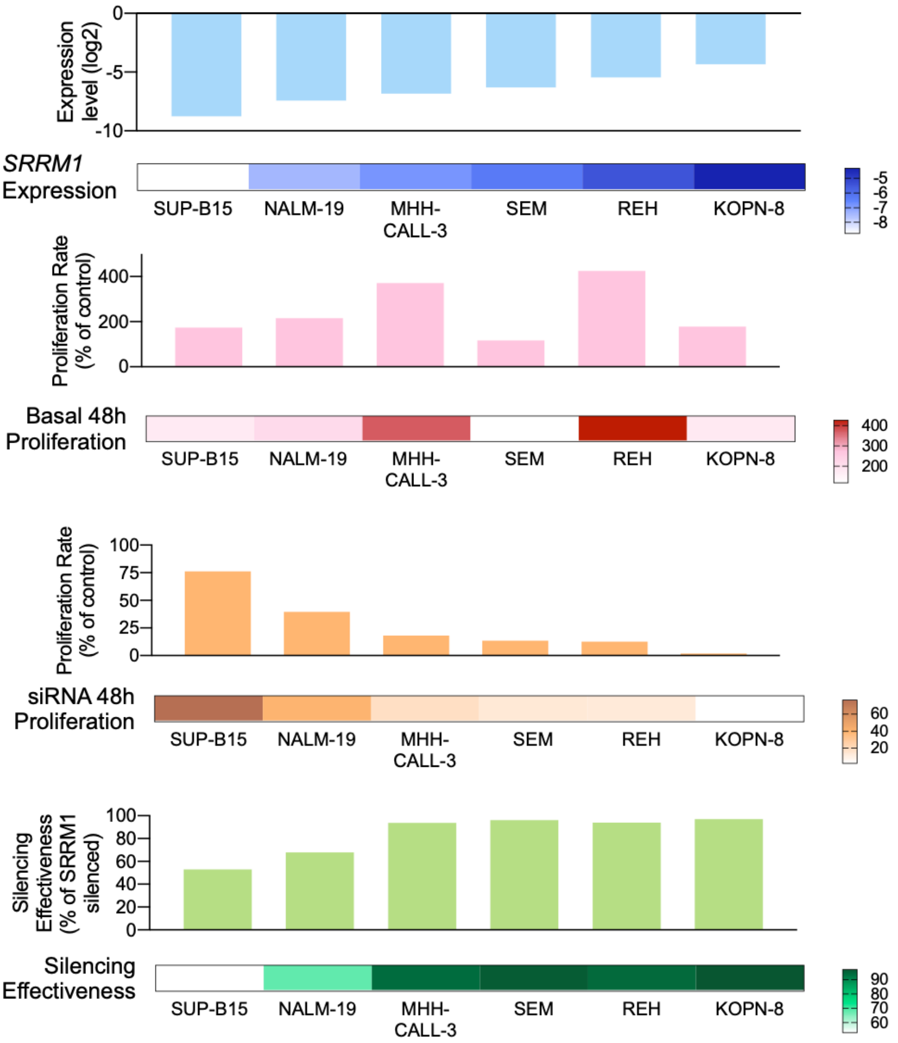


**Supplementary Figure 20.** Shows from top to bottom: comparison of SRRM1 mRNA levels, cell line proliferation in normal conditions, cell line proliferation under SRRM1 silencing and cell line silencing effectiveness across the different human leukemia cell lines used in the functional assays by qPCR (n=3).

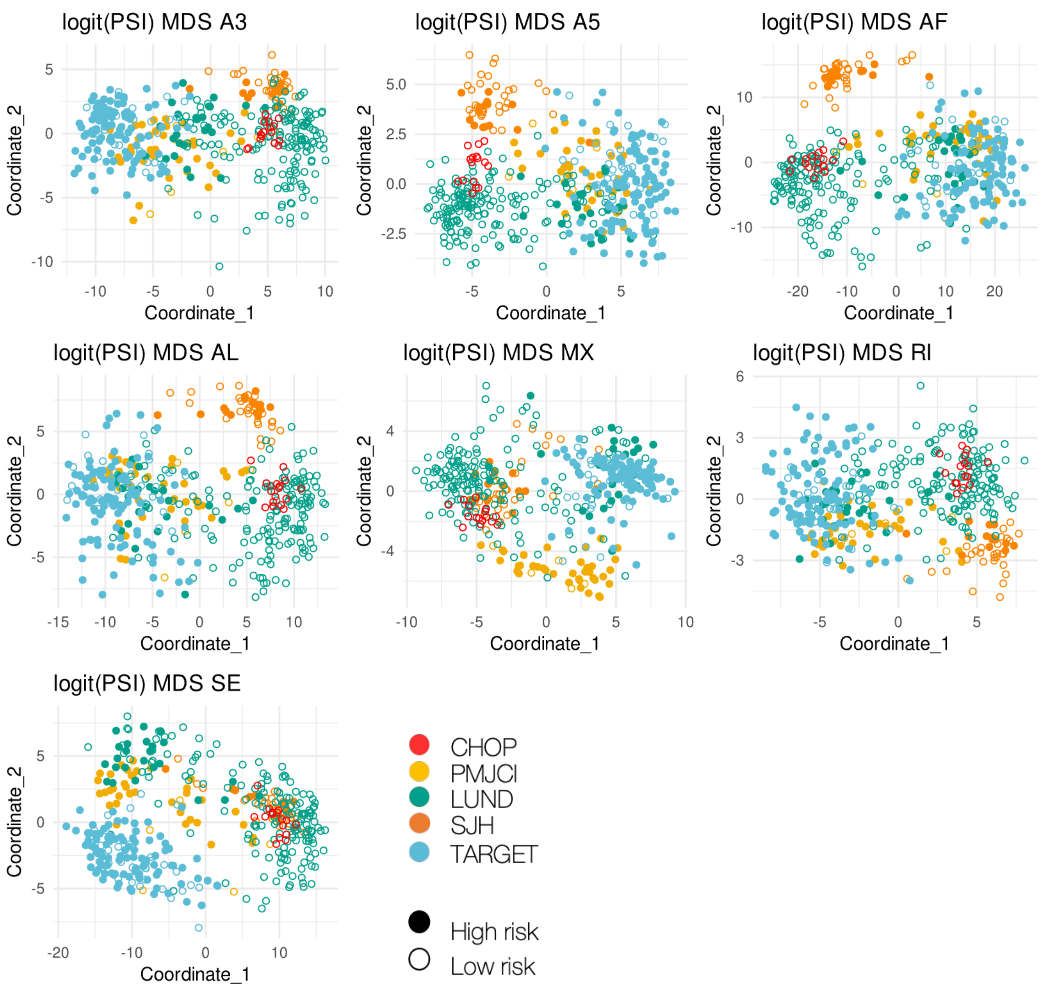


**Supplementary Figure 21. Splicing events associated with high risk.** The plot shows the multidimensional scale (MDS) analysis of the risk-associated events. Each dot represents a patient. The MDS was performed with the set of events associated with high risk, separated by the event type. Every color indicates the cohort of origin of the patient. A full dot is a patient predicted as high risk, whereas an empty dot is a patient predicted as low risk.

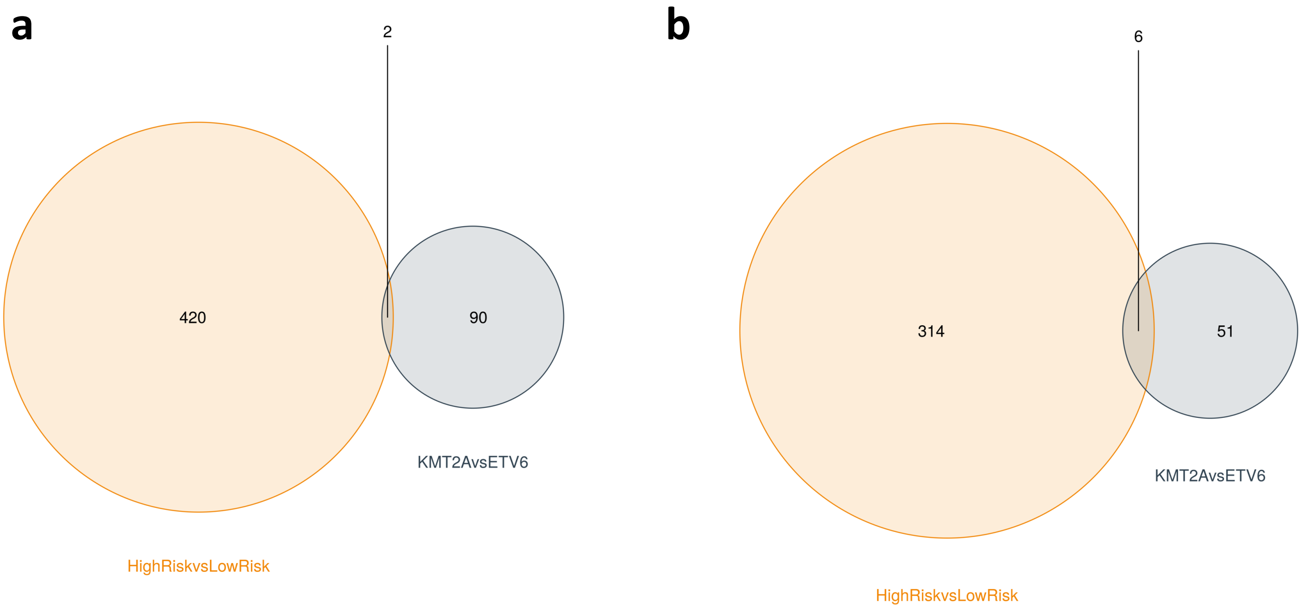


**Supplementary Figure 22. Vend Diagram comparison high risk vs low risk and KMT2A-r vs ETV6-r. (a)** Venn diagram overlapping differential included events comparing the High-risk vs low-risk patients from the predictor and comparing patients with KMT2A-r vs ETV6-r. **(b)** Venn diagram overlapping the genes affected by differential included events from the previous comparison on (a).

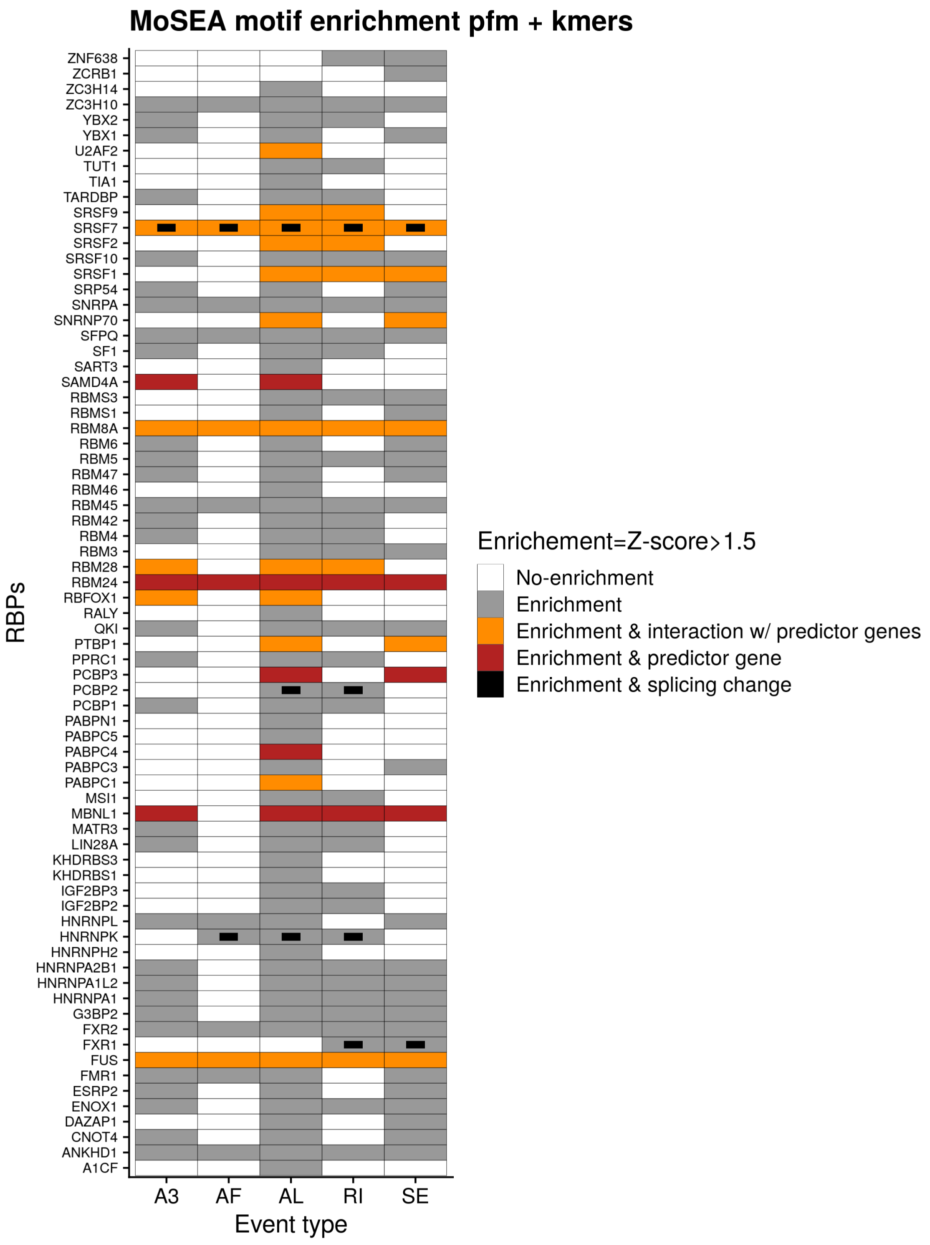


**Supplementary Figure 23. Motif enrichment.** Heatmap with the RBPs with enriched motifs in the events associated with high risk from the predictor. Motif enrichment was performed using MoSEA with k-mers and position frequency matrices (PFMs). MoSEA calculates a z-score to determine the association of the motif with the events that change inclusion between high and low risk, relative to the events that do not undergo any change. We indicate in white no enrichment, in grey when there is an enrichment, in orange when there is enrichment and the RBP interacts (via PPIs) with genes included in the predictor, and in red when there is motif enrichment and the RBP is part of the predictor. The black square inside indicates that there was a significant splicing change in the RBP gene.

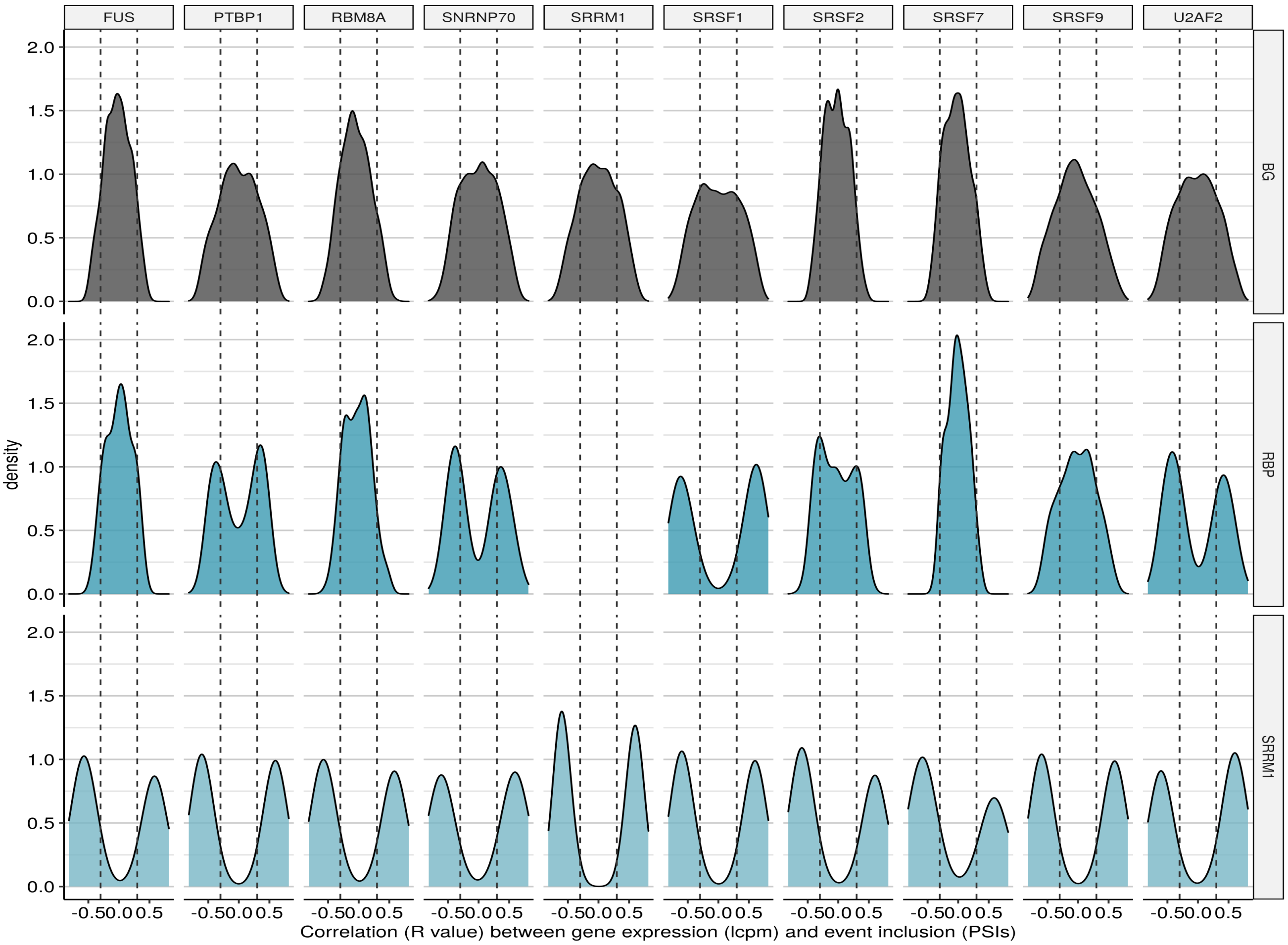


**Supplementary Figure 24. Correlation plot using RNA processing events with RBPs and SRRM1.** Distributions of the correlation values between splicing events and the expression of SRRM1 and the RBPs that interact with it along all the samples. The top row shows the distribution for background events randomly selected. The middle row show the events with a significant change between high and low risk and that also have a motif for the corresponding RBP. The bottom row show the same events from the middle row, but the distributions correspond to the correlation with SRRM1 expression.

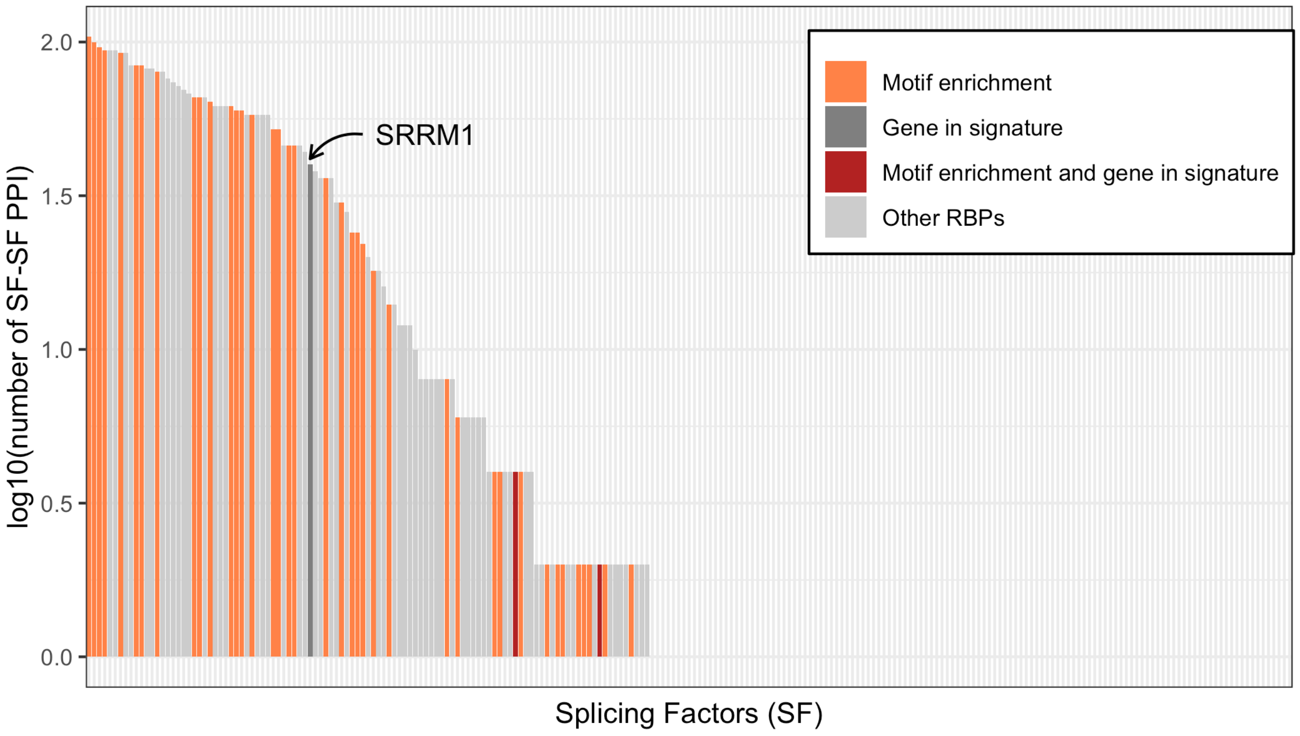


**Supplementary Figure 25. Total number of interaction by splicing factor.** Barplot distribution of the total number of interaction by every splicing factor. In orange are splicing factor with a significative motive enrichment. Dark grey, splicing factors that appear on the high-risk signature. Red, genes that appear on the signature and present a significative motive enrichment. Grey, other RBPs. We only show splicing factor with interactions obtained from STRING that pass the threshold described on methods.

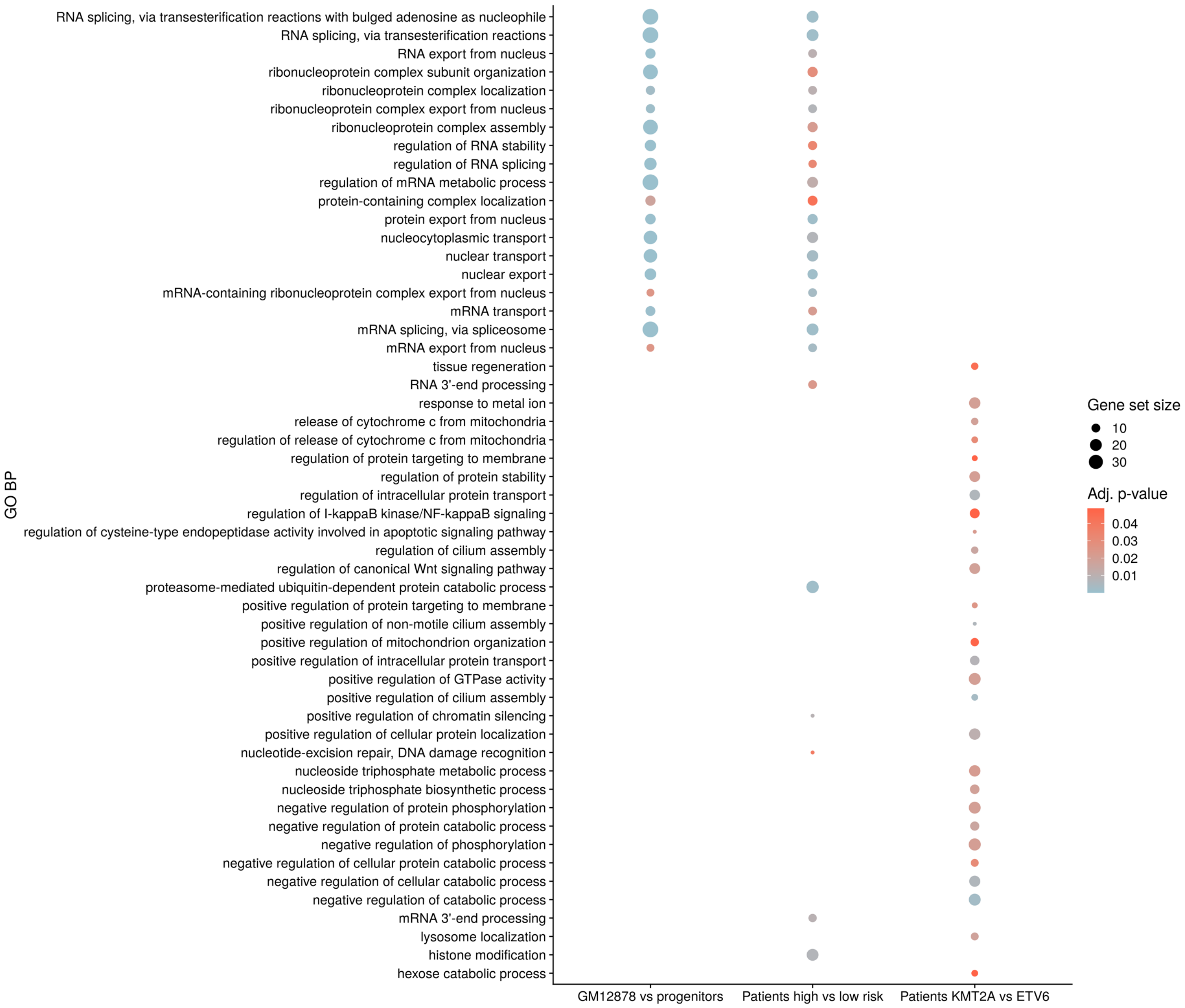


**Supplementary Figure 26. Gene set enrichment analysis with splicing events.** Pathway enrichment analysis using genes with splicing events that change significantly in any of 3 different comparisons: 1) GM12878 vs. B-cell progenitors, 2) high-risk vs. low-risk B-ALL patients, and 3) *KMT2A-r* vs *ETV6-r* patients.

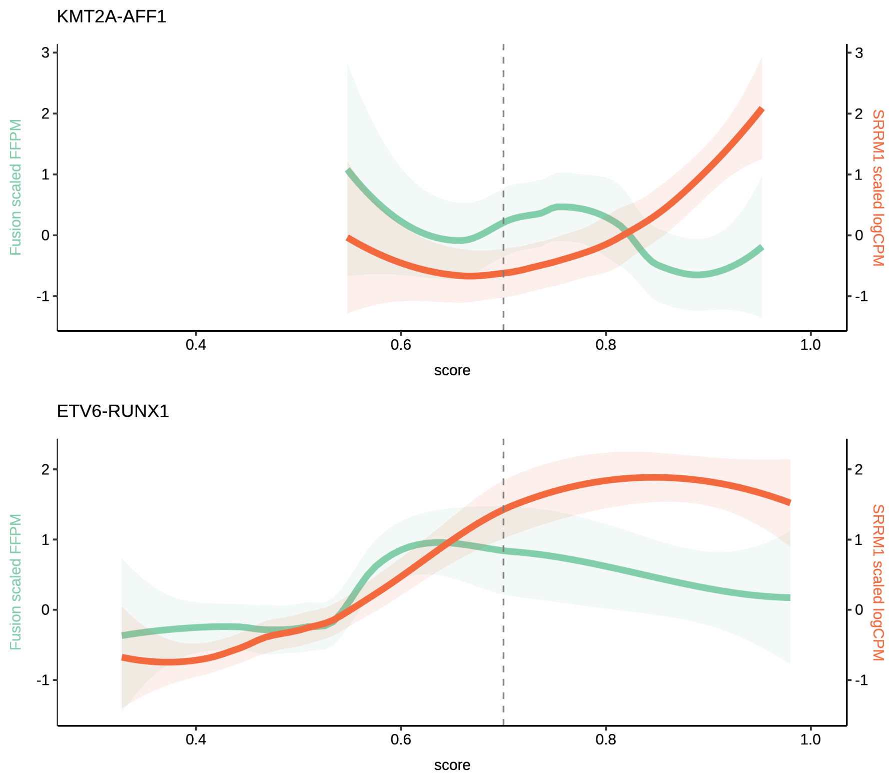


**Supplementary Figure 27. Score distribution in relation to fusion and SRRM1 expression.** For the KMT2A-AFF1 (upper panel) and the ETV6-RUNX1 (lower panel) we represent on the x axis our signature score, on the left Y axis the fusion expression scaled and centred at 0 FFPM, and on the right Y axis the scaled and centred at 0 log2CPM expression of SRRM1.

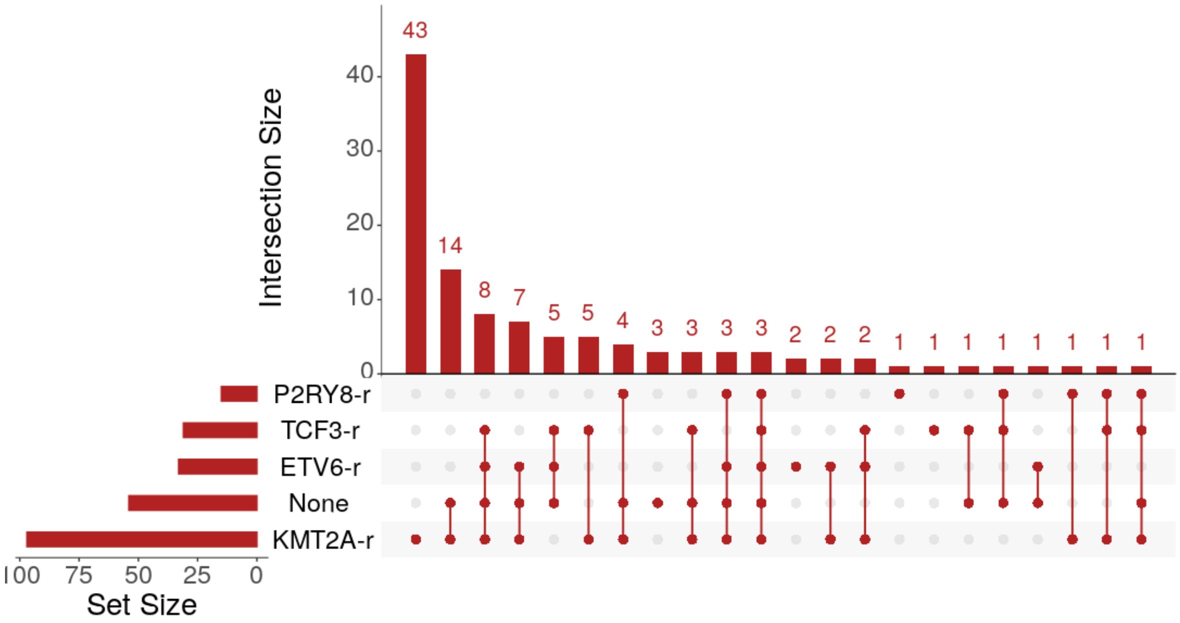


**Supplementary Figure 28. Events co-occurring in high-risk patients from different fusion backgrounds.** UpSet plot with the total number of events that co-occur with high risk associated with every fusion group. The bar plot indicates the total of number of events in each subset and the dots and lines the overlap of the events with every fusion group. Co-occurrence was measured as having average PSI = 0.5 or higher in two or more groups.

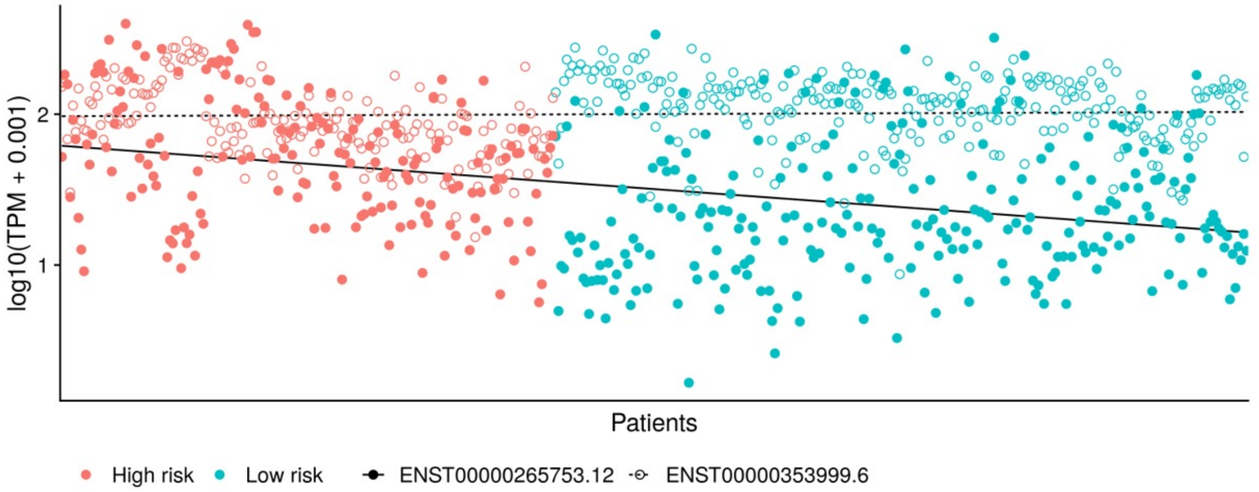


**Supplementary Figure 29. EIF4H major isoforms expression distribution**. Expression of the two most abundant EIF4H isoforms across patients separated by risk group. Isoform expression is given as log10(TPM+0.001) (y axis). The lines indicates the trend distribution for each isoform using smoothing with a linear model (method=*lm* in R).

a

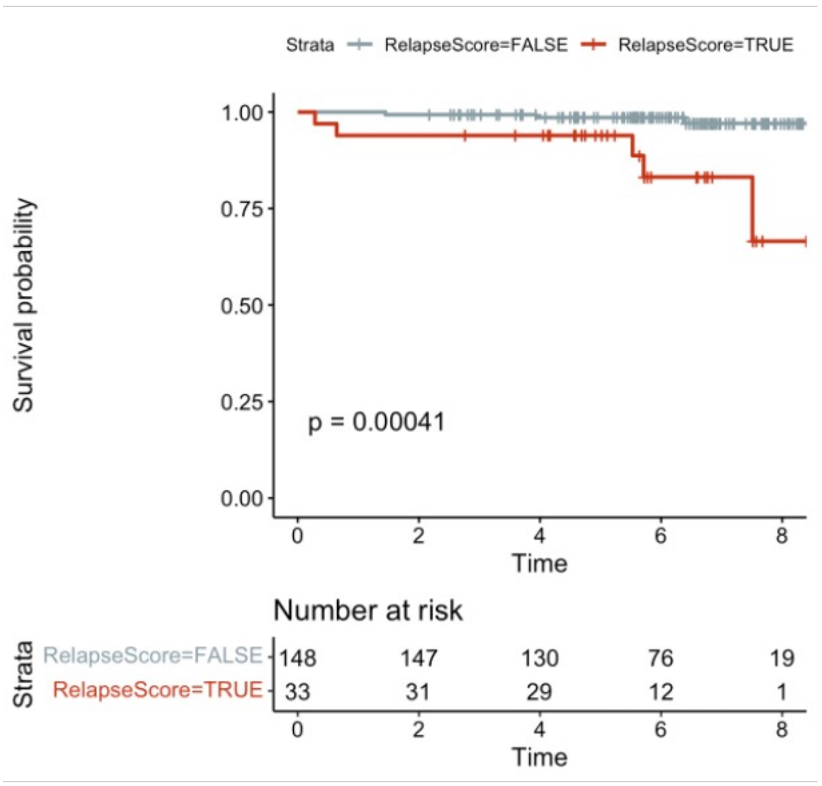


b

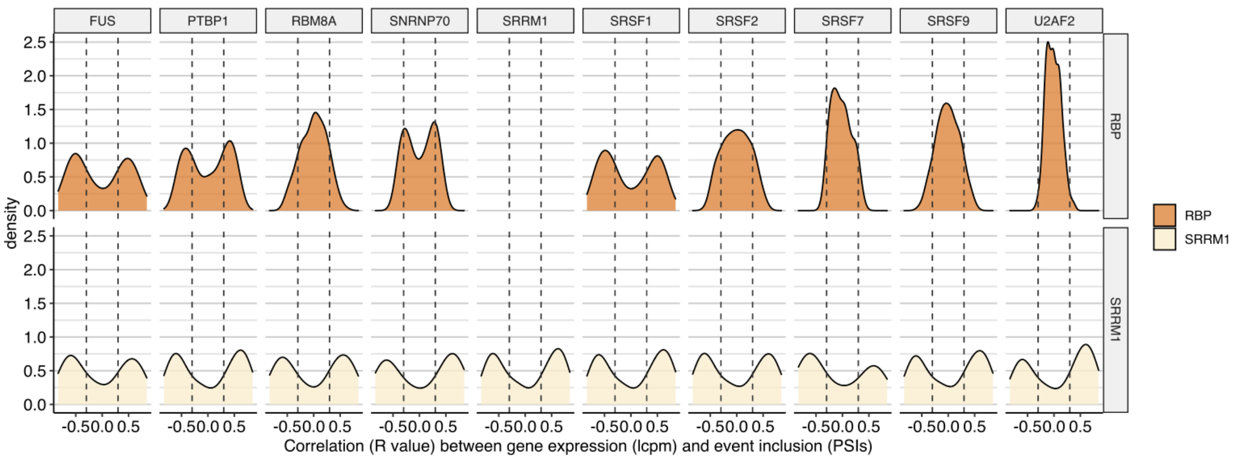


**Supplementary Figure 30. Validation of our risk score and RBP correlations in an independent cohort**. **(a)** Kaplan-Meyer analysis of the patients separated as high risk (red) (risk score >= 0.75) or low risk (grey) (risk score < 0.75). The p-value corresponds to a log-rank test (rlog p-value). **(b)** Distributions of the correlation values between splicing events and the expression of SRRM1 and the RBPs that interact with SRRM1 across the samples of the independent cohort. The events are those that were identified to have significant differential splicing between high and low-risk patients in the initial discovery cohorts and additionally have one or more motifs for the RBPs interacting with SRRM1. Out of those 422 events initially found in the discovery cohort, only 329 (78%) were also expressed in the validation cohort (had a defined PSI value calculated as described in Methods). The top panels show the correlation between those events' PSIs and the expression of each RBP, and the bottom panel shows the correlations between those events' PSIs and SRRM1 expression, both using the RNA-seq data from the new independent cohort.

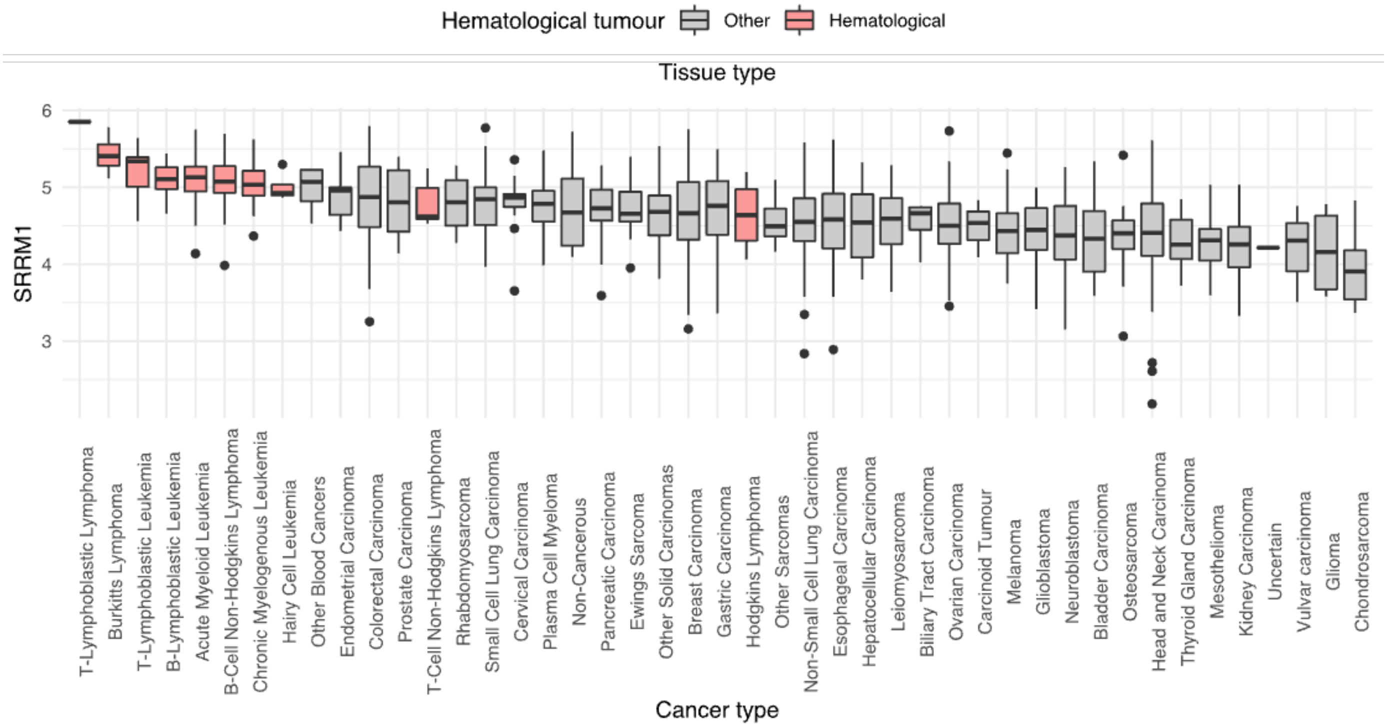


**Supplementary Figure 31. SRM1 protein level expression 946 human cell lines**. Protein expression of SRRM1 using the data from cancer cell lines grouped on the X axis by cancer type and SRRM1 expression level on Y axis. On red are coloured all the cancer types related with haematological tumours.

| Project Name | Project Database ID | Sequencing platform | Read length | Seq. Read types |
|---|---|---|---|---|
| SJH | EGAS00001000246 | Illumina HiSeq 2000 | 100 | paired-end |
| LUND | EGAS00001001795 | Illumina HiScanSQ | 100 | paired-end |
| CHOP | GSE115656 | Illumina HiSeq 2500 | 100 | paired-end |
| TARGET | phs000463 (ALL phase1) / phs000464 (ALL phase2) | Illumina HiSeq 2000 | 100 | paired-end |
| PMJCI | N/A | Illumina HiSeq 2500 | 76 | paired-end |

**Supplementary Table 1.** Summary table with the sequencing platform details from the cohorts used on this study.

| Variable | Stats / Values | Freqs (% of Valid) | Missing(%) |
|---|---|---|---|
| Project | CHOP | 18 (3.5%) | 0 (0.0%) |
| | LUND | 193 (37.8%) | |
| | PMJCI | 50 (9.8%) | |
| | SJH | 58 (11.4%) | |
| | TARGET_phase1 | 12 (2.4%) | |
| | TARGET_phase2 | 179 (35.1%) | |
| Type | ALL | 510 (100%) | 0 (0.0%) |
| Tissue | Blood | 102 (20%) | 0 (0.0%) |
| | Bone Marrow (BM) | 408 (80%) | |
| Blasts | Mean (sd): 93.4 (6.8); IQR (CV): 8 (0.1);<br>min < med < max: 43 < 95 < 100 | — | 404 (79.2%) |
| Time sample extraction | DX (diagnosis) | 428 (83.9%) | 0 (0.0%) |
| | RL (relapse) | 82 (16.1%) | |
| Gender | Female (F) | 240 (47.1%) | 0 (0.0%) |
| | Male (M) | 270 (52.9%) | |
| Age (month) | Mean (sd): 67.9 (62.8); IQR (CV): 96 (0.9);<br>min < med < max: 0 < 48 < 364.9 | — | 2 (0.4%) |
| Fusions | None | 239 (46.9%) | 0 (0.0%) |
| | KMT2A-AFF1 | 59 (11.6%) | |
| | ETV6-RUNX1 | 56 (11%) | |
| | TCF3-PBX1 | 31 (6.1%) | |
| | KMT2A-MLLT3 | 18 (3.5%) | |
| | KMT2A-MLLT1 | 15 (2.9%) | |
| | P2RY8-CRLF2 | 9 (1.8%) | |
| | IGH-DUX4 | 6 (1.2%) | |
| | BCR-ABL1 | 4 (0.8%) | |
| | [ 57 others ] | 73 (14.3%) | |
| Cell of Origin | B Cell ALL | 320 (62.7%) | 0 (0.0%) |
| | B-Precursor | 190 (37.3%) | |
| First event | None | 31 (16.2%) | 319 (62.5%) |
| | Relapse | 160 (83.8%) | |
| Event Free Survival Time (days) | Mean (sd): 1144.8 (1046.1); IQR (CV): 683.5 (0.9);<br>min < med < max: 77 < 851 < 4383 | — | 319 (62.5%) |
| Vital Status | Alive | 80 (41.9%) | 319 (62.5%) |
| | Dead | 111 (58.1%) | |
| Overall Survival Time (days) | Mean (sd): 1810.7 (1230.1); IQR (CV): 2122 (0.7);<br>min < med < max: 187 < 1290 < 4383 | — | 319 (62.5%) |

**Supplementary Table 2.** Summary with the clinical information for the samples selected from the multiple cohort study used on the analysis. For the rows Blast, Age (month), Event Free Survival Time (days) and Overall Survival Time (days), we calculated a Mean, a standard deviation (sd), the interquartile range (IQR), coefficient of variation (cv), minimum value (min), median (med) and maximum value (max).

| Gene name | beta | HR (95% CI for HR) | Wald-test | P-value | LRT-test | LRT-pvalue |
|---|---|---|---|---|---|---|
| BAG2 | 0.75 | 2.1 (1.3-3.4) | 11 | 0.012 | 9.7 | 0.021 |
| BOP1 | 1 | 2.9 (1.7-4.7) | 18 | 0.00039 | 15 | 0.0015 |
| C8orf88 | 0.51 | 1.7 (1.1-2.5) | 8 | 0.045 | 8 | 0.045 |
| DHX33 | 0.35 | 1.4 (0.95-2.1) | 4.9 | 0.18 | 4.9 | 0.18 |
| DHX37 | 0.6 | 1.8 (1.2-2.7) | 11 | 0.013 | 10 | 0.015 |
| DPH6 | 0.54 | 1.7 (1.1-2.6) | 8.2 | 0.043 | 7.7 | 0.053 |
| EIF2AK4 | 0.47 | 1.6 (0.76-3.4) | 3.5 | 0.32 | 3.7 | 0.3 |
| EIF3C | 0.8 | 2.2 (1.4-3.4) | 15 | 0.0016 | 14 | 0.0029 |
| EIF5 | 0.47 | 1.6 (0.88-2.9) | 4.3 | 0.23 | 4.6 | 0.2 |
| EIF5A2 | 0.88 | 2.4 (1.5-3.9) | 14 | 0.0031 | 13 | 0.0058 |
| EXOSC6 | 0.8 | 2.2 (1.5-3.3) | 17 | 0.00071 | 17 | 0.00086 |
| FAM120A | 0.79 | 2.2 (1.3-3.8) | 9.9 | 0.02 | 9 | 0.029 |
| FAM207A | 0.9 | 2.5 (1.6-3.9) | 18 | 5.00E-04 | 15 | 0.0015 |
| GLUL | 0.75 | 2.1 (1.4-3.3) | 14 | 0.0034 | 13 | 0.0052 |
| GNL3L | 0.99 | 2.7 (1.3-5.6) | 9.2 | 0.027 | 11 | 0.012 |
| IGF2BP2 | 0.47 | 1.6 (1.1-2.4) | 7.2 | 0.066 | 7.3 | 0.063 |
| KRI1 | 0.76 | 2.1 (1.2-3.8) | 9.2 | 0.026 | 8 | 0.047 |
| MBNL1 | 0.75 | 2.1 (1.1-4) | 8 | 0.045 | 6.7 | 0.081 |
| MTRF1L | 1 | 2.8 (1.5-5.3) | 12 | 0.0083 | 15 | 0.0022 |
| MYBBP1A | 0.78 | 2.2 (1.3-3.7) | 9.7 | 0.022 | 8.6 | 0.035 |
| MYC | 1.2 | 3.2 (1.8-5.6) | 19 | 0.00029 | 15 | 0.0015 |
| NAF1 | -0.4 | 0.67 (0.35-1.3) | 3.5 | 0.32 | 3.4 | 0.34 |
| NAP1L1 | 0.62 | 1.9 (1.2-2.8) | 11 | 0.01 | 11 | 0.011 |
| NLE1 | 1.3 | 3.7 (2.3-5.9) | 29 | 2.00E-06 | 25 | 1.80E-05 |
| NOL6 | 0.91 | 2.5 (1.4-4.5) | 11 | 0.012 | 9.8 | 0.02 |
| NOL9 | 0.79 | 2.2 (1.2-3.9) | 9.2 | 0.026 | 11 | 0.014 |
| NOM1 | 0.5 | 1.6 (1-2.6) | 6.2 | 0.1 | 5.8 | 0.12 |
| NOP14 | 0.78 | 2.2 (1.3-3.8) | 9.6 | 0.023 | 8.4 | 0.039 |

| Gene name | beta | HR (95% CI for HR) | Wald-test | P-value | LRT-test | LRT-pvalue |
|---|---|---|---|---|---|---|
| NRIP2 | 0.32 | 1.4 (0.71-2.7) | 2.8 | 0.42 | 2.9 | 0.4 |
| NSUN3 | 0.9 | 2.5 (1.6-3.7) | 21 | 0.00012 | 21 | 0.00013 |
| NSUN4 | 0.45 | 1.6 (0.99-2.5) | 5.5 | 0.14 | 5.9 | 0.12 |
| ODC1 | 1 | 2.7 (1.8-4.1) | 26 | 1.10E-05 | 26 | 1.20E-05 |
| PABPC1 | 0.54 | 1.7 (1-2.9) | 6.2 | 0.1 | 5.7 | 0.13 |
| PABPC4 | 0.71 | 2 (1.3-3.1) | 13 | 0.004 | 13 | 0.0052 |
| PCBP3 | 0.95 | 2.6 (1.3-5) | 9.9 | 0.019 | 12 | 0.0085 |
| PPP1R15B | -0.25 | 0.78 (0.39-1.5) | 2.5 | 0.48 | 2.5 | 0.47 |
| PPRC1 | 0.63 | 1.9 (1.1-3.3) | 6.9 | 0.076 | 6.2 | 0.1 |
| RBM15 | 0.43 | 1.5 (0.98-2.4) | 5.5 | 0.14 | 5.8 | 0.12 |
| RBM24 | 0.72 | 2.1 (1.3-3.2) | 12 | 0.0069 | 12 | 0.0069 |
| RIOX2 | 0.91 | 2.5 (1.2-5.2) | 8 | 0.047 | 9.6 | 0.022 |
| RNU2_1 | 0.86 | 2.4 (1-5.4) | 6.3 | 0.099 | 7.3 | 0.063 |
| RNU6_1 | 0.7 | 2 (1.2-3.3) | 9.4 | 0.024 | 8.9 | 0.03 |
| RRP15 | 0.82 | 2.3 (1.4-3.7) | 13 | 0.0042 | 12 | 0.0085 |
| RRP8 | 0.69 | 2 (1.3-3) | 12 | 0.007 | 13 | 0.0054 |
| RRS1 | 1.4 | 4.2 (2.4-7.2) | 28 | 3.80E-06 | 22 | 8.00E-05 |
| SAMD4A | 0.74 | 2.1 (1.4-3.2) | 14 | 0.0024 | 14 | 0.0033 |
| SLC29A2 | 0.71 | 2 (1.4-3) | 14 | 0.0031 | 13 | 0.0043 |
| SORD | 0.86 | 2.4 (1.4-3.9) | 14 | 0.0035 | 12 | 0.0078 |
| SRRM1 | 0.54 | 1.7 (1.2-2.6) | 9.2 | 0.027 | 8.9 | 0.03 |
| TCOF1 | 0.64 | 1.9 (1.1-3.2) | 7.7 | 0.052 | 8.4 | 0.039 |
| TOP1MT | 1.1 | 3.1 (1.9-5.1) | 23 | 5.00E-05 | 19 | 0.00034 |
| UPF1 | 0.67 | 2 (0.9-4.2) | 4.7 | 0.19 | 5.5 | 0.14 |
| URB1 | -0.43 | 0.65 (0.34-1.3) | 3.6 | 0.31 | 3.8 | 0.29 |
| URB2 | 0.69 | 2 (1.3-3.2) | 11 | 0.014 | 12 | 0.0089 |
| UTP20 | -0.79 | 0.45 (0.24-0.84) | 8 | 0.045 | 7.2 | 0.065 |

**Supplementary Table 3.** Summary table with the Cox proportional hazards regression results per gene adjusted by age and gender. Beta is the regression coefficient, a positive sign means that the hazard (risk of relapse) is higher, and thus the prognosis worse, for subjects with higher values of that variable. HR(95% CI for HR) is the hazard ratio with upper and lower 95% confidence interval. Log Ratio test (LRT) was added to compliment Wald test.
